# Species-specific traits mediate avian demographic responses under past climate change

**DOI:** 10.1101/2022.08.16.504093

**Authors:** Ryan R Germain, Shaohong Feng, Guangji Chen, Gary R. Graves, Joseph A. Tobias, Carsten Rahbek, Fumin Lei, Jon Fjeldså, Peter A. Hosner, M. Thomas P. Gilbert, Guojie Zhang, David Nogués-Bravo

## Abstract

Anticipating species’ responses to environmental change is a pressing mission in biodiversity conservation. Despite decades of research investigating how climate change may affect population sizes, historical context is lacking and the traits which mediate demographic sensitivity to changing climate remain elusive. We use whole-genome sequence data to reconstruct the demographic histories of 263 bird species over the past million years and identify networks of interacting morphological and life-history traits associated with changes in effective population size (*N_e_*) in response to climate warming and cooling. Our results identify direct and indirect effects of key traits representing dispersal, reproduction, and survival on long-term demographic responses to climate change, thereby highlighting traits most likely to influence population responses to on-going climate warming.

**One-Sentence Summary:** Interacting traits influence sensitivity of bird population sizes to climate warming and cooling over the past million years.

---

Human-induced changes to the global environment are affecting biodiversity at an unprecedented rate, with animal populations having declined drastically since 1970^1–4^. Despite efforts to quantify contemporary population trends and disentangle the effects of different drivers of global change, we currently lack historical context as to whether similar declines have occurred before, and if species-specific traits influence long-term demographic sensitivity to environmental challenges such as climate change^5–7^. In particular, identifying common demographic patterns over evolutionary timescales, and before the Anthropocene epoch, can reveal how life-history strategies influence population dynamics during periods of wide-spread climatic stress^8–10^. Elucidating these strategies will aid conservation efforts through identifying species with characteristics prone to demographic decline under current and future challenges.

Climate change is regarded as a key environmental regulator of demographic change. It is hypothesized to affect demography via its effects on reproduction, survival/growth, and dispersal^11,12^. However, responses to climate change can vary dramatically among even closely-related or co-occurring species because of differing life-history traits and strategies^13–15^. Theoretical predictions and empirical evidence suggest that larger-bodied, slower-reproducing species with limited dispersal capacity are more sensitive to sustained climate change (table S11) because of reduced adaptive potential and/or limited ability to exploit climate refugia. However, evaluation of the role of traits on demographic sensitivity to climate change is typically tested only with contemporary data over shorter ecological timescales. Because additional environmental stressors such as land-use change and over-exploitation may mask or confound demographic responses^9,16–18^, the ability to identify relationships between life-history traits and demographic sensitivity to climate change remains constrained when limited to contemporary data.

Periods of climate warming and cooling over the Earth’s history offer a unique opportunity to quantify effects of species-specific traits on demographic sensitivity to climate change in the absence of confounding anthropogenic stressors^5,6^. We use whole-genome sequence data^19^, and Pairwise Sequential Markovian Coalescent (PSMC) analysis^20^ to reconstruct the long-term (~ one million years) demographic histories of 263 bird species, representing 39 orders distributed from the poles to the tropics. We then quantify demographic responses to the most recent warming and cooling periods before widespread human activity, and identify network effects of morphological and life-history traits related to survival/growth, reproduction, and dispersal which influence overall demographic sensitivity to climate change.

## Results and Discussion

Effective population sizes (*N_e_*) varied substantially across avian species and over time and space (table S1, Supplementary Text S1). Demographic clustering revealed seven main demographic patterns, inferred from temporal patterns of *N_e_* over the past million years (Fig. 1A, table S3). Overall position of each species in the avian phylogeny did not explain the observed differences among demographic patterns (fig. S7). Passerines (order Passeriformes, which represent more than half of all extant bird species globally^21^; here n = 123 species) and Non-Passerines (n = 140 species) were unequally distributed among the demographic clusters (*χ* = 23.81, df = 6, p < 0.001), where Passerines were most heavily represented in clusters 1, 3, 4, and 7 (i.e., demographic peaks in the more recent upper/middle Pleistocene; Fig 1A; Table S3). In contrast, Non-Passerines were most represented in clusters 5, 6, and 7, depicting demographic peaks during the more ancient periods of the middle/lower Pleistocene (Fig 1A, Table S3). These results were further reflected in lower mean *N_e_* values for Passerines in the more distant past, despite Passerines exhibiting consistently higher mean *N_e_* than Non-Passerines over the past million years (Fig. 1B). Species currently classified as “threatened” or “near-threatened” (IUCN Red List; here n = 34 species) were evenly distributed among the seven main demographic patterns (*χ*^2^ = 5.77, df = 6, p = 0.45) and exhibited varying demographic trends over time (Fig. 1C), indicating that current conservation status is unlikely to be the result of long-term demographic variation.

**Fig. 1.**
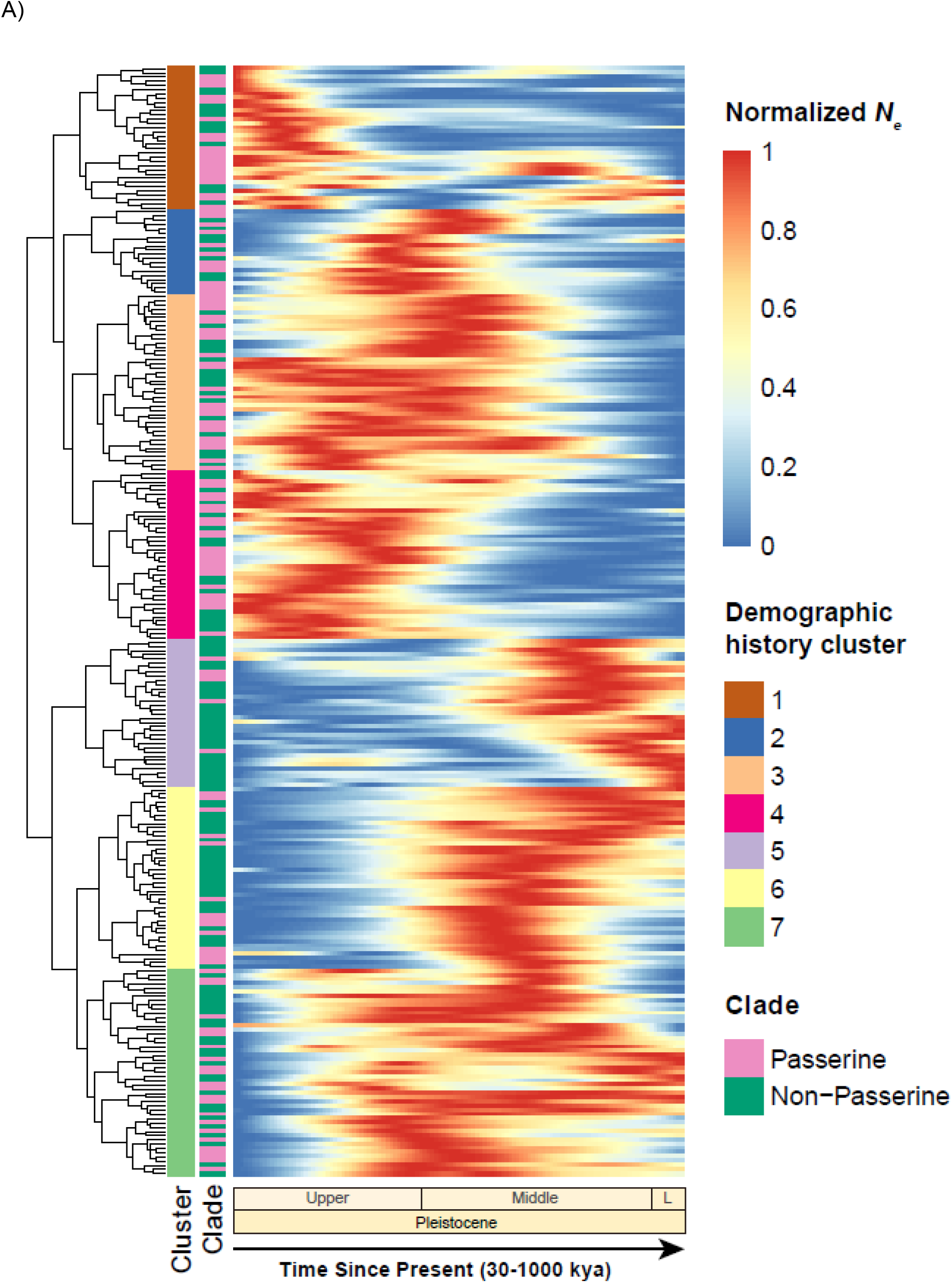
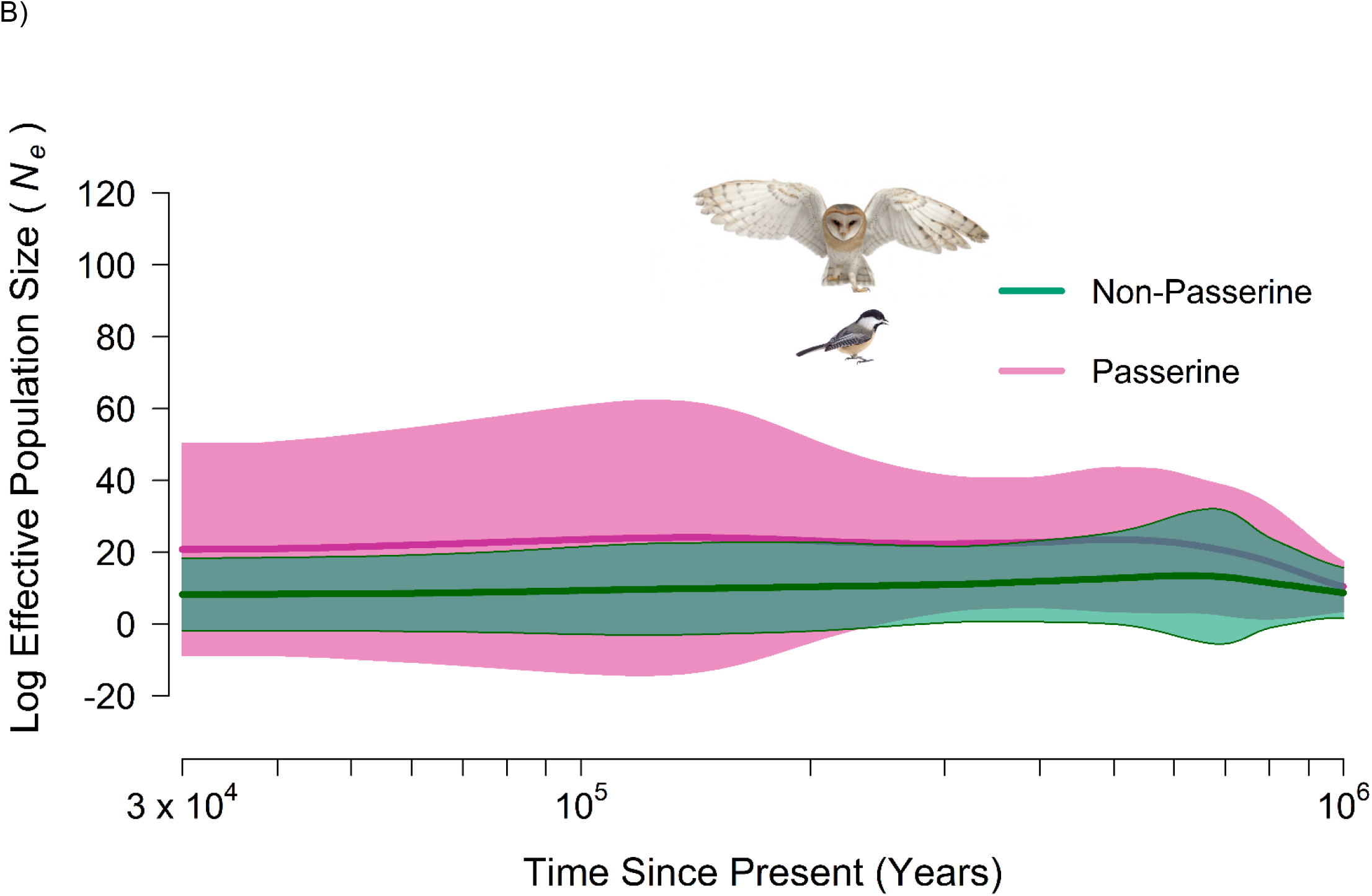
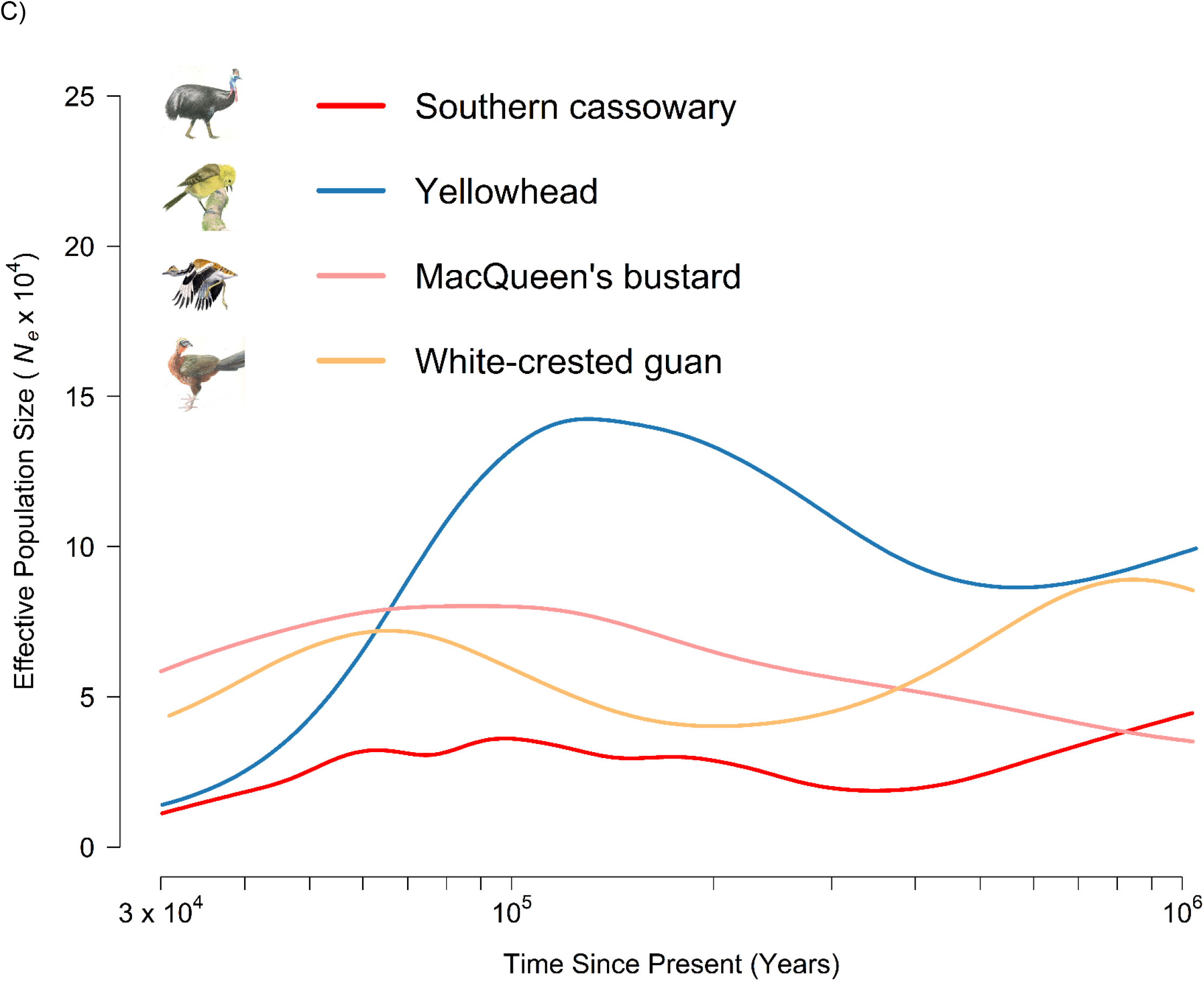
Demographic histories of 263 avian species from 30,000 to 1 million years ago (all x-axes presented on the log_10_ scale). A) Cluster analysis of normalized *N_e_* values reveal seven main demographic patterns over the Upper/Middle/Lower(L) Pleistocene. Clusters designating Groups 1–7 (see table S1 for species included in each group) were based on overall similarity of long-term *N_e_* patterns, but can be distinguished by when the majority of species reached their relative peak (see table S3). B) Mean log effective population size (*N_e_*) values (±SD) of Passerine and Non-Passerine species. Passerines exhibited consistently higher *N_e_* values (mean sample difference = 9.21) across 120 equally-spaced (loglinear) time points from 30–1000kya (paired t-test, t_119_ = 27.9, p <0.0001). C) Examples of differing demographic histories of species currently designated as “threatened” under IUCN Red List status, where species arriving at similar *N_e_* values at ~30kya follow differing demographic patterns over time (illustrations by JF).

Globally, mean normalized *N_e_* for all 263 species increased from 1mya to ~500-600kya, after which it steadily declined (Fig 2, ‘Global Average’). Across the Earth’s major zoogeographic realms (Supplementary Text S1), species differed only slightly in when they reached their mean demographic peaks. Most species followed similar patterns of higher mean *N_e_* in more ancient periods and lower *N_e_* values closer to 30kya (Fig 2), and there was little concordance between geographic realm and allocation to a given demographic cluster group (fig. S21). Overall, we detected minimal variation in demographic trends among realms (fig. S22), indicating little effect of geographic variation on overall patterns of demographic change over time.

**Fig. 2.**
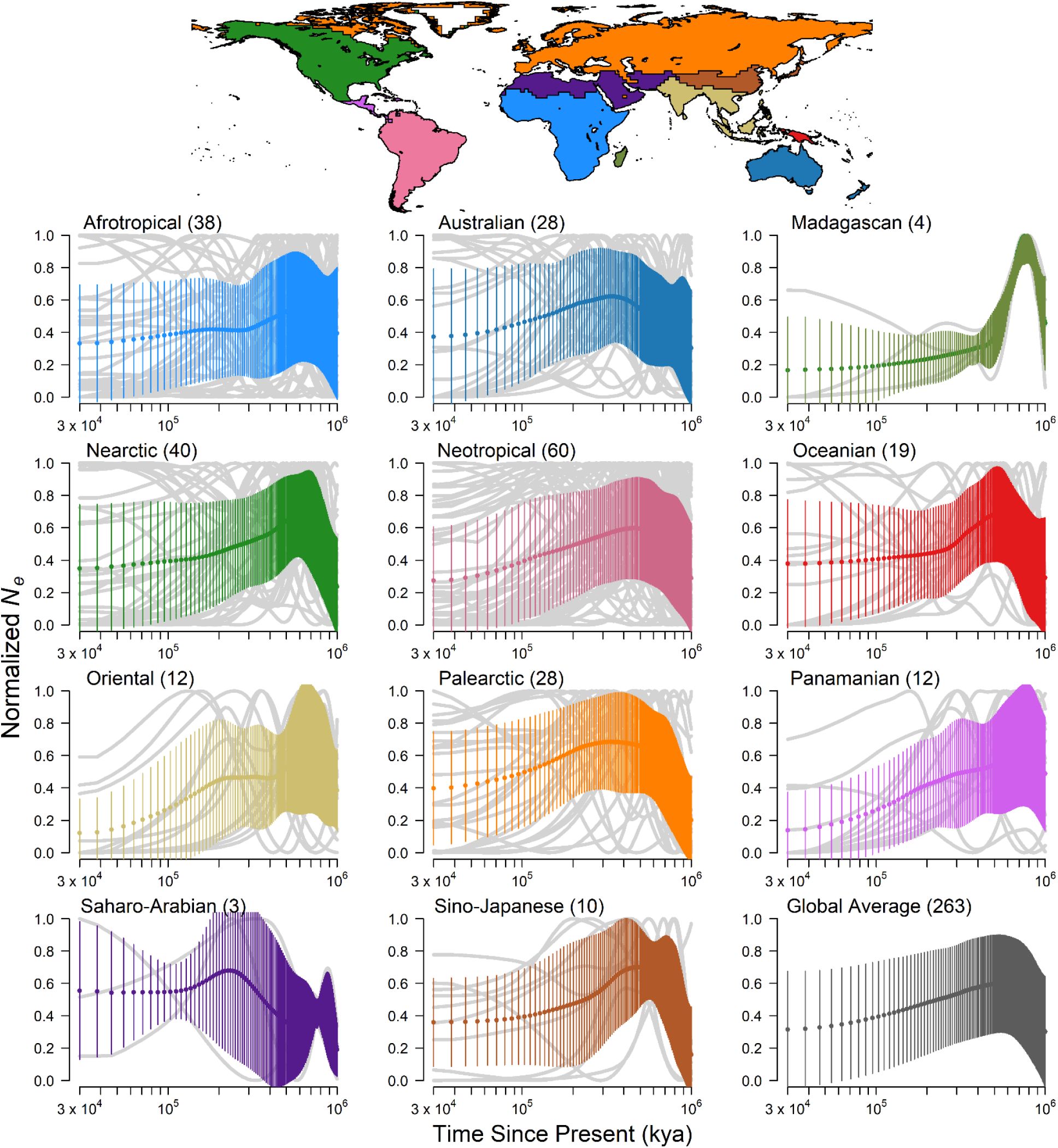
Mean change in normalized *N_e_* from 30,000 to 1 million years ago (30–1000kya) for 263 avian species, summarized by zoogeographic realm (top panel, following Holt et al.^28^). For each panel, the number of species representing each realm is given in parenthesis. Colored dots depict mean normalized *N_e_* at 120 equally-spaced (loglinear) time points from 30– 1000kya, while shaded area depicts SD at each point. Full demographic curves of each species are provided in the background (grey) to show overall variation within each realm.

We used linear mixed-effect models (Gaussian distribution) to first identify the set of morphological and life-history traits most likely to be associated with overall *N_e_* dynamics in response to climate change, for use in downstream analyses (i.e., those explicitly considering phylogenetic and trait interactions, below). For this variable selection step, we designated two periods of relatively recent warming (~147–123kya) and cooling (~122–65kya), which represent some of the most dramatic changes in global climate (i.e., Δ~8°C in Global Average Surface Temperature [GAST] in <60k years) over the past million years (fig. S8), and are within the time window (~30–200kya) in which PSMC-based estimates of *N_e_* are most precise^20^). We quantified demographic sensitivity to climate change as the relationship between species-specific *N_e_* and GAST during these two periods via Pearson correlation coefficients, and designated these relationships as *“Climate Warming”* and *“Climate Cooling”* responses. Of eight initial traits (body mass, brain-body ratio, tarsus length, bill length, egg mass, clutch size, incubation duration, and hand-wing index [HWI – a metric of dispersal ability]) expected to influence these estimates of demographic sensitivity to climate change (Supplementary Text S2), six were identified as key potential influencers (i.e., retained in a subset of ‘best fitting’ models, Table 1) for *Climate Warming* and/or *Climate Cooling* responses, and retained in subsequent analysis. Of these six traits, longer incubation durations and larger clutch sizes were most closely associated with increasing *N_e_* during *Climate Warming,* whereas shorter incubation duration, lower HWI, and longer bill lengths were most closely associated with increasing *N_e_* during *Climate Cooling* (Table 1). While goodness-of-fit for these models were low (*R^2^* = 0.06, 0.05), given the low expectation of variation in a single trait directly influencing *N_e_* responses to climate over evolutionary time scales^22^, these results reveal the suite of key traits among our initial candidate traits which are most likely to be associated with long-term demographic variation under climate change.

**Table 1.** Parameter estimates, standard errors (SE), and upper and lower 95% confidence intervals (UCI, LCI) from averaged linear mixed models (random effect = Passerine/Non-passerine) evaluating the relative effects of morphological and life-history traits on demographic responses to climate warming and cooling (measured as correlation coefficient between *N_e_* change and climate change). For each response, we ran 256 models involving all possible combinations of eight predictor variables (body mass [g], ratio of brain size to body mass, tarsus length [mm], hand-wing index [HWI], bill length [mm], egg mass [g], incubation duration [days], and clutch size; all z-scaled to remove the effects of measurement scale), selected a ‘best models’ subset (ΔAIC ≤ 5 from the best-fitting model) and averaged parameter estimates within this subset. *R^2^* represents the goodness-of-fit of the global model (including all explanatory variables) for each response. Values marked in bold highlight statistically significant predictors (i.e., CIs do not overlap zero).

|  |  | Estimate | SE | z-value | p | LCI | UCI |
| --- | --- | --- | --- | --- | --- | --- | --- |
| *Climate Warming*<br>($R^2 = 0.06$) | Intercept | 0.0003 | 0.07 | | | | |
|  | Incubation duration | 0.15 | 0.08 | 1.92 | 0.06 | -0.003 | 0.30 |
|  | Clutch size | 0.13 | 0.07 | 1.91 | 0.06 | -0.004 | 0.27 |
|  | Egg mass | 0.12 | 0.08 | 1.61 | 0.11 | -0.03 | 0.27 |
|  | HWI | -0.09 | 0.07 | 1.34 | 0.18 | -0.22 | 0.04 |
|  | Body mass | 0.05 | 0.07 | 0.82 | 0.41 | -0.08 | 0.19 |
| *Climate Cooling*<br>($R^2 = 0.05$) | Intercept | -0.0005 | 0.08 | | | | |
|  | Incubation duration | -0.14 | 0.07 | 1.93 | 0.05 | -0.28 | 0.002 |
|  | HWI | -0.14 | 0.07 | 2.00 | **0.05** | **-0.28** | **-0.003** |
|  | Bill length | 0.05 | 0.07 | 0.70 | 0.49 | -0.09 | 0.18 |

Using these six key traits, we further characterize the phylogenetically-explicit interacting trait network of influences on demographic sensitivity to climate change. We categorized each species by their combined *Climate Warming* and *Climate Cooling* responses and used Phylogenetic Path Analysis (PPA). PPA is a hypothesis-driven framework for assessing direct and indirect effects of each trait on differentiating defined species categories, independent of their phylogenetic relationships (fig. S4). Species exhibiting decreasing *N_e_* under *Climate Cooling* and increasing *N_e_* under *Climate Warming* were categorized as “Warming Positive” (Fig 2, table S5). Those exhibiting increasing *N_e_* under *Climate Cooling* and decreasing *N_e_* under *Climate Warming* were categorized as “Warming Negative”, a scenario expected for many temperate and cold-adapted species during the 21^st^ century. We quantified the network of trait effects on differentiating Warming Positive species from all other species in our analysis, and repeated this process for Warming Negative species. Further, we evaluated species which exhibited overall sensitivity to climate warming or cooling (i.e., Warming Positive + Warming Negative responses, hereafter “Climate Sensitive”) versus those with consistent *N_e_* increases or decreases, and those which exhibited consistent decreases in *N_e_* for both the *Climate Warming* and *Climate Cooling* responses (categorized as “Consistent *N_e_* Decrease”) versus remaining species (table S5).

PPA revealed that Warming Positive species were best differentiated by direct effects of reproductive, survival/growth, and dispersal traits (table S6; Core Model D in fig. S11). Averaging the best-performing models from this comparison indicates that *N_e_* of species with larger body masses, lower HWI, and smaller egg masses were more likely to increase under *Climate Warming* and decrease under *Climate Cooling* (Fig. 3A). Interestingly, body mass also had indirect effects on differentiating Warming Positive species from remaining species via its significant positive influence on egg mass and its indirect correlation with a lower HWI (Fig. 3A). These results highlight potential trade-offs between the influences of reproductive, survival/growth, and dispersal traits on changes in population size under climate warming, where the positive effects of larger body mass are offset by the associated increase in egg mass or decrease in HWI. Warming Negative species were also best differentiated via the direct effects of reproductive, survival/growth and dispersal traits, but in the opposite directions compared to what we found in Warming Positive species (table S6; Core Model D in fig. S11). Model averaging further revealed that smaller-bodied species with larger eggs, longer incubation durations, and greater HWI were more likely to exhibit decreasing *N_e_* under *Climate Warming* and increasing *N_e_* under *Climate Cooling,* and again highlighted potential trade-offs between body mass and HWI (Fig. 3B). Trait network effects on differentiating all Climate Sensitive species were less defined, where no traits were found to have significant, direct effects on differentiating both Warming Positive and Warming Negative species from remaining species (Fig. 2C). However, species with consistently decreasing *N_e_* tendency for both the *Climate Warming* and *Climate Cooling* responses were differentiated by smaller clutch sizes and tended to have shorter incubation durations than remaining species (Fig. 3D), providing additional evidence for the directional role of these key traits in mediating demographic responses under climate change.

**Fig. 3.**
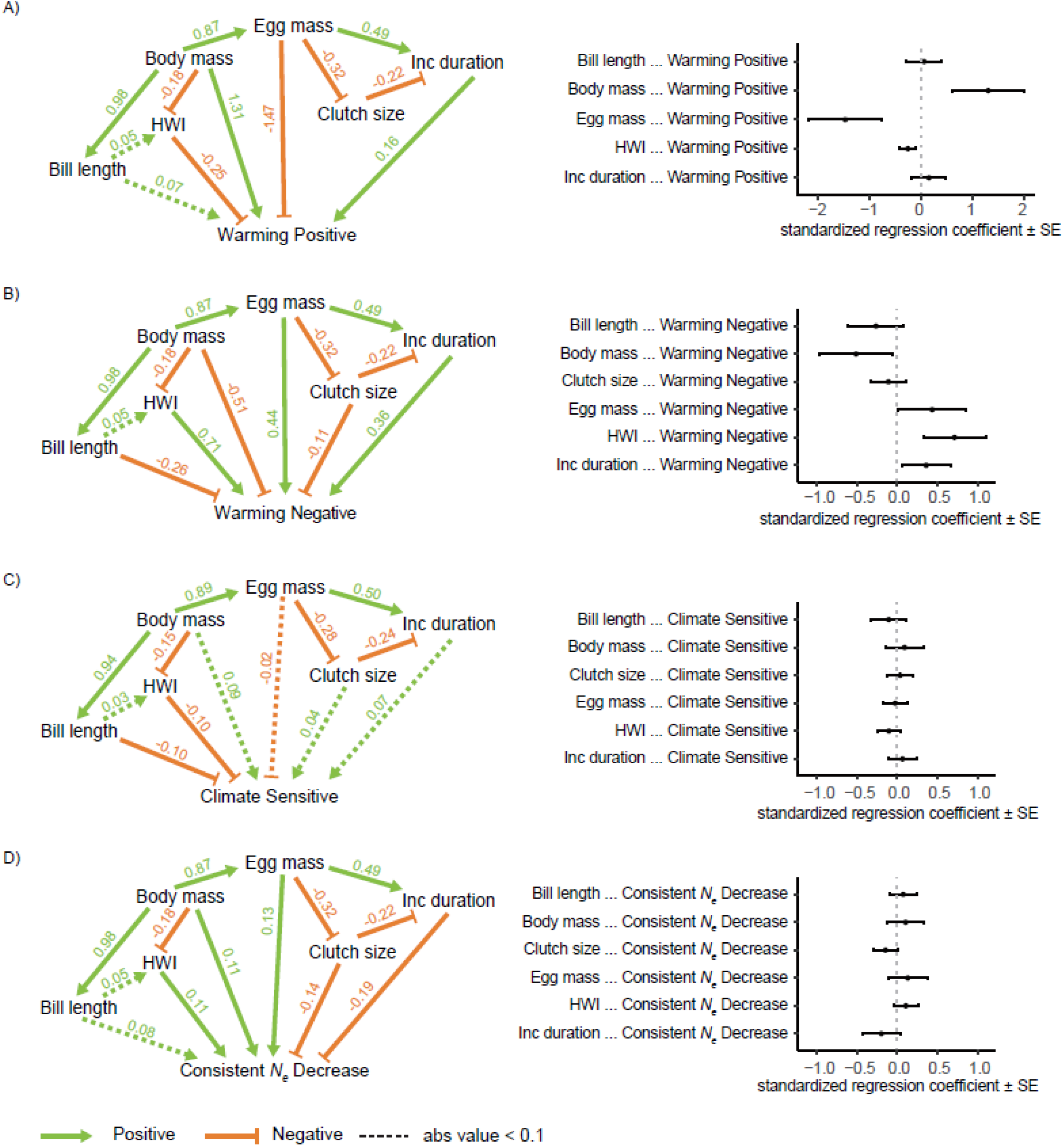
Network effects of key morphological and life-history traits on demographic responses to climate warming and climate cooling, shown as directed acyclic graphs (left) and corresponding standardized regression coefficients (±SE; right) for averaged best performing models. A) Comparison of “Warming Positive” responses (n = 33) versus all remaining species (n = 230) reveals that larger body masses, smaller egg masses, and lower HWI are associated with demographic increases during *Climate Warming* and decreases during *Climate Cooling* (statistical significance of trait effects in all panels assessed by whether SEs overlap zero). B) Comparison of “Warming Negative” responses (n = 29) versus all remaining species (n = 234) reveals that smaller body masses, larger egg masses, greater HWI, and longer incubation durations are significantly associated with demographic increases during *Climate Cooling* and decreases during *Climate Warming.* C) Comparison of all species which exhibited sensitivity to *Climate Warming* or *Cooling* (n = 33 + 29) versus those with consistent *N_e_* increases or decreases (n = 98 + 55) reveals no traits significantly influenced overall species sensitivity to changing climate conditions. D) Comparison of species with consistent *N_e_* decreases during *Climate Warming* and *Climate Cooling* (n = 98) versus all remaining species (n = 165) again revealed no significant influences, although species with smaller clutch sizes, higher HWI, and shorter incubation durations did tend to exhibit demographic decrease during both *Climate Warming* and *Climate Cooling*.

Our observations of trait-network influences on long-term demographic sensitivity to climate change agree with theoretical expectations that larger-bodied, slower-reproducing species with limited dispersal capacity are likely to respond strongly to changing climate (Fig. 2). However, they also suggest that such traits may not necessarily lead to population declines under warming climate conditions as predicted from some empirical observations in contemporary populations^12,13,15,23–25^ (table S11). A lower HWI typical of species with limited dispersal ability was the only predictor found to significantly influence both un-networked (Table 1) and networked (Fig. 2A, B) responses to climate change, where such species tended to increase in abundance under *Climate Cooling* and decrease under *Climate Warming*. We further found that larger body mass had direct and indirect effects on increasing *N_e_* under *Climate Warming* and decreasing *N_e_* under *Climate Cooling* (Fig. 2A, B), contrary to results from contemporary studies (i.e., over ecological time scales, Table S11). Such differences are likely due to the much longer (evolutionary) time scales investigated here via our PSMC analysis (see below).

If morphological and life-history traits play a central role in demographic responses to climate change, as determined from our historical *N_e_* analyses, they may also be reflected in the contemporary global distributions of species. Specifically, trait effects leading to an increasing *N_e_* trend under the *Climate Warming* period may be similar to those of species found in tropical locations today, under the assumption that such species are adapted to warmer climates. Using all available mean trait and breeding/resident range information for 2,745 avian species sampled from around the world, we found that species in tropical latitudes tend to have longer incubation durations and longer bills, but smaller clutch sizes, smaller eggs, lower HWIs, and lower body mass (Supplementary Text S3; fig. S23, S24). Thus, our findings of historical responses to periods of climate warming do partially explain contemporary biodiversity patterns. Specifically, tropical species exhibit lower HWIs, in concordance with species exhibiting such traits showing increasing *N_e_* trends under *Climate Warming* (n = 108; fig. S11, S12). However, the opposing findings of smaller clutch sizes among tropical species and the tendency toward larger clutch sizes among *Climate Warming* responses (averaged model, fig. S12), as well as the remaining traits having no significant direct effect among *Climate Warming* responses, indicate that historical responses to climate change alone are only partially predictive of contemporary distributions.

Our study reveals seven main demographic patterns among a broad geographic (Fig. 2) and phylogenetic (fig. S7) sample of birds over the past million years, and a key trait-adaptive network associated with population responses to long-term climate change in the absence of additional human impacts. Unlike short-term responses to climate change (e.g., studies listed in Table S11) where single traits dictate adaptive functions under strong natural selection^12,15^, long-term adaptation (i.e., over evolutionary timescales) is influenced by overall genomic variation within species, where the effects of individual traits become saturated and diminish over time^22,23,26,27^. This is reflected both in the very low goodness-of-fit associated with individual trait effects (i.e., un-networked) models (Table 1) and in trade-offs among several traits in PPA analysis of responses to periods of climate warming and cooling (Fig 3, fig. S11, S12). Over hundreds of thousands of years, interacting trait networks and trade-offs among survival/growth and reproductive traits may develop which potentially mask direct trait effects identified in studies of contemporary populations. Reconstructing long-term population dynamics from genomic data is thus a crucial component of revealing how past climatic events influenced the genetic makeup of contemporary populations over time, and for providing demographic baselines before the Anthropocene^5,6^. Such analyses across the tree of life will provide unique insight into the natural variability of long-term demography, and help direct conservation efforts towards species more sensitive to broad-scale global environmental change.

## Acknowledgments

This work was made possible by the generous efforts of field biologists and museum staff who contributed samples to the B10k project.

## Author contributions

R.R.G., S.F., G.Z. and D.N-B. designed the study and wrote the paper. R.R.G., S.F., and G.C. carried out analyses. All authors contributed towards the assembly of genomic and morphological/life-history data, discussion of results, and reviewing and editing the manuscript.

## Data and materials availability

All data and code underlying these analyses are available in the Dryad Digital Repository^29^.

## Supplemental Materials

Materials and Methods

Supplementary Text S1 to S3

Tables S1 to S12

Figures S1 to S24

References (*30–104*)

## References

1. Butchart, S. H. M. et al. Global Biodiversity: Indicators of Recent Declines. Science 328, 1164–1168 (2010).

2. Dirzo, R. et al. Defaunation in the Anthropocene. Science 345, 401–406 (2014).

3. IPBES. Global assessment report on biodiversity and ecosystem services of the Intergovernmental Science-Policy Platform on Biodiversity and Ecosystem Services. (IPBES secretariat, 2019).

4. Rosenberg, K. V. et al. Decline of the North American avifauna. Science 366, 120–124 (2019).

5. Fordham, D. A. et al. Using paleo-archives to safeguard biodiversity under climate change. Science 369, eabc5654 (2020).

6. Nogués-Bravo, D. et al. Cracking the Code of Biodiversity Responses to Past Climate Change. Trends Ecol. Evol. 33, 765–776 (2018).

7. Foden, W. B. et al. Climate change vulnerability assessment of species. WIREs Climate Change 10, e551 (2019).

8. Chattopadhyay, B., Garg, K. M., Ray, R. & Rheindt, F. E. Fluctuating fortunes: genomes and habitat reconstructions reveal global climate-mediated changes in bats’ genetic diversity. Proc. R. Soc. B. 286, 20190304 (2019).

9. Chen, L. et al. Large-scale ruminant genome sequencing provides insights into their evolution and distinct traits. Science 364, (2019).

10. Peart, C. R. et al. Determinants of genetic variation across eco-evolutionary scales in pinnipeds. Nat. Ecol. Evol. 4, 1095–1104 (2020).

11. Jenouvrier, S. Impacts of climate change on avian populations. Glob. Change Biol. 19, 2036–2057 (2013).

12. Pacifici, M. et al. Species’ traits influenced their response to recent climate change. Nat. Clim. Chang. 7, 205–208 (2017).

13. Foden, W. B. et al. Identifying the World’s Most Climate Change Vulnerable Species: A Systematic Trait-Based Assessment of all Birds, Amphibians and Corals. PLOS ONE 8, e65427 (2013).

14. Telenský, T., Klvaňa, P., Jelínek, M., Cepák, J. & Reif, J. The influence of climate variability on demographic rates of avian Afro-palearctic migrants. Sci. Rep. 10, 17592 (2020).

15. Jiguet, F., Gadot, A.-S., Julliard, R., Newson, S. E. & Couvet, D. Climate envelope, life history traits and the resilience of birds facing global change. Glob. Change Biol. 13, 1672–1684 (2007).

16. Lorenzen, E. D. et al. Species-specific responses of Late Quaternary megafauna to climate and humans. Nature 479, 359–364 (2011).

17. Hung, C.-M. et al. Drastic population fluctuations explain the rapid extinction of the passenger pigeon. Proc. Natl. Acad. Sci. U.S.A. 111, 10636–10641 (2014).

18. Feng, S. et al. The Genomic Footprints of the Fall and Recovery of the Crested Ibis. Curr. Biol. 29, 340–349.e7 (2019).

19. Feng, S. et al. Dense sampling of bird diversity increases power of comparative genomics. Nature 587, 252–257 (2020).

20. Li, H. & Durbin, R. Inference of human population history from individual whole-genome sequences. Nature 475, 493–496 (2011).

21. Fjeldså, J., Christidis, L. & Ericson, P. G. P. The Largest Avian Radiation. The Evolution of Perching Birds, or the Order Passeriformes. (Lynx ed, 2020).

22. Bay, R. A. et al. Genomic signals of selection predict climate-driven population declines in a migratory bird. Science 359, 83–86 (2018).

23. Parmesan, C. Ecological and Evolutionary Responses to Recent Climate Change. Annu. Rev. Ecol. Evol. Syst. 37, 637–669 (2006).

24. Angert, A. L. et al. Do species’ traits predict recent shifts at expanding range edges? Ecol. Lett. 14, 677–689 (2011).

25. McCain, C. M. & King, S. R. B. Body size and activity times mediate mammalian responses to climate change. Glob. Change Biol. 20, 1760–1769 (2014).

26. Barrett, R. D. H. & Schluter, D. Adaptation from standing genetic variation. Trends Ecol. Evol. 23, 38–44 (2008).

27. Hancock, A. M. et al. Adaptation to Climate Across the Arabidopsis thaliana Genome. Science 334, 83–86 (2011).

28. Holt, B. G. et al. An Update of Wallace’s Zoogeographic Regions of the World. Science 339, 74–78 (2013).

29. Germain, R. et al. Data from: Species-specific traits mediate avian demographic responses under past climate change. Dryad, Dataset, http://datadryad.org/stash/dataset/doi:10.5061/dryad.fn2z34tz8 (2022).

