## Supporting Information for "Species-specific traits mediate avian demographic responses under past climate change"

**This PDF file includes:**

Materials and Methods

Supplementary Text S1 to S3

Tables S1 to S12

Figures S1 to S24

References (30–104)

**Materials and Methods**

SNP calling, and mutation rate estimation

Whole-genome sequencing data for 345 bird species were collected by Feng et al.[19] and released as genomic resources of the Bird 10,000 Genomes (*B10K*) Project Phase II (https://b10k.genomics.cn). Genomes for each species were generated under a standardized protocol of library building, sequencing, and assembly as part of this previous study, thereby minimizing potential bioinformatic artefacts which could affect our analyses and interpretations. We inferred heterozygous information of each species based on whole-genome sequencing data by using BWA+GATK pipeline[20,30]. Four filtering steps were applied to obtain high-quality SNPs[31], including: a) removing homozygous SNPs (i.e., those where the genotype was encoded as 'Minor/Minor' in the variant call format file) and SNPs with more than two alternative alleles, b) removing SNPs with an interval below 10bp, c) removing SNPs with a read depth below 1/3 or over twice the average read depth across the genome, and d) removing SNPs with a root-mean-square mapping quality lower than 25.

We used branch-specific estimates of the substitution rate per site ($R$) from a dated phylogeny provided by the latest B10K family-phase phylogenetic study as proxies for mutation rate ($\mu$). This latest topology was inferred based on the whole-genome data of 363 species, assumed to be the most effective in addressing deep evolutionary relationships[32–34]. Divergence times ($t$) were estimated by MCMCTree[35] with a large number of fossil records to provide by far the best calibration information[36,37]. Both of these measures ensure the highest-possible accuracy of species-specific substitution estimates, as well as divergence times. To further convert the unit of mutation rate from per site to per site per generation, we scaled the species-specific mutation rates ($\mu$) as $\mu = \frac{R}{t} \times T$, where T is the generation time for each species.

Demographic reconstruction over the past one million years

We used the Pairwise Sequential Markovian Coalescent (PSMC) method[20] to reconstruct long-term changes in effective population size ($N_e$) for each species. Diploid genomes consist of thousands of independent loci; the coalescent approach underlying PSMC estimates the time to the most recent common ancestor (TMRCA) of the two alleles at each locus, creating an overall TMRCA distribution across the genome. The rate of coalescent events is inversely proportion to $N_e$, therefore the PSMC method can identify periods of changing $N_e$ over time (e.g., if many loci were observed to coalescent at a given time point, it would indicate a lower $N_e$ at that time)[20]. To ensure reliability of PSMC analyses, we removed 20 species with lower-quality heterozygosity information (i.e., <18x genome-wide coverage or >25% missing data, as recommended by Nadachowska-Brzyska et al[38]). For the remaining 325 qualified species (table S1), we used the PSMC settings "-N30 -t5 -r5 -p 4+251*1+4+6+10" with a reduced dataset, and then scaled results to real time using estimated mutation rate (above) and generation time[39]. This set of parameters had an increased number of free atomic intervals (-p) on the basis of previous study[31], which could generate more PSMC estimates without changing the shape of the PSMC curves (fig. S1). Data representing $N_e$ estimates (x$10^4$) scaled to real time via PSMC for all 325 species are available in the Dryad Digital Repository[29]. Because recent and drastic population bottlenecks, and associated severe inbreeding, can lower PSMC-based estimates of $N_e$ in more recent time periods and erase information regarding ancient $N_e$ dynamics, we restricted downstream analyses to the 263 species with full demographic coverage over the focal time period (30kya–1mya) for which we had the most confidence in PSMC-based estimates of effective population size. Of the 62 species excluded via this step for which conservation status was available, 19 are currently listed as 'threatened' or 'near-threatened' (IUCN Redlist; www.iucnredlist.org), indicating a tendency ($X^2$ = 10.78, p-value = 0.001) towards such species to have limited polymorphism information needed to quantify long-term demographic history, once again likely due to recent population bottlenecks.

Since its initial development[20], the PSMC method has been widely applied across taxa to detect changes in population sized over time, based on whole-genome sequence information. Because PSMC does not rely on population data and can straightforwardly construct historical $N_e$ dynamics over time, it is favored among *de novo* genome projects for its accuracy and precision[40], particularly regarding demographic changes in the more distant past[41–44]. Although there are some generally accepted filtering strategies for sequence data to improve the accuracy of demographic reconstructions[38], potential biases in PSMC output can be the result of both scaling parameters (e.g., mutation rate) and population structure. For the former type, changes in the scaling parameters cause the PSMC curve to move along the axes while maintaining the overall curve shape[31]. To account for this, we performed a robustness test of the PSMC curves in terms of sampling mutation rates from the posterior distributions on a dated phylogenetic tree (see below, fig. S1). For the later type of bias, theoretical work has shown that the accuracy and interpretation of PSMC results can be affected by population structure, in that changes in connectivity among sub-populations may influence overall patterns of demographic change[45–47]. While we have no *a priori* information on changes in connectivity over the last million years for our 263 species, we assume that the patterns observed are likely the result of a combination of changes in both population size and a degree of changes in population connectivity, both of which would be mediated by the effects of trait networks under periods of climate change (i.e., species with stronger dispersal ability [Hand-Wing Index] maintaining greater connectivity during periods of extreme environmental change). Thus, while not infallible, our analyses and results present a key step forward in our understanding of how trait networks can mediate demographic responses to periods of acute climate warming and cooling over the past million years.

Cluster analysis of demographic patterns and robustness tests

To investigate patterns in overall demographic fluctuations among the 263 bird species, we applied clustering analysis based on normalized $N_e$ values. We first selected 121 time points with equal time intervals (after log10 transformation) from 30 to 1,000kya. For each species, we extracted the corresponding $N_e$ values for these 121 time points along the species-specific $N_e$ trajectory and used min-max normalization to rescale values between 0 and 1. These normalized $N_e$ values resulted in a 263*121 matrix, which was then used as the input for a hierarchical clustering analysis based on the Euclidean distance performed in *pheatmap* (with 'cluster_row = T'). Clustering methods aim to define clusters such that the total intra-group variation is minimized. In this instance (e.g., Fig. 1A), species with similar $N_e$ trajectories are found closer together on the clustering dendrogram. To summarize the major demographic patterns among species for the past million years, we used the 'cutree' function in the R package *dendextend*[48] to split the resulting clustering dendrogram into 3, 4, 5, 6 and 7 groups (i.e., k = 3–7), respectively (figs. S2–S5). Compared with a lower value of k (e.g., k = 3, fig S2), dividing the dendrogram into k = 7 subtrees (Fig. 1A) made the species contained in each subtree more compact in terms of $N_e$ pattern consistency, and was sufficient to clearly identify the major fluctuation patterns.

Different methods of data transformation can lead to different interpretations of both the appropriate number and organization of clusters. The min/max normalization approach used here focuses on the mode of demographic fluctuations, whereas other methods (e.g., rescaling the mean) take into account both the mode and degrees of change. Because our analyses center on the positive vs. negative relationships between demographic change and climate change during periods of warming and cooling (below), our normalization method should likewise focus on the modes of positive/negative demographic responses, rather than the degrees of change. However, to examine the robustness of our clustering analysis to the choice of data transformation, we used the 'TreeDistance' function in the R package *TreeDist*[49] to calculate the distance (0-1, where higher values indicate more similarity) between the output clustering trees generated after the min/max normalization approach and three other data transformation methods (rescaling the mean, Z-transformation, and coefficient of variation).The distance value between the min/max normalization and Z-transformation was 0.83, and the distance value between the min/max normalization and the coefficient of variation was 0.82, indicating that the cluster designations from these three separate methods were relatively consistent. While the distance value between min/max normalization and rescaling the mean was relatively lower (0.66), indicating less consistency between clustering results, this was expected given the points made above (rescaling being based on combining the modes and degrees of fluctuations). We further used Rezende's 'phylo.signal.disc' algorithm[50] to examine the phylogenetic signal underlying cluster designations from these four data transformation methods (see below for full results from min/max normalization). For rescaling the mean, the observed number of transitions was 153, which was not significantly different than expected by chance (randomized mean transitions = 154, p = 0.46), indicating that phylogeny alone did not explain the observed differences among demographic patterns. Similarly, observed transitions for the Z-transformation and coefficient of variation methods did not significantly differ from chance (p = 0.21 and 0.99, respectively). Thus, in terms of phylogenetic signal testing, all alternative normalization methods investigated returned consistent results/conclusions to the min/max normalization we used throughout our analyses.

Considering that transforming PSMC outputs into real time is sensitive to the value of mutation rates[20], we let the species-specific mutation rates vary within a reasonable range to generate random input matrixes and assess the robustness of the above clustering result. As a time-estimation algorithm based on Bayesian theory, MCMCTree provides the posterior age distributions for each node[35]. Therefore, we first randomly sampled 100 estimates of the divergence time from the posterior distribution of nodes corresponding to each species. With the fixed substitution rates and generation times, 100 mutation rates were calculated for each species using these random values based on the formulas provided above and then used to scale the PSMC output from the coalescent unit to the real time unit. This gave 100 $N_e$ trajectories per species based on varying mutation rates. Next, we randomly selected one of these trajectories from each species and extracted the same 121 time points as before to form a new 263*121 matrix. This step was repeated 100 times and the resulting 100 matrixes were next used to produce the clustering results, as well as the split results under different number of groups (k). When k = 7, we first randomly selected 500 pairs of species and obtained the information on whether each pair of species belonged to the same group in the clustering dendrogram shown in Fig. 1A. Then, for the same pair of species, we checked how many times the clustering results obtained from 100 random matrixes are consistent with the Fig. 1A clustering dendrogram, in terms of the group information. For example, species A and species B were from the same group in Fig. 1A, and they were also considered to be from the same group in 100 random clustering results. The consistency ratio for these two species was then calculated as a percentage. The average consistency ratio for all 500 pairs of species was used to represent the robustness value for k = 7, and we repeated this process for all other k values (Table S2). Robustness values increased with the number of clustering groups used, and was 83.49% when k = 7. A higher k value implies higher intra-group similarity and lower inter-group similarity. Thus, a pair of species from the same group at k = 7 is more likely to have the sufficient similarity of $N_e$

trajectories to be clustered together even when there is some estimation bias in the mutation rate, compared to species from the same group at k = 4. Given that the major fluctuation patterns are identified and the robustness is over 80% at k = 7, we used splitting results from this grouping number in subsequent analysis (Fig. 1A).

We next ran a Chi-square test to examine whether the splitting pattern for k = 7 could be explained in part by when species within each cluster group reached their maximum effective population size (here, normalized $N_e$ value greater than 0.9 is considered as a maximum value; table S3). Compared to expected background patterns, the $N_e$ fluctuations of species in Group 1, Group 3 and Group 4 tended to reach their maximums during the upper Pleistocene (30–129kya), while Group 6 and Group 7 reached their maximums in the middle Pleistocene (129–774kya), and species in Group 5 reached their maximums during the middle and lower Pleistocene (774–1,000kya; table S3). Unlike these groups, Group 2 did not exhibit significant differences from expected background patterns during any of the time periods. To further determine the association between clustering and mean effective population size, we averaged $N_e$ values over 30-1000kya for each species and quantified intra-group and inter-group differences (ANOVA) based on the k = 7 clustering groups. From these comparisons, we detected significant differences in mean $N_e$ between Group 5 & Group 1 and between Group 5 & Group 3, where individuals clustered in Group 5 had lower mean $N_e$ over time than those in Groups 1 & 3, but where all other pairwise comparisons revealed no significant differences in mean $N_e$ over time (fig. S6).

To analyze how lineages may resemble each other in overall demographic fluctuations (i.e., whether phylogenetically related species are classified into the same cluster groups), we used Rezende's 'phylo.signal.disc' algorithm[50]. This algorithm compares the minimum number of character-state transitions at each node that account for the observed character distribution in the phylogeny, assuming maximum parsimony with the median of a randomized distribution (1000 randomizations were used). If the observed evolutionary transitions are significantly less than the randomized median, a phylogenetic signal is inferred. The observed transitions were 171 when k = 7, which were not significantly lower than expected by chance (the randomized median transitions were 175, p-value = 0.21, fig. S7A). Even when some phylogenetically related lineages were observed to have similar $N_e$ trajectories under k = 4 (red background in fig. S7B), the similarities were still not significant (observed transitions = 123, randomized median transitions = 128, p-value = 0.13; fig. S7B). These results indicates that there are factors other than phylogeny that contribute to the convergent patterns of demographic fluctuations.

Quantifying demographic responses to recent periods of climate warming and cooling

The last one million years of the Earth's history is punctuated by periods of abrupt climate warming and cooling[51], the most dramatic of which have occurred in relatively recent paleo-ecological time (cooling from ~122–65kya; warming from ~147–123kya; fig. S8). Such periods of more recent paleo-ecological time also correspond to when PSMC-based estimates of $N_e$ provide more detailed representations of demographic fluctuations within a defined window of time, and thus allow for the most accurate quantification of the relationship between climate change and changing $N_e$ for each species.

To quantify the effects of climate warming and cooling on species-specific demographic trends over time, we first obtained the real time points and $N_e$ values scaled from PSMC estimates for each individual within these two climate periods, and inferred the climate values for these time points from Snyder[51]. Snyder[51] presented climate data over the last million years (and beyond; estimated from a spatially-weighted, multi-proxy database of over 20,000 sea surface temperature reconstructions) as the change in global average surface temperature (one value per thousand years), for example: -6.12 in 65kya and -5.99 in 66kya. Thus, we could infer the climate value corresponding to any time point between two adjacent time points based on the slope of the line between them. For example, based on the above values, the climate value of 65.70kya is calculated as -6.029. We then quantified overall $N_e$ responses to warming and cooling (hereafter designated "*Climate Warming*" and "*Climate Cooling*") via Pearson correlation coefficients for two variables: $N_e$ estimate and climate value. Following David[52], the recommend sample size for running Pearson's $r$ is 25, or higher. In our study, the sample size is the number of corresponding $N_e$ estimates and climate values for each individual during *Climate Warming* or *Climate Cooling*. Under the PSMC settings "-N30 -t5 -r5 -p 4+251*1+4+6+10", the average sample size in *Climate Cooling* was 26.91, while *Climate Warming* only contained 8.60 $N_e$ estimates in average due to a shorter period. To avoid the under-powered calculations caused by such a small sample size in *Climate Warming,* we reran the PSMC analysis with the settings "-N30 -t5 -r5 -p 4+800*1+4+6+10". These modified parameter settings allowed us to increase the average number of samples to 26.03 in *Climate Warming*, without changing the shape of the PSMC curves (fig. S1). In addition, we used 0.55 as the threshold for correlation coefficient to indicate statistical significance of a correlation, based on the algorithm described by Guenther[53] with *Power* = 80%, *Alpha* = 0.05 and *Sample Size* = 25. Guenther's algorithm also allowed us to perform a further analysis on whether the correlation coefficients calculated for few individuals with sample sizes less than 25 were significantly different from zero. If the sample size of such species is smaller than the minimum sample size estimated from its correlation coefficient based on Guenther's algorithm, we consider this correlation coefficient to be 0. For example, the minimum sample size is 13 for Pearson's r = 0.7, when *Power* = 80% and *Alpha* = 0.05. Therefore, if there were only 10 $N_e$ estimates for a species, we cannot accept the correlation between $N_e$ estimates and climate values even if its Pearson's *r* was as high as 0.7. After passing the above criteria, a positive correlation (at $p < 0.05$) corresponded to $N_e$ tendency tracking changing temperature (i.e., increasing $N_e$ under increasing temperature or decreasing $N_e$ under decreasing temperature). All significant positive/negative correlations were visually inspected to ensure they represented a predominantly linear relationship. The above analysis criteria are outlined visually in fig. S9.

To test the robustness of our estimated demographic responses to climate change, we randomly generated 1,000 mutation rates for each individual using the same strategy as the clustering robustness analysis (above). This gave 1,000 $N_e$ trajectories per species based on changes in mutation rates. By using the same quantitative criteria as our observed response results (fig. S9), we further assigned response labels to the estimated 1,000 trajectories per individual during the *Climate Warming* and *Climate Cooling* periods, respectively. Based on these response labels, we used the confidence level to represent the robustness, which is the percentage of response labels obtained from 1,000 random sampling events that are consistent with the observed response results for each individual. For example, for a species exhibiting a positive correlation in our observed results, if its response labels inferred from the random sampling were 950 positives and 50 negatives, its confidence level was 0.95. In summary, the mean confidence level is 0.90 during *Climate Warming*, and 0.92 during *Climate Cooling*, which implied strong confidence in our observed demographic response estimates. Three types of demographic responses ("Increase", "Decrease", "Unrelated"; determined by the strength of positive/negative correlations at different confidence levels) are summarized in table S4.

For later inclusion in Phylogenetic Path Analysis (below), we categorized each species by their combined *Climate Warming* and *Climate Cooling* responses. Species which exhibited decreasing $N_e$ under *Climate Cooling* and increasing $N_e$ under *Climate Warming* were categorized as "Warming Positive" (n = 33), while those with the opposite response ($N_e$ increase under *Climate Cooling* and decrease under *Climate Warming*) were categorized as "Warming Negative" (n = 29; table S5). There were 48 species where demographic change was not correlated with changing temperature during either *Climate Warming* and/or *Climate Cooling* (categorized as "$N_e$ independent of climate change"). The remaining 153 species either showed consistent $N_e$ increases under both *Climate Warming* and *Climate Cooling* (n = 55), or consistent $N_e$ decreases under both these periods of climate change (n = 98; table S5).

Quantifying the relative influence of individual traits on population responses to climate change

We used linear mixed-effect models (Gaussian distribution) and multi-model inference via the *lme4*[54] and *MuMIn*[55] R packages as a variable selection step to identify the set of morphological and life-history traits most likely to be associated with population responses during the periods of *Climate Warming* and *Climate Cooling* (based on Pearson correlation coefficients, above). For each response variable, we constructed a set of models with Passerine/Non-passerine as a random effect to account for Passerines exhibiting both greater mean and greater variance in $N_e$ values across the full study period (Fig. 1B), since we aimed to identify traits linked to demographic responses to climate change while reducing the potential effects of broad-scale phylogenetic signal on such results (see Phylogenic Path Analysis below). Each global (i.e., all variables included) model also included eight non-collinear morphological/life-history traits predicted to influence population responses to climate change (Supplementary Text S2) as fixed effects. These eight traits (selected from an initial set of 17 candidate traits, Table S11) were: mean unsexed mass or mean of male and female masses (g, 'body mass'), ratio of brain size to body mass ('brain-body ratio'), mean unsexed tarsus length or mean of male and female tarsus length (mm, 'tarsus length'), mean unsexed bill length or mean of male and female bill length (mm of total exposed culmen, 'bill length'), mean mass of fresh eggs (g, 'egg mass'), mean number of eggs per clutch ('clutch size'), duration of clutch incubation (days, 'incubation duration'), and 'Hand-Wing Index', measured as Kipp's Distance (distance [mm] between the tip of the first secondary feather to the tip of the longest primary feather) divided by total wing chord length (length [mm] from bend of the wing to the longest primary of the unflattened wing) and multiplied by 100. All fixed effects were standardized to mean = 0, sd = 1 to reduce potential influence of measurement scale on results and to allow direct comparison of model coefficients[56]. We further confirmed key model structure assumptions of homogeneity of variance (residuals vs. predicted values) and normal distribution of residuals (Q-Q plots) for each global model. Goodness of fit ($R^2$) for each model was assessed by the conditional coefficient of determination[57]. We then ran n = 256 models for all possible combinations of these eight fixed effects for each response variable (correlation coefficient during *Climate Warming* and *Climate Cooling*, respectively) and selected a subset with a difference in Akaike Information Criterion (ΔAIC) ≤5 from the best-fitting model. For each response variable, we then averaged parameter estimates for each predictor included in this subset of models to create one representative (full-average) estimate of the relative effects of each component on $N_e$ responses to *Climate Warming* and $N_e$ responses to *Climate Cooling* (Table 1). Statistical significance for each fixed effect was assessed by whether 95% confidence intervals (CIs) overlapped zero.

<u>Phylogenetic Path Analysis to identify trait network effects on demographic responses to climate change</u>

We implemented the hypothesis-driven framework of Phylogenetic Path Analysis (PPA; *phylopath* R package[58]) to quantify the network of direct and indirect effects of key morphological/life-history traits (identified via multi-model inference; above) on demographic responses to climate change while accounting for phylogenetic non-independence of species. PPA allows for a large number of models with complex configurations and inter-correlation of variables. In our study, six continuous variables representing our key morphological/life history traits (egg mass, clutch size, incubation duration, body mass, bill length, and hand-wing index) and one binary variable representing differing demographic responses to climate change (see below) were included in each model. A total of 14 core models with different configurations of these variables were evaluated for each of our four comparisons using both p-values and the C-statistic Information Criterion (CICc) corrected for small sample sizes (fig. S10). Models were designed to test all possible networks of the traits and their effects on the demographic responses. Since the "Reproduction" category contained three traits (egg mass, clutch size, and incubation duration), core models assuming a direct effect of "Reproduction" on demographic responses in fact have seven derived models (i.e., at least one of the three traits having a direct effect on the demographic responses; fig. S10). The same is true for the "Survival/Growth" category with two traits (body mass and bill length). Thus, depending on the network assumed by each core model, one core model would further be modified into 1, 3, 7 or 21 submodels.

In total, we implemented 164 models for four comparisons of the overall responses of species to *Climate Warming* and *Climate Cooling* (table S5) at different confidence levels to again assess the robustness of our main results (i.e., those assessed without confidence level restrictions): 1) "Warming Positive" responses versus all remaining species, 2) "Warming Negative" species versus all remaining species, 3) species which exhibited overall sensitivity to climate warming or cooling (i.e., Warming Positive + Warming Negative species) versus species with consistent $N_e$ increases or decreases for both the *Climate Warming* and *Climate Cooling* responses, and 4) species which exhibited consistently decreasing $N_e$ versus all remaining species. For each comparison, PPA provides the number of independence claims made by the model, the number of parameters, the C-statistic, and the accompanying p-value, where significance (at $p < 0.05$) indicates that the available evidence rejects the model (i.e., the model does not provide a good fit to the data[59]). Thus, after discarding rejected models we calculated the top-ranked model (based on ΔCICc) and the average of the best performing models (ΔCICc ≤

2) for each comparison. Detailed results of the best performing models for each confidence level are provided in tables S6-10, with their associated directed acyclic graphs and regression coefficients provided in figs. S11-20.

**Supplementary Text S1: Geographic variation in demographic histories**

At a global scale, close distributional and phylogenetic relationships of species experiencing similar geographic and climatic conditions (i.e., zoogeographic realms) are a cornerstone for macro-ecological studies[60,61]. Species occupying a common zoogeographic realm may experience similar levels of environmental variation, and thus may have similar demographic responses, compared to species with the same life-history strategies in other areas of the world[16]. Because geographic variation in the magnitude of changing climate may dictate the context and severity of selection acting on morphological and/or life history traits in different regions of the world[62,63], broad-scale studies of phylogenetically distinct groups native to differing geographic areas may facilitate tests of the mediating role of traits on demographic responses to climate change. We evaluated variation in species-specific demographic histories across zoogeographic realms to quantify the extent of possible geographic structure in our clustering of demographic patterns (Fig. 1A, fig. S21), and determine if species in certain zoogeographic realms exhibited atypical patterns of demographic change over the past one million years.

We assigned all 263 species in our analysis to one of the 11 main zoogeographic realms identified via multi-taxon species assemblages by Holt et al.[28] (Fig 2). Following this previous work, migratory species were assigned to the realm corresponding to their breeding range, and species with larger ranges extending across multiple realms were assigned to the realm where the individual sampled for whole-genome sequencing originated. To quantify whether realm-specific demographic patterns deviated from random expectations and if our designation of a given species to a given realm influenced our results, we performed a modified permutation test for the six realms that contained at least 19 species. For each of these six realms, we first generated a distance matrix among samples based on the focal realm and the remaining species. For instance, species in the Oceanian realm (n = 19) represent a subset of the 263*121 matrix of normalized $N_e$ values generated as part of our analysis (see Materials and Methods – 'Demographic reconstruction over the past one million years'). We generated a distance matrix containing differences in the $N_e$ fluctuations between each Oceanian sample and every remaining species using the 'distmat' function in the R package *pracma*[64]. Next, we calculated the overall variance among these distances, which represents the discrete distance in values between these 19 samples and all remaining species. If individuals within the Oceanian realm exhibited substantially different $N_e$ trajectories compared to all remaining species (e.g., they formed a subtree of the cluster tree in Fig. 1A), this process would produce distances with similar values, which would result in a smaller variance. Another scenario is that $N_e$ trajectories of some species from the Oceanian realm are more similar than other Oceanian birds (e.g., they formed multiple subtrees). In this case, the variance increases as the intra-realm groupings are more visible. We then generated a background distribution of potential variances by randomly sampling 19 species from the 263*121 matrix, calculating the variance in distances between this subset and remaining species in the matrix (as above), and repeating this process 1,000 times. If the actual variance between $N_e$ trajectories from the Oceanian realm and remaining species was found to be greater than or equal to the 95$^{th}$ percentile values generated from this background distribution, the demographic histories of species in the Oceanian realm would be considered to exhibit significant regionality. We repeated this process for the remaining five realms with n ≥19, using their respective sample sizes to generate the actual and background distance matrices. Results from the above analysis indicate little geographic variation in demographic trends among realms, and that assignment to a given geographic realm (i.e., among species with distributions spanning multiple realms) had no significant effect on our results and interpretations (fig, S22).

**Supplementary Text S2: Selection and measurement of morphological and life-history traits**

Trait data for all species involved in our study were collected from a combination of live-caught individuals, museum specimens, and existing published databases following previous comparative studies of avian morphological and life-history variation[65–67]. Our initial set of potential morphological and life-history traits consisted of 17 traits typically found to exhibit links with demographic responses to climate change in contemporary studies (see Table S11 for full description of all traits). Because equal observation numbers are required to compare AIC values among competing models[68], we used the *missForest* R package[69] to impute missing values (via a random forest algorithm) for some traits with less than 100% complete records across all species (mass = 98%, clutch size = 84%, egg mass = 74%, incubation duration = 64%, brain size = 57%, generation time = 96% [but see below]), based on phylogenetic relatedness among species[69–71]. Normalized root mean squared error for missing value imputation was 0.038, indicating high accuracy in estimating known trait values (where values closer to zero indicate good performance of the random forest algorithm and values close to 1 indicate low accuracy in estimating known trait values)[69]. Prior to imputing missing values, we excluded both maximum longevity and mortality rate from further analyses as each had less than 40% complete records, and generation time because species-specific generation times factored into our PSMC-based estimates of $N_e$ over time (see Materials and Methods). Further, we excluded range size and elevation min/max since these represent more fluid traits that are highly influenced by measurement under contemporary climate parameters, and may not represent the historic geographic and elevation niches used by species under past instances of climate warming and cooling across the globe. All remaining traits are assumed to remain consistent within these distinct species over the past million years, as this represents a relatively brief period in avian evolution[72]. Further, we have no *a priori* rationale to suggest that species have undergone directional changes in morphological or life-history traits over this period of cyclical climate warming and cooling, thus any trait changes over time may add additional noise but are unlikely to bias our results and interpretations.

We tested for multicollinearity among the remaining 12 traits using Pearson pairwise correlation analysis[73] and identified those which were co-linear at a level of $r \geq 0.7$. For these pairs of co-linear traits, we calculated univariate ANOVAs (with mean overall correlation coefficients between $N_e$ and GAST during periods of *Climate Warming* and *Climate Cooling* [above] as response variables) and compared their F-values, retaining variables with the highest F-value while excluding those with lower F-values from downstream analysis[73–75]. Following this step, eight morphological and life-history traits remained: body mass, brain-body ratio, tarsus length, bill length, egg mass, clutch size, incubation duration, and hand-wing index (see Germain et al.[76] for further description of relationships among key traits and Principal Component Analysis). These eight traits were then incorporated into further variable selection analyses using linear mixed-effect models to identify the best suite of traits likely to influence demographic responses during *Climate Warming* and *Climate Cooling* (Results and Discussion, main text).

**Supplementary Text S3: Contemporary latitudinal variation in key morphological/life-history traits**

The key morphological and life-history traits related to survival/growth, reproduction, and dispersal that we identified as being associated with species-specific $N_e$ responses to changing climate may also be reflected in how traits contribute to the contemporary distribution of species globally, given that the Earth's climate is currently undergoing a period of dramatic warming. Using a combination of trait data collected from museum specimens and the published literature for contemporary bird species distributed globally, we tested whether species currently distributed in tropical (i.e., warmer) latitudes express traits consistent with those species that exhibited increasing $N_e$ under the most recent period of *Climate Warming* (~147-123 kya), under the assumption that such traits may be indicative of a warm-adapted life-history.

We first constructed an additional series of PPA models aimed at identifying the trait network distinguishing species which exhibited increasing $N_e$ tendency under *Climate Warming* (n = 88 species) verses those with decreasing $N_e$ tendency (n = 127) during this period without limiting the confidence level. Note that this PPA measures only the response to climate warming, not climate warming and cooling as measured in the Warming Positive and Warming Negative PPA models. We also performed this analysis at different confidence levels. Details of best performing models are presented in table S12. Results from the best supported and final averaged models for this analysis are presented in figs. S11-20 (at differing confidence levels), and indicate that species which increased effective population size under *Climate Warming* were clearly differentiated from remaining species by lower HWI.

Next, we assembled morphological (i.e., body mass, bill length, and hand-wing index) and distributional (mean centroid longitude and latitude of breeding/resident range) data for all

10,950 contemporary bird species (example – fig. S23), again from datasets compiled for previous avian comparative studies[65–67]. In addition, estimates of egg mass for species not included in our previous analyses were collected from Rotenberry and Balasubramaniam[77], incubation duration from Cooney et al.[78], and clutch size from Jetz et al.[79], Werner and Griebler[80], and Cooney et al.[78]. From these combined sources, complete data were available for 2,745 species in total. Using a linear model with absolute latitude as the response and our six key traits as (scaled) predictor variables, we found that all traits were indeed significant predictors of current global distribution (fig. S24). Specifically, species currently centered at lower latitudes (i.e., warmer tropical regions) were found to have longer incubation durations and longer bills, but smaller clutch sizes, lighter eggs, lighter body mass, and lower hand-wing indexes.

Our results reveal some degree of concordance between the trait-mediated influences on demographic change under past climate warming and the contemporary distribution of species along a latitudinal gradient. Both sets of analyses revealed that less dispersive species appear better adapted (temporarily and spatially) to a 'warmer' life-history. Overall, these results from contemporary distributions conform to long-standing expectations (e.g., Bergmann's rule[81,82]) where larger bodied species are more likely to adapt to and therefore be found in cooler latitudes, and also highlight the relationship between migration ability and latitudinal variation, where temperate species are more likely to be migratory (and thus have higher HWI values) than tropical species[67]. Likewise, these global data also confirm predictions of larger clutch sizes at higher latitudes both within and across species[83–86], that parental investment towards offspring development (e.g., egg mass) is greater in cooler temperate latitudes[87–89], and that species at lower latitudes tend to exhibit longer incubation durations[78]. Thus, while these findings provide additional support that the six key traits identified through our analyses play a central role in mediating demographic responses to warming global temperatures across both space (i.e., latitudinal variation) and time (i.e., response to climate warming), they indicate that historical responses to climate change alone are not fully indicative of contemporary distributions.

**Table S1** – List of 325 species for which whole-genome sequencing data were constructed as part of the B10k Genomes Project Phase II (https://b10k.genomics.cn) by Feng et al.[19], including taxonomic order and assignment to one of the 11 major zoogeographic realms (where possible) identified by Holt et al.[28]. Species were assigned to the realm corresponding to their breeding range, and species present in multiple realms were assigned the realm corresponding to the sampling location. The 263 species which passed quality control checks for genome-wide coverage and missing data, as well as full demographic coverage over the focal time period (30 kya–1 mya) are identified as "Analyzed = YES", and demographic clustering groups (where k = 7 groups) are provided for each. *Warming* and *Cooling* (i.e. *Climate Warming* and *Climate Cooling* in-text) refer to demographic responses ("Increase", "Decrease", "Unrelated"; quantified by significant positive/negative correlations between $N_e$ and Global Average Surface Temperature) during periods of warming (~147–123kya) and cooling (~122–65kya, fig. S8). IUCN refers to current conservation status (LC = Least Concern, NT = Near Threatened, VU = Vulnerable, EN = Endangered, CR = Critically Endangered) according to the International Union for Conservation of Nature Red List (www.iucnredlist.org). The mean (±SD) of effective population size ($N_e$) estimates for each species from 30kya–1mya are given (x $10^4$).

| Species | Realm | Order | Cluster | Warming | Cooling | IUCN | Analyzed | Mean (SD) $N_e$ x $10^4$ |
|---|---|---|---|---|---|---|---|---|
| *Acanthisitta chloris* | Australian | Passeriformes | Group 4 | Increase | Increase | LC | YES | 12.14 (4.76) |
| *Acrocephalus arundinaceus* | Palearctic | Passeriformes | Group 7 | Increase | Decrease | LC | YES | 11.09 (3.73) |
| *Aegithalos caudatus* | Palearctic | Passeriformes | Group 6 | Decrease | Unrelated | LC | YES | 6.8 (1.1) |
| *Aegotheles bennettii* | Oceanian | Caprimulgiformes | Group 6 | Decrease | Decrease | LC | YES | 26.95 (11.31) |
| *Agapornis roseicollis* | Afrotropical | Psittaciformes | | | | LC | NO | 3.04 (0.34) |
| *Agelaius phoeniceus* | Nearctic | Passeriformes | Group 6 | Decrease | Decrease | LC | YES | 23.73 (14.67) |
| *Alaudala cheleensis* | Sino-Japanese | Passeriformes | Group 6 | Increase | Decrease | | YES | 63.14 (16.25) |
| *Alca torda* | Palearctic | Charadriiformes | Group 3 | Increase | Unrelated | NT | YES | 3.12 (0.53) |
| *Aleadryas rufinucha* | Oceanian | Passeriformes | Group 1 | Increase | Unrelated | LC | YES | 13.22 (2.66) |
| *Alectura lathami* | Australian | Galliformes | | | | LC | NO | 2.1 (0.46) |
| *Alopecoenas beccarii* | Oceanian | Columbiformes | Group 5 | Decrease | Decrease | LC | YES | 12.29 (1.66) |
| *Amazona guildingii* | Neotropical | Psittaciformes | Group 5 | Decrease | Decrease | VU | YES | 2.52 (1.58) |
| *Anas platyrhynchos* | | Anseriformes | Group 6 | Decrease | Decrease | LC | YES | 13.49 (7.5) |
| *Anas zonorhyncha* | | Anseriformes | Group 6 | Decrease | Decrease | LC | YES | 18.57 (8.28) |
| *Anhinga anhinga* | Nearctic | Suliformes | Group 6 | Decrease | Decrease | LC | YES | 5.34 (2.21) |
| *Anhinga rufa* | Afrotropical | Suliformes | | | | LC | NO | 2.94 (0.75) |
| *Anser cygnoides* | | Anseriformes | Group 6 | Decrease | Decrease | VU | YES | 8.73 (3.43) |
| *Anseranas semipalmata* | Australian | Anseriformes | Group 4 | Increase | Unrelated | LC | YES | 16.54 (4.53) |
| *Anthoscopus minutus* | Afrotropical | Passeriformes | Group 4 | Increase | Decrease | LC | YES | 11.07 (3.49) |
| *Antrostomus carolinensis* | Nearctic | Caprimulgiformes | Group 6 | Decrease | Decrease | NT | YES | 21.41 (7.02) |
| *Apaloderma vittatum* | Afrotropical | Trogoniformes | Group 4 | Unrelated | Increase | LC | YES | 7.42 (3.12) |
| *Aptenodytes forsteri* | | Sphenisciformes | Group 4 | Decrease | Decrease | NT | YES | 4.18 (0.59) |
| *Apteryx australis* | Australian | Apterygiformes | | | | VU | NO | 1.42 (0.44) |
| *Apteryx owenii* | Australian | Apterygiformes | | | | NT | NO | 1.25 (0.52) |
| *Apteryx rowi* | Australian | Apterygiformes | Group 5 | Increase | Increase | VU | YES | 1.83 (0.7) |
| *Aquila chrysaetos* | Nearctic | Accipitriformes | Group 7 | Increase | Decrease | LC | YES | 1.46 (0.25) |
| *Aramus guarauna* | Neotropical | Gruiformes | Group 4 | Increase | Unrelated | LC | YES | 48.84 (23.62) |
| *Ardeotis kori* | Afrotropical | Otidiformes | Group 5 | Decrease | Unrelated | NT | YES | 1.87 (0.56) |
| *Arenaria interpres* | Panamanian | Charadriiformes | Group 7 | Increase | Decrease | LC | YES | 21.8 (6.2) |

| Species | Realm | Order | Cluster | Warming | Cooling | IUCN | Analyzed | Mean (SD) $N_e$ x $10^4$ |
|---|---|---|---|---|---|---|---|---|
| *Asarcornis scutulata* | Nearctic | Anseriformes | | | | EN | NO | 2.91 (0.44) |
| *Atlantisia rogersi* | Palearctic | Gruiformes | | | | VU | NO | 1.14 (0.42) |
| *Atrichornis clamosus* | Australian | Passeriformes | Group 7 | Unrelated | Decrease | EN | YES | 9.95 (2.56) |
| *Balaeniceps rex* | Afrotropical | Pelecaniformes | | | | VU | NO | 0.34 (0.1) |
| *Balearica regulorum* | Afrotropical | Gruiformes | Group 7 | Decrease | Decrease | EN | YES | 3.53 (0.49) |
| *Bombycilla garrulus* | Nearctic | Passeriformes | Group 3 | Decrease | Increase | LC | YES | 17.73 (4.98) |
| *Brachypteracias leptosomus* | Madagascan | Coraciiformes | Group 5 | Decrease | Decrease | VU | YES | 3.49 (1.51) |
| *Bucco capensis* | Neotropical | Piciformes | Group 7 | Decrease | Decrease | LC | YES | 24.03 (10.92) |
| *Buceros rhinoceros* | Oriental | Bucerotiformes | | | | VU | NO | 0.82 (0.35) |
| *Bucorvus abyssinicus* | Afrotropical | Bucerotiformes | Group 6 | Decrease | Decrease | VU | YES | 3.25 (1.17) |
| *Buphagus erythrorhynchus* | Afrotropical | Passeriformes | Group 6 | Decrease | Decrease | LC | YES | 12.14 (4.97) |
| *Burhinus bistriatus* | Neotropical | Charadriiformes | Group 5 | Increase | Increase | LC | YES | 2.98 (0.69) |
| *Cairina moschata* | | Anseriformes | Group 7 | Decrease | Decrease | LC | YES | 12.78 (3.97) |
| *Calcarius ornatus* | Nearctic | Passeriformes | Group 5 | Decrease | Decrease | VU | YES | 29.71 (6.35) |
| *Callaeas wilsoni* | Australian | Passeriformes | Group 4 | Increase | Unrelated | NT | YES | 19.43 (9.86) |
| *Callipepla squamata* | Nearctic | Galliformes | Group 7 | Increase | Decrease | LC | YES | 11.24 (2.13) |
| *Calonectris borealis* | Palearctic | Procellariiformes | Group 4 | Decrease | Increase | LC | YES | 4.68 (0.83) |
| *Calypte anna* | Nearctic | Caprimulgiformes | Group 4 | Increase | Increase | LC | YES | 10.47 (3.14) |
| *Calyptomena viridis* | Oriental | Passeriformes | Group 6 | Decrease | Decrease | NT | YES | 24.47 (10.54) |
| *Campylorhamphus procurvoides* | Neotropical | Passeriformes | Group 4 | Increase | Decrease | LC | YES | 7.41 (2.03) |
| *Cardinalis cardinalis* | Nearctic | Passeriformes | Group 4 | Decrease | Increase | LC | YES | 11.62 (4.78) |
| *Cariama cristata* | Neotropical | Cariamiformes | Group 6 | Decrease | Decrease | LC | YES | 4.11 (1.17) |
| *Casuarius casuarius* | Oceanian | Casuariiformes | Group 7 | Increase | Unrelated | LC | YES | 2.8 (0.71) |
| *Cathartes aura* | Nearctic | Cathartiformes | Group 7 | Unrelated | Decrease | LC | YES | 5.03 (0.67) |
| *Catharus fuscescens* | Nearctic | Passeriformes | Group 1 | Increase | Increase | LC | YES | 34.47 (10.83) |
| *Centropus unirufus* | Oriental | Cuculiformes | Group 2 | Increase | Decrease | NT | YES | 12.96 (3.78) |
| *Cephalopterus ornatus* | Neotropical | Passeriformes | Group 4 | Increase | Unrelated | LC | YES | 5.95 (2.48) |
| *Cepphus grylle* | Nearctic | Charadriiformes | Group 3 | Unrelated | Unrelated | LC | YES | 4.65 (0.89) |
| *Cercotrichas coryphaeus* | Afrotropical | Passeriformes | Group 6 | Decrease | Decrease | LC | YES | 42.14 (16.4) |

| Species | Realm | Order | Cluster | Warming | Cooling | IUCN | Analyzed | Mean (SD) $N_e$ x $10^4$ |
|---|---|---|---|---|---|---|---|---|
| *Certhia brachydactyla* | Palearctic | Passeriformes | | | | LC | NO | 6.44 (0.41) |
| *Certhia familiaris* | Palearctic | Passeriformes | Group 7 | Decrease | Decrease | LC | YES | 6.63 (1.31) |
| *Ceuthmochares aereus* | Afrotropical | Cuculiformes | Group 6 | Decrease | Decrease | LC | YES | 11.8 (4.07) |
| *Ceyx cyanopectus* | Oriental | Coraciiformes | | | | LC | NO | |
| *Chaetorhynchus papuensis* | Oceanian | Passeriformes | Group 7 | Decrease | Decrease | LC | YES | 38.03 (9.39) |
| *Chaetura pelagica* | Nearctic | Caprimulgiformes | | | | VU | NO | 3.34 (0.33) |
| *Charadrius vociferus* | Neotropical | Charadriiformes | | | | LC | NO | 4.02 (1.52) |
| *Chauna torquata* | Neotropical | Anseriformes | Group 6 | Decrease | Decrease | LC | YES | 6.74 (2.3) |
| *Chionis minor* | Afrotropical | Charadriiformes | | | | LC | NO | 0.68 (0.1) |
| *Chlamydotis macqueenii* | Saharo-Arabian | Otidiformes | Group 3 | Increase | Unrelated | VU | YES | 6.05 (1.45) |
| *Chloroceryle aenea* | Neotropical | Coraciiformes | Group 5 | Decrease | Decrease | LC | YES | 18.99 (6.36) |
| *Chloropsis cyanopogon* | Oriental | Passeriformes | Group 6 | Decrease | Decrease | NT | YES | 20.07 (12.06) |
| *Chloropsis hardwickii* | Sino-Japanese | Passeriformes | Group 3 | Decrease | Decrease | LC | YES | 9.6 (2.65) |
| *Chordeiles acutipennis* | Neotropical | Caprimulgiformes | Group 6 | Decrease | Decrease | LC | YES | 10.62 (2.17) |
| *Chroicocephalus maculipennis* | Neotropical | Charadriiformes | Group 4 | Increase | Increase | LC | YES | 7.65 (3.26) |
| *Chunga burmeisteri* | Neotropical | Cariamiformes | Group 2 | Increase | Decrease | LC | YES | 12.39 (4.87) |
| *Ciccaba nigrolineata* | Panamanian | Strigiformes | Group 5 | Decrease | Increase | LC | YES | 2.87 (1.07) |
| *Ciconia maguari* | Neotropical | Ciconiiformes | Group 1 | Increase | Increase | LC | YES | 1.43 (0.26) |
| *Cinclus mexicanus* | Nearctic | Passeriformes | | | | LC | NO | 3.16 (0.67) |
| *Circaetus pectoralis* | Afrotropical | Accipitriformes | Group 1 | Increase | Increase | LC | YES | 2.21 (0.16) |
| *Cisticola juncidis* | Sino-Japanese | Passeriformes | | | | LC | NO | 6.06 (2.18) |
| *Climacteris rufus* | Australian | Passeriformes | Group 4 | Increase | Increase | LC | YES | 10.83 (3.49) |
| *Cnemophilus loriae* | Oceanian | Passeriformes | Group 2 | Decrease | Decrease | LC | YES | 16.8 (6.74) |
| *Cochlearius cochlearius* | Neotropical | Pelecaniformes | Group 7 | Increase | Decrease | LC | YES | 32.03 (7.11) |
| *Colinus virginianus* | Nearctic | Galliformes | Group 6 | Decrease | Decrease | NT | YES | 25.03 (9.48) |
| *Colius striatus* | Afrotropical | Coliiformes | Group 5 | Unrelated | Increase | LC | YES | 8.36 (1.85) |
| *Columba livia* | Palearctic | Columbiformes | Group 7 | Decrease | Decrease | LC | YES | 12.63 (2.79) |
| *Columbina picui* | Neotropical | Columbiformes | Group 3 | Unrelated | Increase | LC | YES | 45.01 (11.38) |
| *Copsychus sechellarum* | Oceanian | Passeriformes | | | | EN | NO | 2.24 (0.73) |

| Species | Realm | Order | Cluster | Warming | Cooling | IUCN | Analyzed | Mean (SD) $N_e$ x $10^4$ |
|---|---|---|---|---|---|---|---|---|
| *Corvus brachyrhynchos* | Nearctic | Passeriformes | | | | LC | NO | 1.56 (1.27) |
| *Corvus cornix* | Palearctic | Passeriformes | Group 3 | Decrease | Decrease | LC | YES | 5.98 (3.28) |
| *Corvus moneduloides* | Oceanian | Passeriformes | | | | LC | NO | 1.49 (0.67) |
| *Corythaeola cristata* | Afrotropical | Musophagiformes | Group 3 | Increase | Increase | LC | YES | 4.95 (1.51) |
| *Corythaixoides concolor* | Afrotropical | Musophagiformes | Group 5 | Increase | Increase | LC | YES | 4.81 (0.88) |
| *Coturnix japonica* | Sino-Japanese | Galliformes | Group 6 | Decrease | Decrease | NT | YES | 79.27 (64.14) |
| *Crotophaga sulcirostris* | Panamanian | Cuculiformes | Group 4 | Increase | Unrelated | LC | YES | 18.2 (8.03) |
| *Crypturellus cinnamomeus* | Panamanian | Tinamiformes | Group 6 | Decrease | Decrease | LC | YES | 6.92 (2.66) |
| *Crypturellus soui* | Neotropical | Tinamiformes | Group 6 | Decrease | Decrease | LC | YES | 12.83 (5.43) |
| *Crypturellus undulatus* | Neotropical | Tinamiformes | Group 2 | Decrease | Decrease | LC | YES | 27.86 (14.92) |
| *Cuculus canorus* | Palearctic | Cuculiformes | Group 6 | Decrease | Decrease | LC | YES | 5.58 (1.29) |
| *Daphoenositta chrysoptera* | Australian | Passeriformes | Group 1 | Increase | Unrelated | LC | YES | 50.43 (5.49) |
| *Dasyornis broadbenti* | Australian | Passeriformes | Group 7 | Decrease | Decrease | LC | YES | 7.55 (3.05) |
| *Dicaeum eximium* | Oceanian | Passeriformes | Group 4 | Decrease | Increase | LC | YES | 26.44 (14.61) |
| *Dicrurus megarhynchus* | Oceanian | Passeriformes | | | | NT | NO | 5.52 (3.78) |
| *Donacobius atricapilla* | Neotropical | Passeriformes | Group 6 | Decrease | Decrease | LC | YES | 6.61 (2.05) |
| *Dromaius novaehollandiae* | Australian | Casuariiformes | Group 7 | Decrease | Unrelated | LC | YES | 3.92 (0.99) |
| *Dromas ardeola* | Afrotropical | Charadriiformes | Group 5 | Decrease | Decrease | LC | YES | 2.15 (0.85) |
| *Drymodes brunneopygia* | Australian | Passeriformes | Group 5 | Increase | Decrease | LC | YES | 6.8 (1.44) |
| *Dryoscopus gambensis* | Afrotropical | Passeriformes | Group 1 | Increase | Increase | LC | YES | 7.95 (0.69) |
| *Dyaphorophyia castanea* | Afrotropical | Passeriformes | Group 1 | Increase | Increase | LC | YES | 48.98 (76.27) |
| *Edolisoma coerulescens* | Oriental | Passeriformes | Group 7 | Decrease | Increase | LC | YES | 14.49 (1.27) |
| *Egretta garzetta* | Sino-Japanese | Pelecaniformes | Group 5 | Decrease | Decrease | LC | YES | 4.16 (0.34) |
| *Emberiza fucata* | Sino-Japanese | Passeriformes | Group 2 | Unrelated | Decrease | LC | YES | 21.72 (16.58) |
| *Erpornis zantholeuca* | Oriental | Passeriformes | Group 6 | Decrease | Decrease | LC | YES | 12.34 (7.66) |
| *Erythrocercus mccallii* | Afrotropical | Passeriformes | Group 3 | Decrease | Unrelated | LC | YES | 14.21 (3.7) |
| *Eubucco bourcierii* | Panamanian | Piciformes | | | | LC | NO | 5.29 (1.83) |
| *Eudromia elegans* | Neotropical | Tinamiformes | Group 7 | Decrease | Decrease | LC | YES | 8.54 (2.23) |

| Species | Realm | Order | Cluster | Warming | Cooling | IUCN | Analyzed | Mean (SD) $N_e$ x $10^4$ |
|---|---|---|---|---|---|---|---|---|
| *Eulacestoma nigropectus* | Oceanian | Passeriformes | Group 1 | Decrease | Decrease | LC | YES | 8.02 (0.46) |
| *Eurypyga helias* | Neotropical | Eurypygiformes | Group 7 | Increase | Decrease | LC | YES | 11.66 (2.42) |
| *Eurystomus gularis* | Afrotropical | Coraciiformes | Group 2 | Increase | Decrease | LC | YES | 16.29 (2.17) |
| *Falco cherrug* | Saharo-Arabian | Falconiformes | Group 1 | Decrease | Unrelated | EN | YES | 3.1 (0.94) |
| *Falco peregrinus* | Saharo-Arabian | Falconiformes | | | | LC | NO | 1.08 (0.05) |
| *Falcunculus frontatus* | Australian | Passeriformes | Group 3 | Decrease | Increase | LC | YES | 14.94 (4.09) |
| *Formicarius rufipectus* | Panamanian | Passeriformes | | | | LC | NO | 4.21 (1.82) |
| *Fregata magnificens* | Nearctic | Suliformes | | | | LC | NO | 0.78 (0.06) |
| *Fregetta grallaria* | Oceanian | Procellariiformes | Group 3 | Unrelated | Decrease | LC | YES | 14.15 (6.58) |
| *Fulmarus glacialis* | Palearctic | Procellariiformes | Group 3 | Increase | Increase | LC | YES | 2.14 (0.2) |
| *Furnarius figulus* | Neotropical | Passeriformes | Group 4 | Increase | Decrease | LC | YES | 6.66 (2.58) |
| *Galbula dea* | Neotropical | Piciformes | Group 4 | Increase | Unrelated | LC | YES | 11.22 (4.69) |
| *Gallus gallus* | Oriental | Galliformes | | | | LC | NO | 12.36 (6.29) |
| *Gavia stellata* | Palearctic | Gaviiformes | Group 7 | Unrelated | Decrease | LC | YES | 6.58 (1.09) |
| *Geococcyx californianus* | Nearctic | Cuculiformes | | | | LC | NO | 5.37 (1.29) |
| *Geospiza fortis* | Neotropical | Passeriformes | | | | LC | NO | 1.98 (0.34) |
| *Glareola pratincola* | Saharo-Arabian | Charadriiformes | Group 1 | Increase | Increase | LC | YES | 20.29 (3.73) |
| *Glaucidium brasilianum* | Neotropical | Strigiformes | Group 1 | Increase | Increase | LC | YES | 15.11 (13.93) |
| *Grallaria varia* | Neotropical | Passeriformes | Group 4 | Decrease | Increase | LC | YES | 7.17 (2.81) |
| *Grantiella picta* | Australian | Passeriformes | Group 7 | Increase | Increase | VU | YES | 19.62 (5.21) |
| *Grus americana* | Nearctic | Gruiformes | | | | EN | NO | 1.37 (0.28) |
| *Gymnorhina tibicen* | Australian | Passeriformes | Group 3 | Decrease | Increase | LC | YES | 22.01 (7.75) |
| *Halcyon senegalensis* | Afrotropical | Coraciiformes | Group 5 | Decrease | Decrease | LC | YES | 20.75 (8.9) |
| *Haliaeetus albicilla* | Palearctic | Accipitriformes | Group 4 | Increase | Unrelated | LC | YES | 0.93 (0.28) |
| *Haliaeetus leucocephalus* | Nearctic | Accipitriformes | | | | LC | NO | |
| *Heliornis fulica* | Neotropical | Gruiformes | Group 5 | Unrelated | Increase | LC | YES | 8.91 (4.46) |
| *Hemignathus wilsoni* | Nearctic | Passeriformes | | | | EN | NO | 4.09 (1.6) |
| *Hemiprocne comata* | Oriental | Caprimulgiformes | Group 6 | Decrease | Decrease | LC | YES | 20.8 (6.94) |

| Species | Realm | Order | Cluster | Warming | Cooling | IUCN | Analyzed | Mean (SD) $N_e$ x $10^4$ |
|---|---|---|---|---|---|---|---|---|
| *Herpetotheres cachinnans* | Panamanian | Falconiformes | Group 6 | Decrease | Decrease | LC | YES | 8.5 (2.39) |
| *Himantopus himantopus* | Neotropical | Charadriiformes | Group 4 | Increase | Increase | LC | YES | 13.75 (4.38) |
| *Hippolais icterina* | Palearctic | Passeriformes | Group 1 | Increase | Increase | LC | YES | 16.09 (3.5) |
| *Hirundo rustica* | Nearctic | Passeriformes | Group 3 | Decrease | Decrease | LC | YES | 26.49 (14.85) |
| *Horornis vulcanius* | Oriental | Passeriformes | | | | | NO | 6.29 (1.54) |
| *Hydrobates tethys* | Nearctic | Procellariiformes | Group 6 | Decrease | Decrease | LC | YES | 14.18 (3.24) |
| *Hylia prasina* | Afrotropical | Passeriformes | Group 1 | Increase | Increase | LC | YES | 26.98 (16.97) |
| *Hypocryptadius cinnamomeus* | Oriental | Passeriformes | | | | LC | NO | 5.89 (1.65) |
| *Ibidorhyncha struthersii* | Sino-Japanese | Charadriiformes | | | | LC | NO | 1.59 (0.18) |
| *Ifrita kowaldi* | Oceanian | Passeriformes | Group 4 | Decrease | Unrelated | LC | YES | 10.26 (2.68) |
| *Illadopsis cleaveri* | Afrotropical | Passeriformes | Group 3 | Unrelated | Decrease | LC | YES | 48.98 (26.59) |
| *Indicator maculatus* | Afrotropical | Piciformes | Group 7 | Decrease | Decrease | LC | YES | 14.24 (3.37) |
| *Jacana jacana* | Panamanian | Charadriiformes | Group 6 | Decrease | Decrease | LC | YES | 11.92 (2.86) |
| *Lanius ludovicianus* | Nearctic | Passeriformes | | | | NT | NO | 6.12 (4.62) |
| *Larus smithsonianus* | Nearctic | Charadriiformes | Group 7 | Unrelated | Decrease | LC | YES | 3.53 (0.57) |
| *Leiothrix lutea* | Sino-Japanese | Passeriformes | Group 6 | Decrease | Decrease | LC | YES | 17.23 (7.62) |
| *Lepidothrix coronata* | Neotropical | Passeriformes | Group 2 | Decrease | Decrease | LC | YES | 51.28 (35.51) |
| *Leptocoma aspasia* | Oceanian | Passeriformes | Group 4 | Increase | Increase | LC | YES | 13.68 (11.7) |
| *Leptosomus discolor* | Madagascan | Leptosomiformes | Group 5 | Increase | Increase | LC | YES | 12.43 (1.9) |
| *Leucopsar rothschildi* | Oriental | Passeriformes | | | | CR | NO | 0.75 (0.19) |
| *Limosa lapponica* | Australian | Charadriiformes | Group 1 | Decrease | Increase | NT | YES | 6.27 (2.1) |
| *Locustella ochotensis* | Palearctic | Passeriformes | Group 2 | Increase | Unrelated | LC | YES | 19.04 (4.96) |
| *Lonchura striata* | Oriental | Passeriformes | Group 2 | Increase | Decrease | LC | YES | 21.43 (16.37) |
| *Lophotis ruficrista* | Afrotropical | Otidiformes | Group 5 | Decrease | Unrelated | LC | YES | 6.08 (1.96) |
| *Loxia curvirostra* | Nearctic | Passeriformes | Group 4 | Decrease | Increase | LC | YES | 8.34 (3.39) |
| *Loxia leucoptera* | Nearctic | Passeriformes | Group 4 | Decrease | Increase | LC | YES | 35.91 (23.13) |
| *Machaerirhynchus nigripectus* | Oceanian | Passeriformes | Group 3 | Increase | Unrelated | LC | YES | 31.55 (9.34) |
| *Malurus elegans* | Australian | Passeriformes | Group 3 | Increase | Increase | LC | YES | 6.77 (2.6) |
| *Manacus manacus* | Panamanian | Passeriformes | Group 7 | Increase | Increase | LC | YES | 10.46 (2.13) |

| Species | Realm | Order | Cluster | Warming | Cooling | IUCN | Analyzed | Mean (SD) $N_e$ x $10^4$ |
|---|---|---|---|---|---|---|---|---|
| *Melanocharis versteri* | Oceanian | Passeriformes | Group 3 | Decrease | Decrease | LC | YES | 37.28 (14.14) |
| *Melopsittacus undulatus* | Australian | Psittaciformes | Group 1 | Increase | Increase | LC | YES | 6.58 (1.49) |
| *Melospiza melodia* | Nearctic | Passeriformes | Group 4 | Increase | Decrease | LC | YES | 12.89 (6) |
| *Menura novaehollandiae* | Australian | Passeriformes | Group 7 | Increase | Decrease | LC | YES | 3.43 (0.76) |
| *Merops nubicus* | Afrotropical | Coraciiformes | Group 5 | Increase | Increase | LC | YES | 12.36 (4.45) |
| *Mesembrinibis cayennensis* | Panamanian | Pelecaniformes | Group 6 | Decrease | Decrease | LC | YES | 9.48 (3.68) |
| *Mesitornis unicolor* | Madagascan | Mesitornithiformes | Group 5 | Decrease | Increase | VU | YES | 3.63 (0.37) |
| *Mionectes macconnelli* | Neotropical | Passeriformes | Group 5 | Increase | Decrease | LC | YES | 13.08 (4.83) |
| *Mohoua ochrocephala* | Australian | Passeriformes | Group 7 | Increase | Decrease | EN | YES | 9.83 (3.4) |
| *Molothrus ater* | Nearctic | Passeriformes | Group 3 | Decrease | Decrease | LC | YES | 16.35 (9.42) |
| *Motacilla alba* | Palearctic | Passeriformes | Group 2 | Decrease | Decrease | LC | YES | 13.56 (7.82) |
| *Mystacornis crossleyi* | Madagascan | Passeriformes | Group 6 | Decrease | Decrease | LC | YES | 8.6 (3.64) |
| *Neodrepanis coruscans* | Madagascan | Passeriformes | | | | LC | NO | 6.29 (2.1) |
| *Neopipo cinnamomea* | Neotropical | Passeriformes | Group 5 | Decrease | Decrease | LC | YES | 7.17 (2.71) |
| *Nesospiza acunhae* | Palearctic | Passeriformes | | | | VU | NO | 0.67 (0.08) |
| *Nestor notabilis* | Australian | Psittaciformes | Group 5 | Decrease | Unrelated | EN | YES | 1.9 (0.25) |
| *Nicator chloris* | Afrotropical | Passeriformes | Group 7 | Decrease | Decrease | LC | YES | 10.53 (3.34) |
| *Nothocercus julius* | Neotropical | Tinamiformes | Group 7 | Decrease | Decrease | LC | YES | 4.73 (0.92) |
| *Nothocercus nigrocapillus* | Neotropical | Struthioniformes | Group 3 | Increase | Increase | LC | YES | 8.52 (2.01) |
| *Nothoprocta ornata* | Neotropical | Struthioniformes | Group 5 | Unrelated | Decrease | LC | YES | 12.18 (2.24) |
| *Nothoprocta pentlandii* | Neotropical | Struthioniformes | Group 7 | Decrease | Decrease | LC | YES | 16.71 (4.69) |
| *Nothoprocta perdicaria* | Neotropical | Struthioniformes | Group 3 | Increase | Increase | LC | YES | 16.06 (8.76) |
| *Notiomystis cincta* | Australian | Passeriformes | | | | VU | NO | 7.86 (2.83) |
| *Numida meleagris* | Afrotropical | Galliformes | Group 7 | Increase | Decrease | LC | YES | 15.61 (3.5) |
| *Nyctibius bracteatus* | Neotropical | Caprimulgiformes | Group 5 | Decrease | Decrease | LC | YES | 15.76 (4.09) |
| *Nyctibius grandis* | Neotropical | Caprimulgiformes | Group 5 | Decrease | Increase | LC | YES | 6.75 (1.79) |
| *Nycticryphes semicollaris* | Neotropical | Charadriiformes | Group 1 | Increase | Increase | LC | YES | 8.67 (3.24) |
| *Nyctiprogne leucopyga* | Neotropical | Caprimulgiformes | Group 5 | Decrease | Increase | LC | YES | 30.5 (8.88) |
| *Oceanites oceanicus* | Nearctic | Procellariiformes | Group 5 | Decrease | Increase | LC | YES | 4.59 (1.43) |

| Species | Realm | Order | Cluster | Warming | Cooling | IUCN | Analyzed | Mean (SD) $N_e$ x $10^4$ | |
|---|---|---|---|---|---|---|---|---|---|
| *Odontophorus gujanensis* | Neotropical | Galliformes | Group 2 | Decrease | Decrease | NT | YES | 25.11 (23.23) | |
| *Oenanthe oenanthe* | Palearctic | Passeriformes | Group 1 | Increase | Increase | LC | YES | 19.71 (14.89) | |
| *Onychorhynchus coronatus* | Panamanian | Passeriformes | | | | LC | NO | 5.21 (2.29) | |
| *Opisthocomus hoazin* | Neotropical | Opisthocomiformes | | | LC | NO | 0.23 (0.01) | | |
| *Oreocharis arfaki* | Oceanian | Passeriformes | | Group 4 | Decrease | Increase | LC | YES | 62.32 ( |
| *Origma solitaria* | Australian | Passeriformes | Group 4 | Increase | Increase | LC | YES | 7.39 (3.8) | |
| *Oriolus oriolus* | Palearctic | Passeriformes | Group 3 | Decrease | Decrease | LC | YES | 19.19 (11.47) | |
| *Orthonyx spaldingii* | Australian | Passeriformes | Group 5 | Increase | Decrease | LC | YES | 4.77 (1.1) | |
| *Oxyruncus cristatus* | Neotropical | Passeriformes | Group 3 | Decrease | Decrease | LC | YES | 34.37 (24.26) | |
| *Pachycephala philippinensis* | Oriental | Passeriformes | Group 1 | Increase | Increase | LC | YES | 16.7 (5.62) | |
| *Pachyramphus minor* | Neotropical | Passeriformes | Group 7 | Decrease | Decrease | LC | YES | 50.29 (16.55) | |
| *Pandion haliaetus* | Nearctic | Accipitriformes | | | | LC | NO | 1.08 (0.18) | |
| *Panurus biarmicus* | Palearctic | Passeriformes | | | | LC | NO | 7.54 (2.55) | |
| *Paradisaea raggiana* | Oceanian | Passeriformes | Group 7 | Decrease | Decrease | LC | YES | 6.42 (1.83) | |
| *Pardalotus punctatus* | Australian | Passeriformes | Group 3 | Decrease | Increase | LC | YES | 56.96 (23.48) | |
| *Parus major* | Palearctic | Passeriformes | Group 4 | Increase | Increase | LC | YES | 7.23 (3.1) | |
| *Passer domesticus* | Palearctic | Passeriformes | Group 7 | Unrelated | Decrease | LC | YES | 14.85 (5.63) | |
| *Passerina amoena* | Nearctic | Passeriformes | Group 6 | Decrease | Decrease | LC | YES | 42.64 (26.2) | |
| *Patagioenas fasciata* | | Columbiformes | Group 2 | Increase | Decrease | LC | YES | 13.3 (6.3) | |
| *Pedionomus torquatus* | Australian | Charadriiformes | Group 4 | Increase | Increase | CR | YES | 16.63 (9.04) | |
| *Pelecanoides urinatrix* | Neotropical | Procellariiformes | Group 6 | Decrease | Decrease | LC | YES | 10.1 (3.37) | |
| *Pelecanus crispus* | Saharo-Arabian | Pelecaniformes | | | | NT | NO | 0.8 (0.05) | |
| *Penelope pileata* | Neotropical | Galliformes | Group 5 | Increase | Increase | VU | YES | 6.07 (1.61) | |
| *Peucedramus taeniatus* | Nearctic | Passeriformes | Group 3 | Unrelated | Increase | LC | YES | 24.53 (5.23) | |
| *Phaethon lepturus* | Afrotropical | Phaethontiformes | Group 6 | Decrease | Decrease | LC | YES | 3.77 (1.52) | |
| *Phalacrocorax auritus* | Nearctic | Suliformes | Group 2 | Increase | Unrelated | LC | YES | 2.68 (0.55) | |
| *Phalacrocorax brasilianus* | Neotropical | Suliformes | Group 7 | Increase | Unrelated | LC | YES | 9.63 (1.17) | |
| *Phalacrocorax carbo* | Palearctic | Suliformes | Group 3 | Decrease | Decrease | LC | YES | 2.53 (0.39) | |
| *Phalacrocorax harrisi* | Neotropical | Suliformes | | | | VU | NO | 0.48 (0.05) | |

| Species | Realm | Order | Cluster | Warming | Cooling | IUCN | Analyzed | Mean (SD) $N_e$ x $10^4$ |
|---|---|---|---|---|---|---|---|---|
| *Phalacrocorax pelagicus* | Nearctic | Suliformes | Group 7 | Increase | Decrease | LC | YES | 2.5 (0.29) |
| *Phasianus colchicus* | Sino-Japanese | Galliformes | Group 3 | Increase | Increase | LC | YES | 24.11 (8.62) |
| *Pheucticus melanocephalus* | Nearctic | Passeriformes | Group 2 | Decrease | Decrease | LC | YES | 19.45 (8.48) |
| *Phoenicopterus ruber* | | Phoenicopteriformes | Group 7 | Decrease | Decrease | LC | YES | 8.17 (1.75) |
| *Phylloscopus trochilus* | Palearctic | Passeriformes | Group 7 | Increase | Unrelated | LC | YES | 25.23 (8.82) |
| *Piaya cayana* | Neotropical | Cuculiformes | Group 6 | Decrease | Decrease | LC | YES | 17.63 (7.13) |
| *Picathartes gymnocephalus* | Afrotropical | Passeriformes | | | | VU | NO | 1.69 (0.15) |
| *Picoides pubescens* | Nearctic | Piciformes | | | | LC | NO | 4.99 (1.19) |
| *Piprites chloris* | Neotropical | Passeriformes | Group 1 | Increase | Increase | LC | YES | 32.94 (59.32) |
| *Pitta sordida* | Oriental | Passeriformes | Group 6 | Decrease | Decrease | LC | YES | 12.81 (5.38) |
| *Ploceus nigricollis* | Afrotropical | Passeriformes | Group 3 | Decrease | Decrease | LC | YES | 36.23 (14.06) |
| *Pluvianellus socialis* | Neotropical | Charadriiformes | | | | NT | NO | 1.54 (0.34) |
| *Podargus strigoides* | Australian | Caprimulgiformes | Group 6 | Decrease | Decrease | LC | YES | 19.63 (6.59) |
| *Podiceps cristatus* | Nearctic | Podicipediformes | Group 3 | Decrease | Decrease | LC | YES | 5.62 (1.24) |
| *Podilymbus podiceps* | Neotropical | Podicipediformes | Group 4 | Increase | Increase | LC | YES | 10.06 (2.97) |
| *Poecile atricapillus* | Nearctic | Passeriformes | Group 2 | Increase | Decrease | LC | YES | 20.99 (8.28) |
| *Polioptila caerulea* | Nearctic | Passeriformes | Group 1 | Decrease | Increase | LC | YES | 63.55 (25.38) |
| *Pomatorhinus ruficollis* | Sino-Japanese | Passeriformes | Group 7 | Increase | Decrease | LC | YES | 31 (13.92) |
| *Pomatostomus ruficeps* | Australian | Passeriformes | Group 7 | Increase | Decrease | LC | YES | 7 (1.13) |
| *Promerops cafer* | Afrotropical | Passeriformes | Group 3 | Decrease | Decrease | LC | YES | 10.75 (3.23) |
| *Prunella fulvescens* | Palearctic | Passeriformes | Group 4 | Increase | Increase | LC | YES | 7.87 (1.16) |
| *Prunella himalayana* | Palearctic | Passeriformes | Group 3 | Decrease | Increase | LC | YES | 35.7 (14.15) |
| *Psilopogon haemacephalus* | Oriental | Piciformes | | | | LC | NO | 5.96 (4.05) |
| *Psophia crepitans* | Neotropical | Gruiformes | Group 2 | Increase | Decrease | NT | YES | 17.35 (6.43) |
| *Pterocles burchelli* | Afrotropical | Pterocliformes | Group 5 | Decrease | Unrelated | LC | YES | 11.78 (8.54) |
| *Pterocles gutturalis* | Afrotropical | Pterocliformes | Group 1 | Decrease | Increase | LC | YES | 4.84 (0.94) |
| *Pteruthius melanotis* | Sino-Japanese | Passeriformes | Group 3 | Unrelated | Decrease | LC | YES | 16.02 (5.76) |
| *Ptilonorhynchus violaceus* | Australian | Passeriformes | Group 2 | Increase | Decrease | LC | YES | 2.94 (0.6) |
| *Ptilorrhoa leucosticta* | Oceanian | Passeriformes | Group 7 | Unrelated | Decrease | LC | YES | 22.93 (7.47) |

| Species | Realm | Order | Cluster | Warming | Cooling | IUCN | Analyzed | Mean (SD) $N_e$ x $10^4$ |
|---|---|---|---|---|---|---|---|---|
| *Pycnonotus jocosus* | Nearctic | Passeriformes | Group 1 | Increase | Increase | LC | YES | 17.81 (2.22) |
| *Pygoscelis adeliae* | | Sphenisciformes | Group 4 | Increase | Increase | LC | YES | 2.88 (0.94) |
| *Quiscalus mexicanus* | Panamanian | Passeriformes | | | | LC | NO | 4.97 (1.96) |
| *Ramphastos sulfuratus* | Panamanian | Piciformes | Group 5 | Unrelated | Increase | LC | YES | 5.8 (1.09) |
| *Regulus satrapa* | Nearctic | Passeriformes | Group 3 | Decrease | Increase | LC | YES | 25.2 (8.68) |
| *Rhabdornis inornatus* | Oriental | Passeriformes | Group 1 | Decrease | Increase | LC | YES | 5.84 (0.32) |
| *Rhadina sibilatrix* | Palearctic | Passeriformes | Group 1 | Increase | Increase | LC | YES | 55.92 (25.37) |
| *Rhagologus leucostigma* | Oceanian | Passeriformes | Group 3 | Decrease | Decrease | LC | YES | 20.68 (6.62) |
| *Rhea americana* | Neotropical | Rheiformes | Group 4 | Increase | Increase | NT | YES | 5.61 (2.07) |
| *Rhea pennata* | Neotropical | Rheiformes | Group 3 | Increase | Decrease | LC | YES | 3.01 (0.72) |
| *Rhinopomastus cyanomelas* | Afrotropical | Bucerotiformes | Group 6 | Decrease | Decrease | LC | YES | 6.66 (3.08) |
| *Rhinoptilus africanus* | Afrotropical | Charadriiformes | Group 4 | Increase | Increase | LC | YES | 21.35 (5.37) |
| *Rhipidura dahli* | Oceanian | Passeriformes | | | | LC | NO | 4.29 (3.45) |
| *Rhodinocichla rosea* | Panamanian | Passeriformes | | | | LC | NO | 4.03 (0.94) |
| *Rhynochetos jubatus* | Oceanian | Eurypygiformes | Group 3 | Decrease | Decrease | EN | YES | 9.16 (2.45) |
| *Rissa tridactyla* | Nearctic | Charadriiformes | Group 6 | Decrease | Decrease | VU | YES | 4.19 (2.04) |
| *Rostratula benghalensis* | Oriental | Charadriiformes | Group 7 | Decrease | Increase | LC | YES | 4.54 (0.49) |
| *Rynchops niger* | Neotropical | Charadriiformes | Group 3 | Unrelated | Increase | LC | YES | 5.9 (1.39) |
| *Sakesphorus luctuosus* | Neotropical | Passeriformes | Group 7 | Decrease | Decrease | LC | YES | 7 (1.18) |
| *Sapayoa aenigma* | Panamanian | Passeriformes | Group 4 | Increase | Increase | LC | YES | 7.4 (3.92) |
| *Sclerurus mexicanus* | Neotropical | Passeriformes | Group 4 | Increase | Increase | LC | YES | 7.21 (2.17) |
| *Scopus umbretta* | Afrotropical | Pelecaniformes | | | | LC | NO | 2.88 (0.24) |
| *Scytalopus superciliaris* | Neotropical | Passeriformes | Group 3 | Decrease | Decrease | LC | YES | 8.46 (2.09) |
| *Serilophus lunatus* | Oriental | Passeriformes | | | | LC | NO | 4.32 (0.69) |
| *Setophaga coronata* | Nearctic | Passeriformes | Group 7 | Increase | Increase | LC | YES | 14.59 (4.28) |
| *Setophaga kirtlandii* | Nearctic | Passeriformes | Group 1 | Increase | Increase | NT | YES | 14.98 (9.28) |
| *Sinosuthora webbiana* | Palearctic | Passeriformes | Group 7 | Increase | Unrelated | LC | YES | 20.15 (5.75) |
| *Sitta europaea* | Palearctic | Passeriformes | | | | LC | NO | 3.38 (0.87) |
| *Smithornis capensis* | Afrotropical | Passeriformes | Group 7 | Increase | Unrelated | LC | YES | 11.49 (3.29) |

| Species | Realm | Order | Cluster | Warming | Cooling | IUCN | Analyzed | Mean (SD) $N_e$ x $10^4$ |
|---|---|---|---|---|---|---|---|---|
| *Spizaetus tyrannus* | Neotropical | Accipitriformes | Group 1 | Decrease | Increase | LC | YES | 2.1 (0.51) |
| *Spizella passerina* | Nearctic | Passeriformes | | | | LC | NO | 7.17 (1.3) |
| *Stercorarius parasiticus* | Panamanian | Charadriiformes | Group 1 | Decrease | Increase | LC | YES | 4.09 (2.55) |
| *Sterrhoptilus dennistouni* | Oriental | Passeriformes | Group 4 | Increase | Increase | NT | YES | 8.27 (2.63) |
| *Strix occidentalis* | Nearctic | Strigiformes | | | | NT | NO | 0.66 (0.19) |
| *Struthidea cinerea* | Australian | Passeriformes | Group 6 | Decrease | Decrease | LC | YES | 15.28 (7.21) |
| *Struthio camelus* | Afrotropical | Struthioniformes | | | | LC | NO | 0.49 (0.21) |
| *Sturnus vulgaris* | | Passeriformes | Group 2 | Increase | Decrease | LC | YES | 7.48 (2.87) |
| *Sylvia atricapilla* | Palearctic | Passeriformes | Group 3 | Increase | Increase | LC | YES | 16.73 (4.78) |
| *Sylvia borin* | Palearctic | Passeriformes | Group 3 | Decrease | Decrease | LC | YES | 27.61 (12.37) |
| *Sylvietta virens* | Afrotropical | Passeriformes | Group 1 | Decrease | Decrease | LC | YES | 6.29 (3.52) |
| *Syrrhaptes paradoxus* | Palearctic | Pterocliformes | Group 5 | Decrease | Decrease | LC | YES | 12.61 (4.72) |
| *Tachuris rubrigastra* | Neotropical | Passeriformes | | | | LC | NO | 10.19 (3.44) |
| *Tauraco erythrolophus* | Afrotropical | Musophagiformes | Group 6 | Decrease | Decrease | LC | YES | 6.27 (1.45) |
| *Thinocorus orbignyianus* | Neotropical | Charadriiformes | Group 1 | Increase | Increase | LC | YES | 25.51 (14.64) |
| *Thryothorus ludovicianus* | Nearctic | Passeriformes | Group 1 | Increase | Increase | LC | YES | 21.19 (4.67) |
| *Tichodroma muraria* | Palearctic | Passeriformes | | | | LC | NO | 1.27 (0.03) |
| *Tinamus guttatus* | Neotropical | Tinamiformes | Group 3 | Decrease | Decrease | NT | YES | 11.24 (3.62) |
| *Todus mexicanus* | Panamanian | Coraciiformes | Group 6 | Decrease | Decrease | LC | YES | 9.83 (2.63) |
| *Toxostoma redivivum* | Nearctic | Passeriformes | | | | LC | NO | 3.13 (0.93) |
| *Tricholaema leucomelas* | Afrotropical | Piciformes | Group 3 | Decrease | Decrease | LC | YES | 6.43 (1.1) |
| *Trogon melanurus* | Neotropical | Trogoniformes | Group 7 | Increase | Decrease | LC | YES | 29.15 (4.87) |
| *Turnix velox* | Australian | Charadriiformes | Group 4 | Decrease | Decrease | LC | YES | 21.41 (8.32) |
| *Tyrannus savana* | Neotropical | Passeriformes | Group 1 | Increase | Increase | LC | YES | 57.2 (17.53) |
| *Tyto alba* | Nearctic | Strigiformes | Group 7 | Decrease | Increase | LC | YES | 5.11 (0.58) |
| *Upupa epops* | Sino-Japanese | Bucerotiformes | Group 7 | Increase | Decrease | LC | YES | 17.54 (2.04) |
| *Uria aalge* | Palearctic | Charadriiformes | Group 1 | Decrease | Increase | LC | YES | 3.71 (0.46) |
| *Uria lomvia* | Nearctic | Charadriiformes | Group 5 | Increase | Unrelated | LC | YES | 10.16 (3.82) |
| *Urocolius indicus* | Afrotropical | Coliiformes | Group 1 | Increase | Increase | LC | YES | 11.01 (6.6) |

| Species | Realm | Order | Cluster | Warming | Cooling | IUCN | Analyzed | Mean (SD) $N_e$ x $10^4$ |
|---|---|---|---|---|---|---|---|---|
| *Urocynchramus pylzowi* | Sino-Japanese | Passeriformes | Group 4 | Increase | Increase | LC | YES | 11.57 (3.89) |
| *Vidua chalybeata* | Afrotropical | Passeriformes | | | | LC | NO | 4.41 (1.32) |
| *Vidua macroura* | Afrotropical | Passeriformes | Group 2 | Increase | Decrease | LC | YES | 11.43 (3.76) |
| *Vireo altiloquus* | Nearctic | Passeriformes | Group 2 | Decrease | Decrease | LC | YES | 13.33 (3.34) |
| *Xiphorhynchus elegans* | Neotropical | Passeriformes | Group 7 | Decrease | Decrease | LC | YES | 9.6 (0.85) |
| *Zapornia atra* | Oceanian | Gruiformes | | | | VU | NO | 4.93 (7.53) |
| *Zonotrichia albicollis* | Nearctic | Passeriformes | Group 5 | Decrease | Increase | LC | YES | 8.55 (2.68) |
| *Zosterops hypoxanthus* | Oceanian | Passeriformes | Group 1 | Decrease | Decrease | LC | YES | 18.38 (7.54) |

**Table S2**. Robustness values represented by the average of the consistency ratios at five different values of clustering groups (k).

| **Number of clusters** | **3** | **4** | **5** | **6** | **7** |
|---|---|---|---|---|---|
| Robustness (%) | 0.7189 | 0.7803 | 0.7874 | 0.7999 | 0.8349 |

**Table S3.** Chi-square test to determine whether the clustering pattern (k=7) could be explained in part by when species within each cluster group reached their maximum effective population size. The first value in the brackets represent the observed number of species reaching the maximum $N_e$ values in that period, while the second value represents the number of species reaching the maximum $N_e$ values outside that period. Underlined observed values represent significant differences from the background expectations (*: *p-value* < 0.05 **: *p-value* < 0.01. ***: *p-value* < 0.001).

| | Group1 | Group2 | Group3 | Group4 | Group5 | Group6 | Group7 | Background |
|---|---|---|---|---|---|---|---|---|
| **Upper Pleistocene (30–129kya)** | (30, 4)$^{***}$ | (14, 6) | (32, 10)$^{**}$ | (40, 0)$^{***}$ | (0, 35)$^{***}$ | (0, 43)$^{***}$ | (22, 27) | (138, 125) |
| **Middle Pleistocene (129–774kya)** | (5, 29)$^{***}$ | (16, 4) | (33, 9) | (5, 35)$^{***}$ | (27, 8) | (43, 0)$^{***}$ | (45, 4)$^{***}$ | (174, 89) |
| **Lower Pleistocene (774–1,000kya)** | (3, 31) | (0, 20) | (0, 42)$^{*}$ | (0, 40)$^{*}$ | (16, 19)$^{***}$ | (7, 36) | (10, 39) | (36, 227) |

**Table S4**. Summary of three types of demographic responses (“Increase”, “Decrease”, “Unrelated”; quantified by significant positive/negative correlations between $N_e$ and Global Average Surface Temperature) at different confidence levels (confidence level ≥0; confidence level ≥95%; confidence level ≥90%; confidence level ≥85%; confidence level ≥80%) during periods of warming (~147–123kya) and cooling (~122–65kya; see fig. S8).

| | **Confidence level** | **Number of species showing increase response** | **Number of species showing decrease response** | **Number of species showing unrelated response** |
|---|---|---|---|---|
| *Climate Warming* | 0 | 108 | 136 | 19 |
| | 95% | 80 | 109 | 1 |
| | 90% | 85 | 113 | 1 |
| | 85% | 86 | 115 | 2 |
| | 80% | 88 | 117 | 3 |
| *Climate Cooling* | 0 | 91 | 142 | 30 |
| | 95% | 71 | 124 | 4 |
| | 90% | 74 | 129 | 4 |
| | 85% | 76 | 132 | 7 |
| | 80% | 81 | 133 | 9 |

**Table S5.** Overall demographic responses to focal periods of past warming (147–123kya) and cooling (122–65kya; see fig. S8) at different confidence levels. For both the *Climate Warming* and *Climate Cooling* responses, "Increase" and "Decrease" indicate the direction of significant correlations (Pearson's correlation coefficient) between $N_e$ and Global Average Surface Temperature (GAST; data from[51]), where detected. The number of species exhibiting the combined responses during *Climate Warming* and *Climate Cooling* are given, as well as the categorization label used throughout our study.

| Climate Warming | Climate Cooling | Number of species (Confidence level > 0) | Number of species (Confidence level ≥ 95%) | Number of species (Confidence level ≥ 90%) | Number of species (Confidence level ≥ 85%) | Number of species (Confidence level ≥ 80%) | Categorization label |
|---|---|---|---|---|---|---|---|
| Increase | Decrease | 33 | 10 | 14 | 16 | 18 | Warming Positive |
| Decrease | Increase | 29 | 10 | 12 | 13 | 16 | Warming Negative |
| Increase | Increase | 55 | 40 | 43 | 44 | 47 | Consistent $N_e$ Increase |
| Decrease | Decrease | 98 | 80 | 82 | 84 | 84 | Consistent $N_e$ Decrease |
| Demographic change not related to changing GAST during *Climate Warming* or *Climate Cooling* | | 48 | 4 | 4 | 8 | 10 | $N_e$ independent of climate change |

**Table S6**. Details of best performing Phylogenetic Path Analysis (PPA) models (ΔCICc ≤ 2 from top-ranked model) for four comparison groups of combined demographic responses to *Climate Warming* and *Climate Cooling* (see table S5) without limiting the confidence level. For each comparison, the top-ranked model is given in bold. "Model Group" refers to core model categories depicted in fig. S10), "p" represents the p-value (where $p < 0.05$ indicates that the available evidence rejects the model), "CICc" is the size-corrected C-statistic Information criterion, "ΔCICc" is the difference in CICc between the focal model and the top-ranked model, "l" is the relative likelihood, and "w" represents the CICc weights.

| Model Group | p | CICc | ΔCICc | l | w |
|---|---|---|---|---|---|
| "Warming Positive" responses (n = 33) versus all remaining species (n = 230) | | | | | |
| **D** | **0.223** | **63.199** | **0.000** | **1.000** | **0.158** |
| D | 0.186 | 64.224 | 1.025 | 0.599 | 0.094 |
| D | 0.156 | 65.095 | 1.896 | 0.388 | 0.061 |
| "Warming Negative" responses (n = 29) versus all remaining species (n = 234) | | | | | |
| **D** | **0.208** | **63.575** | **0.000** | **1.000** | **0.076** |
| D | 0.192 | 64.012 | 0.437 | 0.804 | 0.061 |
| D | 0.190 | 64.116 | 0.541 | 0.763 | 0.058 |
| D | 0.182 | 64.320 | 0.745 | 0.689 | 0.052 |
| D | 0.173 | 64.580 | 1.005 | 0.605 | 0.046 |
| K | 0.169 | 64.691 | 1.117 | 0.572 | 0.043 |
| D | 0.158 | 65.050 | 1.475 | 0.478 | 0.036 |
| Species sensitive to *Climate Warming* and *Climate Cooling* (n = 33 + 29) versus species with consistent $N_e$ increase or decrease (n = 98 + 55) | | | | | |
| **N** | **0.768** | **52.891** | **0.000** | **1.000** | **0.062** |
| L | 0.712 | 53.991 | 1.100 | 0.577 | 0.036 |
| F | 0.723 | 54.272 | 1.381 | 0.501 | 0.031 |
| L | 0.696 | 54.287 | 1.395 | 0.498 | 0.031 |
| M | 0.717 | 54.388 | 1.497 | 0.473 | 0.030 |
| M | 0.686 | 54.464 | 1.573 | 0.456 | 0.028 |
| M | 0.680 | 54.578 | 1.687 | 0.430 | 0.027 |
| I | 0.696 | 54.756 | 1.865 | 0.394 | 0.025 |

| Model Group | p | CICc | ΔCICc | l | w |
|---|---|---|---|---|---|
| Species which exhibited consistent $N_e$ decrease (n = 98) versus all remaining species (n = 165) | | | | | |
| **L** | **0.222** | **63.222** | **0.000** | **1.000** | **0.049** |
| I | 0.213 | 63.399 | 0.177 | 0.915 | 0.045 |
| N | 0.211 | 63.410 | 0.188 | 0.910 | 0.044 |
| L | 0.202 | 63.689 | 0.467 | 0.792 | 0.039 |
| I | 0.191 | 64.093 | 0.871 | 0.647 | 0.032 |
| D | 0.185 | 64.212 | 0.989 | 0.610 | 0.030 |
| F | 0.181 | 64.345 | 1.122 | 0.571 | 0.028 |
| K | 0.180 | 64.356 | 1.133 | 0.567 | 0.028 |
| K | 0.176 | 64.479 | 1.256 | 0.534 | 0.026 |
| F | 0.174 | 64.550 | 1.327 | 0.515 | 0.025 |
| F | 0.170 | 64.678 | 1.455 | 0.483 | 0.024 |
| F | 0.170 | 64.685 | 1.462 | 0.481 | 0.023 |
| I | 0.168 | 64.722 | 1.500 | 0.472 | 0.023 |
| D | 0.168 | 64.732 | 1.510 | 0.470 | 0.023 |
| D | 0.168 | 64.743 | 1.520 | 0.468 | 0.023 |
| I | 0.163 | 64.904 | 1.682 | 0.431 | 0.021 |
| I | 0.160 | 64.985 | 1.763 | 0.414 | 0.020 |
| L | 0.161 | 64.997 | 1.775 | 0.412 | 0.020 |
| L | 0.160 | 64.998 | 1.776 | 0.411 | 0.020 |
| L | 0.157 | 65.127 | 1.904 | 0.386 | 0.019 |
| I | 0.155 | 65.168 | 1.946 | 0.378 | 0.018 |

**Table S7.** Details of best performing Phylogenetic Path Analysis (PPA) models (ΔCICc ≤ 2 from top-ranked model) for four comparison groups of combined demographic responses to *Climate Warming* and *Climate Cooling* (see table S5) when setting the confidence level as 95%. For each comparison, the top-ranked model is given in bold. “Model Group” refers to core model categories depicted in fig. S10), “p” represents the p-value (where $p < 0.05$ indicates that the available evidence rejects the model), “CICc” is the size-corrected C-statistic Information criterion, “ΔCICc” is the difference in CICc between the focal model and the top-ranked model, “l” is the relative likelihood, and “w” represents the CICc weights.

| Model Group | p | CICc | ΔCICc | l | w |
|---|---|---|---|---|---|
| “Warming Positive” responses (n = 10) versus all remaining species (n = 134) | | | | | |
| **F** | **0.568** | **58.481** | **0.000** | **1.000** | **0.073** |
| L | 0.530 | 58.549 | 0.069 | 0.966 | 0.071 |
| D | 0.600 | 58.593 | 0.113 | 0.945 | 0.069 |
| D | 0.619 | 58.999 | 0.519 | 0.772 | 0.057 |
| F | 0.567 | 59.108 | 0.627 | 0.731 | 0.054 |
| F | 0.559 | 59.246 | 0.766 | 0.682 | 0.050 |
| L | 0.467 | 59.681 | 1.200 | 0.549 | 0.040 |
| F | 0.518 | 59.909 | 1.428 | 0.490 | 0.036 |
| L | 0.474 | 60.069 | 1.588 | 0.452 | 0.033 |
| D | 0.584 | 60.251 | 1.771 | 0.413 | 0.030 |
| “Warming Negative” responses (n = 10) versus all remaining species (n = 134) | | | | | |
| **L** | **0.784** | **55.470** | **0.000** | **1.000** | **0.164** |
| F | 0.731 | 57.240 | 1.770 | 0.413 | 0.068 |
| Species sensitive to *Climate Warming* and *Climate Cooling* (n = 10 + 10) versus species with consistent $N_e$ increase or decrease (n = 80 + 40) | | | | | |
| **F** | **0.651** | **58.690** | **0.000** | **1.000** | **0.134** |
| F | 0.582 | 59.030 | 0.340 | 0.843 | 0.113 |
| D | 0.623 | 59.890 | 1.200 | 0.549 | 0.074 |
| D | 0.541 | 60.352 | 1.662 | 0.436 | 0.058 |
| F | 0.497 | 60.404 | 1.714 | 0.424 | 0.057 |

| **Model Group** | **p** | **CICc** | **ΔCICc** | **l** | **w** |
|---|---|---|---|---|---|
| Species which exhibited consistent $N_e$ decrease (n = 80) versus all remaining species (n = 64) | | | | | |
| **I** | **0.781** | **56.387** | **0.000** | **1.000** | **0.099** |
| I | 0.709 | 56.809 | 0.422 | 0.810 | 0.080 |
| L | 0.694 | 57.055 | 0.668 | 0.716 | 0.071 |
| I | 0.589 | 58.123 | 1.736 | 0.420 | 0.042 |

**Table S8.** Details of best performing Phylogenetic Path Analysis (PPA) models (ΔCICc ≤ 2 from top-ranked model) for four comparison groups of combined demographic responses to *Climate Warming* and *Climate Cooling* (see table S5) when setting the confidence level as 90%. For each comparison, the top-ranked model is given in bold. "Model Group" refers to core model categories depicted in fig. S10), "p" represents the p-value (where $p < 0.05$ indicates that the available evidence rejects the model), "CICc" is the size-corrected C-statistic Information criterion, "ΔCICc" is the difference in CICc between the focal model and the top-ranked model, "l" is the relative likelihood, and "w" represents the CICc weights.

| Model Group | p | CICc | ΔCICc | l | w |
|---|---|---|---|---|---|
| "Warming Positive" responses (n = 14) versus all remaining species (n = 141) | | | | | |
| **D** | **0.624** | **58.480** | **0.000** | **1.000** | **0.073** |
| M | 0.517 | 58.486 | 0.006 | 0.997 | 0.072 |
| F | 0.566 | 58.740 | 0.260 | 0.878 | 0.064 |
| D | 0.640 | 58.949 | 0.469 | 0.791 | 0.057 |
| N | 0.457 | 59.565 | 1.085 | 0.581 | 0.042 |
| D | 0.591 | 59.651 | 1.171 | 0.557 | 0.040 |
| L | 0.450 | 59.699 | 1.219 | 0.544 | 0.039 |
| D | 0.501 | 59.791 | 1.311 | 0.519 | 0.038 |
| D | 0.616 | 60.039 | 1.558 | 0.459 | 0.033 |
| D | 0.519 | 60.075 | 1.595 | 0.450 | 0.033 |
| F | 0.449 | 60.160 | 1.680 | 0.432 | 0.031 |
| F | 0.474 | 60.225 | 1.745 | 0.418 | 0.030 |
| F | 0.470 | 60.295 | 1.815 | 0.404 | 0.029 |
| F | 0.501 | 60.347 | 1.867 | 0.393 | 0.029 |
| F | 0.493 | 60.480 | 2.000 | 0.368 | 0.027 |
| "Warming Negative" responses (n = 12) versus all remaining species (n = 143) | | | | | |
| **F** | **0.820** | **53.578** | **0.000** | **1.000** | **0.102** |
| L | 0.795 | 54.111 | 0.533 | 0.766 | 0.078 |
| L | 0.833 | 54.114 | 0.536 | 0.765 | 0.078 |
| M | 0.728 | 54.725 | 1.147 | 0.564 | 0.058 |
| L | 0.706 | 55.140 | 1.562 | 0.458 | 0.047 |
| F | 0.779 | 55.188 | 1.610 | 0.447 | 0.046 |

| Model Group | p | CICc | ΔCICc | l | w |
|---|---|---|---|---|---|
| F | 0.803 | 55.554 | 1.976 | 0.372 | 0.038 |
| Species sensitive to *Climate Warming* and *Climate Cooling* (n = 14 + 12) versus species with consistent $N_e$ increase or decrease (n = 82 + 43) | | | | | |
| **D** | **0.727** | **57.011** | **0.000** | **1.000** | **0.070** |
| D | 0.768 | 57.184 | 0.173 | 0.917 | 0.065 |
| F | 0.668 | 57.231 | 0.219 | 0.896 | 0.063 |
| D | 0.680 | 57.768 | 0.757 | 0.685 | 0.048 |
| K | 0.566 | 58.288 | 1.277 | 0.528 | 0.037 |
| F | 0.544 | 58.657 | 1.646 | 0.439 | 0.031 |
| D | 0.658 | 58.864 | 1.853 | 0.396 | 0.028 |
| D | 0.565 | 58.889 | 1.878 | 0.391 | 0.028 |
| F | 0.606 | 58.906 | 1.895 | 0.388 | 0.027 |
| F | 0.650 | 58.977 | 1.966 | 0.374 | 0.026 |
| Species which exhibited consistent $N_e$ decrease (n = 82) versus all remaining species (n = 73) | | | | | |
| **I** | **0.823** | **55.172** | **0.000** | **1.000** | **0.142** |
| L | 0.681 | 56.882 | 1.710 | 0.425 | 0.060 |

**Table S9.** Details of best performing Phylogenetic Path Analysis (PPA) models (ΔCICc ≤ 2 from top-ranked model) for four comparison groups of combined demographic responses to *Climate Warming* and *Climate Cooling* (see table S5) when setting the confidence level as 85%. For each comparison, the top-ranked model is given in bold. "Model Group" refers to core model categories depicted in fig. S10), "p" represents the p-value (where $p < 0.05$ indicates that the available evidence rejects the model), "CICc" is the size-corrected C-statistic Information criterion, "ΔCICc" is the difference in CICc between the focal model and the top-ranked model, "l" is the relative likelihood, and "w" represents the CICc weights.

| Model Group | p | CICc | ΔCICc | l | w |
|---|---|---|---|---|---|
| "Warming Positive" responses (n = 16) versus all remaining species (n = 149) | | | | | |
| **D** | **0.589** | **58.065** | **0.000** | **1.000** | **0.069** |
| D | 0.612 | 58.310 | 0.245 | 0.885 | 0.061 |
| I | 0.538 | 58.374 | 0.309 | 0.857 | 0.059 |
| D | 0.601 | 58.478 | 0.413 | 0.813 | 0.056 |
| D | 0.636 | 58.621 | 0.556 | 0.757 | 0.052 |
| F | 0.513 | 58.789 | 0.724 | 0.696 | 0.048 |
| F | 0.505 | 59.420 | 1.355 | 0.508 | 0.035 |
| D | 0.569 | 59.576 | 1.511 | 0.470 | 0.032 |
| D | 0.616 | 59.604 | 1.539 | 0.463 | 0.032 |
| D | 0.564 | 59.649 | 1.584 | 0.453 | 0.031 |
| D | 0.483 | 59.778 | 1.713 | 0.425 | 0.029 |
| L | 0.432 | 59.793 | 1.728 | 0.421 | 0.029 |
| K | 0.444 | 59.993 | 1.928 | 0.381 | 0.026 |
| D | 0.497 | 60.064 | 1.999 | 0.368 | 0.025 |
| "Warming Negative" responses (n = 13) versus all remaining species (n = 152) | | | | | |
| **L** | **0.852** | **52.597** | **0.000** | **1.000** | **0.098** |
| L | 0.858 | 53.254 | 0.657 | 0.720 | 0.071 |
| F | 0.811 | 53.511 | 0.914 | 0.633 | 0.062 |
| L | 0.766 | 53.746 | 1.148 | 0.563 | 0.055 |
| F | 0.796 | 53.825 | 1.227 | 0.541 | 0.053 |
| L | 0.750 | 54.064 | 1.466 | 0.480 | 0.047 |
| F | 0.798 | 54.516 | 1.919 | 0.383 | 0.038 |

| Model Group | p | CICc | ΔCICc | l | w |
|---|---|---|---|---|---|
| M | 0.726 | 54.526 | 1.929 | 0.381 | 0.037 |
| L | 0.723 | 54.592 | 1.994 | 0.369 | 0.036 |
| Species sensitive to *Climate Warming* and *Climate Cooling* (n = 16 + 13) versus species with consistent $N_e$ increase or decrease (n = 84 + 44) | | | | | |
| **K** | **0.730** | **55.284** | **0.000** | **1.000** | **0.063** |
| D | 0.857 | 55.350 | 0.065 | 0.968 | 0.061 |
| F | 0.754 | 55.579 | 0.295 | 0.863 | 0.055 |
| D | 0.789 | 55.731 | 0.447 | 0.800 | 0.051 |
| D | 0.764 | 56.164 | 0.880 | 0.644 | 0.041 |
| F | 0.763 | 56.182 | 0.898 | 0.638 | 0.040 |
| M | 0.635 | 56.372 | 1.088 | 0.581 | 0.037 |
| I | 0.666 | 56.425 | 1.141 | 0.565 | 0.036 |
| D | 0.701 | 56.481 | 1.196 | 0.550 | 0.035 |
| F | 0.694 | 56.606 | 1.321 | 0.517 | 0.033 |
| D | 0.762 | 57.027 | 1.743 | 0.418 | 0.026 |
| K | 0.661 | 57.147 | 1.863 | 0.394 | 0.025 |
| Species which exhibited consistent $N_e$ decrease (n = 84) versus all remaining species (n = 81) | | | | | |
| **K** | **0.687** | **55.850** | **0.000** | **1.000** | **0.049** |
| I | 0.714 | 56.023 | 0.173 | 0.917 | 0.045 |
| D | 0.756 | 56.044 | 0.194 | 0.908 | 0.044 |
| N | 0.637 | 56.148 | 0.298 | 0.862 | 0.042 |
| I | 0.664 | 56.249 | 0.399 | 0.819 | 0.040 |
| I | 0.725 | 56.543 | 0.693 | 0.707 | 0.034 |
| K | 0.639 | 56.678 | 0.828 | 0.661 | 0.032 |
| F | 0.667 | 56.809 | 0.960 | 0.619 | 0.030 |
| D | 0.662 | 56.894 | 1.045 | 0.593 | 0.029 |
| D | 0.649 | 57.102 | 1.253 | 0.535 | 0.026 |
| F | 0.612 | 57.122 | 1.272 | 0.529 | 0.026 |
| D | 0.728 | 57.254 | 1.404 | 0.495 | 0.024 |
| L | 0.638 | 57.278 | 1.428 | 0.490 | 0.024 |
| L | 0.553 | 57.617 | 1.767 | 0.413 | 0.020 |
| D | 0.615 | 57.649 | 1.799 | 0.407 | 0.020 |
| F | 0.580 | 57.664 | 1.814 | 0.404 | 0.020 |

**Table S10.** Details of best performing Phylogenetic Path Analysis (PPA) models ($\Delta$CICc $\leq 2$ from top-ranked model) for four comparison groups of combined demographic responses to *Climate Warming* and *Climate Cooling* (see table S5) when setting the confidence level as 80%. For each comparison, the top-ranked model is given in bold. "Model Group" refers to core model categories depicted in fig. S10), "p" represents the p-value (where $p < 0.05$ indicates that the available evidence rejects the model), "CICc" is the size-corrected C-statistic Information criterion, "$\Delta$CICc" is the difference in CICc between the focal model and the top-ranked model, "l" is the relative likelihood, and "w" represents the CICc weights.

| **Model Group** | **p** | **CICc** | **ΔCICc** | **l** | **w** |
|---|---|---|---|---|---|
| "Warming Positive" responses (n = 18) versus all remaining species (n = 157) | | | | | |
| **D** | **0.620** | **57.308** | **0.000** | **1.000** | **0.072** |
| F | 0.614 | 57.397 | 0.090 | 0.956 | 0.068 |
| D | 0.640 | 57.578 | 0.271 | 0.873 | 0.063 |
| D | 0.639 | 57.603 | 0.296 | 0.863 | 0.062 |
| D | 0.679 | 57.650 | 0.343 | 0.842 | 0.060 |
| F | 0.589 | 58.360 | 1.053 | 0.591 | 0.042 |
| F | 0.524 | 58.369 | 1.062 | 0.588 | 0.042 |
| D | 0.675 | 58.424 | 1.116 | 0.572 | 0.041 |
| D | 0.614 | 58.600 | 1.292 | 0.524 | 0.038 |
| D | 0.591 | 58.919 | 1.612 | 0.447 | 0.032 |
| F | 0.519 | 58.926 | 1.618 | 0.445 | 0.032 |
| I | 0.480 | 59.117 | 1.810 | 0.405 | 0.029 |
| F | 0.504 | 59.177 | 1.870 | 0.393 | 0.028 |
| F | 0.533 | 59.209 | 1.901 | 0.387 | 0.028 |
| D | 0.529 | 59.271 | 1.964 | 0.375 | 0.027 |
| F | 0.566 | 59.279 | 1.971 | 0.373 | 0.027 |
| "Warming Negative" responses (n = 16) versus all remaining species (n = 159) | | | | | |
| **I** | **0.854** | **53.081** | **0.000** | **1.000** | **0.088** |
| I | 0.808 | 53.348 | 0.267 | 0.875 | 0.077 |
| L | 0.787 | 53.765 | 0.684 | 0.710 | 0.063 |
| I | 0.799 | 54.238 | 1.157 | 0.561 | 0.050 |
| L | 0.724 | 54.360 | 1.279 | 0.528 | 0.047 |

| Model Group | p | CICc | ΔCICc | l | w |
|---|---|---|---|---|---|
| F | 0.744 | 54.584 | 1.503 | 0.472 | 0.042 |
| F | 0.726 | 54.917 | 1.836 | 0.399 | 0.035 |
| F | 0.759 | 54.970 | 1.889 | 0.389 | 0.034 |
| Species sensitive to *Climate Warming* and *Climate Cooling* (n = 18 + 16) versus species with consistent $N_e$ increase or decrease (n = 84 + 47) | | | | | |
| **F** | **0.895** | **52.331** | **0.000** | **1.000** | **0.126** |
| F | 0.902 | 53.076 | 0.745 | 0.689 | 0.087 |
| D | 0.883 | 53.547 | 1.216 | 0.544 | 0.068 |
| F | 0.826 | 53.967 | 1.636 | 0.441 | 0.055 |
| F | 0.862 | 54.029 | 1.698 | 0.428 | 0.054 |
| D | 0.893 | 54.283 | 1.951 | 0.377 | 0.047 |
| Species which exhibited consistent $N_e$ decrease (n = 84) versus all remaining species (n = 91) | | | | | |
| **F** | **0.816** | **53.889** | **0.000** | **1.000** | **0.047** |
| K | 0.778 | 53.938 | 0.049 | 0.976 | 0.046 |
| M | 0.742 | 54.013 | 0.125 | 0.940 | 0.044 |
| F | 0.763 | 54.229 | 0.340 | 0.844 | 0.040 |
| D | 0.835 | 54.286 | 0.398 | 0.820 | 0.039 |
| L | 0.706 | 54.699 | 0.811 | 0.667 | 0.031 |
| F | 0.728 | 54.886 | 0.997 | 0.607 | 0.029 |
| I | 0.715 | 55.118 | 1.229 | 0.541 | 0.025 |
| F | 0.746 | 55.212 | 1.323 | 0.516 | 0.024 |
| D | 0.741 | 55.302 | 1.414 | 0.493 | 0.023 |
| F | 0.779 | 55.345 | 1.456 | 0.483 | 0.023 |
| K | 0.698 | 55.415 | 1.526 | 0.466 | 0.022 |
| L | 0.688 | 55.598 | 1.709 | 0.426 | 0.020 |
| M | 0.657 | 55.600 | 1.712 | 0.425 | 0.020 |
| L | 0.651 | 55.690 | 1.801 | 0.406 | 0.019 |
| D | 0.713 | 55.778 | 1.889 | 0.389 | 0.018 |

**Table S11.** List of morphological and life-history traits initially selected for analysis of trait-based influences on long-term demographic responses to climate change, based on links between each trait and population responses to climate change over ecological time scales from contemporary data.

| Category | Trait | | Predicted response to climate change | Reference |
|---|---|---|---|---|
| **Survival/Growth** | | | | |
| | Body mass | | smaller species less sensitive to climate change (particularly warming) | 25,90,91 |
| | Body size | *measured as:* | small-bodied species better able to exploit shelter/micro-climate under changing temperature | 25,92 |
| | | tarsus length | | |
| | | wing length | | |
| | | bill length/width/depth | | |
| | Relative brain size | | Problem solving and flexible resource use allow increased ability to cope with environmental change | 93,94 |
| | Generation time | | Influence of demographic and environmental stochasticity on population dynamics decreases with longer generation time | 95–98 |
| | Maximum longevity | | longer-lived species respond slowly to selective pressure, higher extinction probability | 96,98 |
| | Mortality Rate | | longer-lived species respond slowly to selective pressure, higher extinction probability | 96,98 |

| Category | Trait | | Predicted response to climate change | Reference |
|---|---|---|---|---|
| **Fecundity** | | | | |
| | Clutch size | | Higher reproductive rate linked to increased colonization opportunity under climate change | 24,99 |
| | Egg mass | | Higher energetic investment in eggs favorable under warm conditions | 24,100,101 |
| | Incubation duration | | Longer incubation periods typical of species with slow responses to selective pressure, higher extinction probability | 96–98,102 |
| **Movement/Dispersal** | | | | |
| | Elevational range | *measured as:* | High-elevation species already near climatic and geographic limit, more likely to decline under climate change | 13,25 |
| | | elevation (min) | | |
| | | elevation (max) | | |
| | Movement ability | *measured as:* | Allows for spatial shifts under adverse climate, particularly given phenological changes on breeding grounds | 15,91,103,104 |
| | | Kipps distance | | |
| | | hand-wing index | | |

**Table S12**. Details of best performing Phylogenetic Path Analysis (PPA) models (ΔCICc ≤ 2 from top-ranked model) for comparison between species which exhibited increasing $N_e$ tendency under *Climate Warming* verses those with decreasing $N_e$ tendency under *Climate Warming* at different confidence levels. The top-ranked model is given in bold. "Model Group" refers to core model categories depicted in fig. S10), "p" represents the p-value (where $p < 0.05$ indicates that the available evidence rejects the model), "CICc" is the size-corrected C-statistic Information criterion, "ΔCICc" is the difference in CICc between the focal model and the top-ranked model, "l" is the relative likelihood, and "w" represents the CICc weights.

| **Model Group** | **p** | **CICc** | **ΔCICc** | **l** | **w** |
|---|---|---|---|---|---|
| Confidence level > 0: species with increasing Ne tendency (n = 108 species) verses those with decreasing Ne tendency (n = 136) under Climate Warming | | | | | |
| N | 0.231 | 63.023 | 0.000 | 1.000 | 0.076 |
| I | 0.225 | 63.267 | 0.244 | 0.885 | 0.068 |
| I | 0.197 | 64.034 | 1.011 | 0.603 | 0.046 |
| L | 0.185 | 64.352 | 1.329 | 0.515 | 0.039 |
| K | 0.184 | 64.405 | 1.382 | 0.501 | 0.038 |
| D | 0.184 | 64.475 | 1.452 | 0.484 | 0.037 |
| I | 0.180 | 64.589 | 1.566 | 0.457 | 0.035 |
| D | 0.169 | 64.914 | 1.891 | 0.389 | 0.030 |
| Confidence level ≥ 95%: species with increasing Ne tendency (n = 80 species) verses those with decreasing Ne tendency (n = 109) under Climate Warming | | | | | |
| N | 0.456 | 58.906 | 0.000 | 1.000 | 0.130 |
| Confidence level ≥ 90%: species with increasing Ne tendency (n = 85 species) verses those with decreasing Ne tendency (n = 113) under Climate Warming | | | | | |
| N | 0.409 | 59.636 | 0.000 | 1.000 | 0.117 |
| I | 0.362 | 60.834 | 1.198 | 0.549 | 0.064 |
| L | 0.321 | 61.431 | 1.794 | 0.408 | 0.048 |
| I | 0.331 | 61.445 | 1.808 | 0.405 | 0.047 |

| **Model Group** | **p** | **CICc** | **ΔCICc** | **l** | **w** |
|---|---|---|---|---|---|
| Confidence level ≥ 85%: species with increasing Ne tendency (n = 86 species) verses those with decreasing Ne tendency (n = 115) under Climate Warming | | | | | |
| I | 0.471 | 58.795 | 0.000 | 1.000 | 0.078 |
| N | 0.414 | 59.514 | 0.720 | 0.698 | 0.055 |
| K | 0.408 | 59.909 | 1.114 | 0.573 | 0.045 |
| I | 0.396 | 60.130 | 1.336 | 0.513 | 0.040 |
| D | 0.387 | 60.608 | 1.814 | 0.404 | 0.032 |
| I | 0.387 | 60.614 | 1.819 | 0.403 | 0.032 |
| I | 0.385 | 60.636 | 1.841 | 0.398 | 0.031 |
| Confidence level ≥ 80%: species with increasing Ne tendency (n = 88 species) verses those with decreasing Ne tendency (n = 117) under Climate Warming | | | | | |
| N | 0.402 | 59.681 | 0.000 | 1.000 | 0.067 |
| I | 0.401 | 59.980 | 0.298 | 0.861 | 0.058 |
| I | 0.393 | 60.425 | 0.744 | 0.689 | 0.046 |
| D | 0.367 | 60.904 | 1.222 | 0.543 | 0.036 |
| I | 0.360 | 61.023 | 1.342 | 0.511 | 0.034 |
| I | 0.346 | 61.037 | 1.355 | 0.508 | 0.034 |
| K | 0.344 | 61.071 | 1.390 | 0.499 | 0.034 |
| K | 0.325 | 61.469 | 1.788 | 0.409 | 0.028 |
| D | 0.330 | 61.615 | 1.934 | 0.380 | 0.026 |

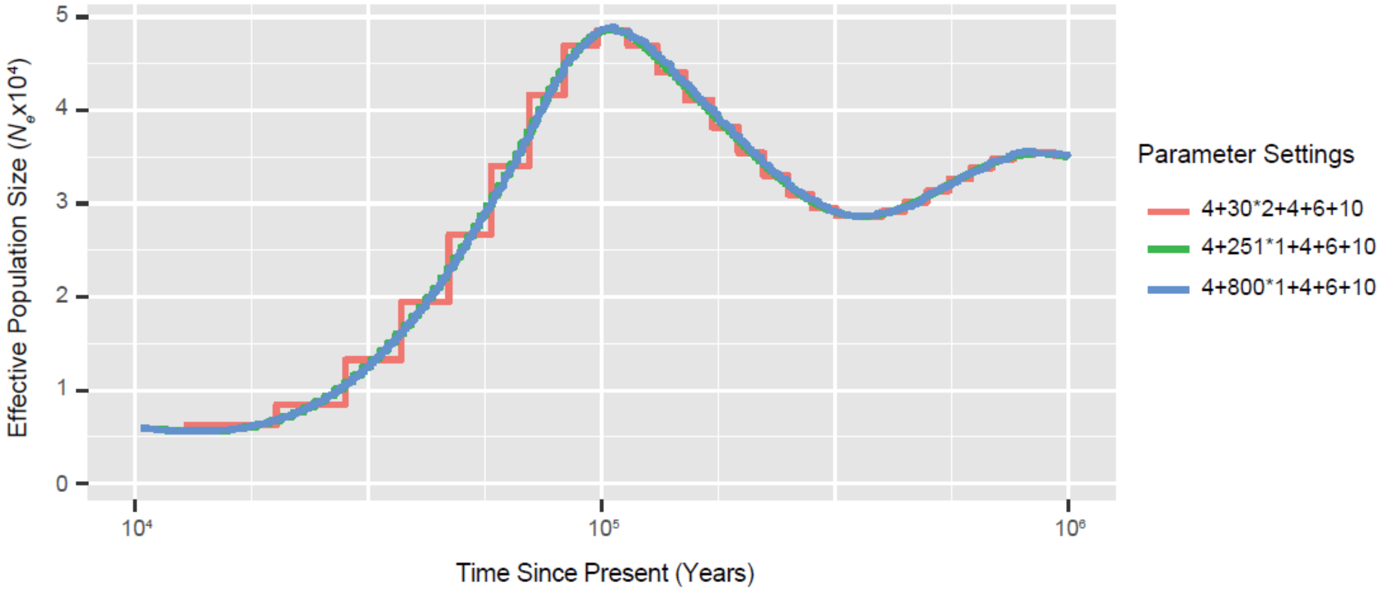

**Fig. S1.** PSMC curves of *Menura novaehollandiae* under three parameter settings: 1) -N30 -t5 -r5 -p 4+30*2+4+6+10; 2) -N30 -t5 -r5 -p 4+251*1+4+6+10; and 3) -N30 -t5 -r5 -p 4+800*1+4+6+10.

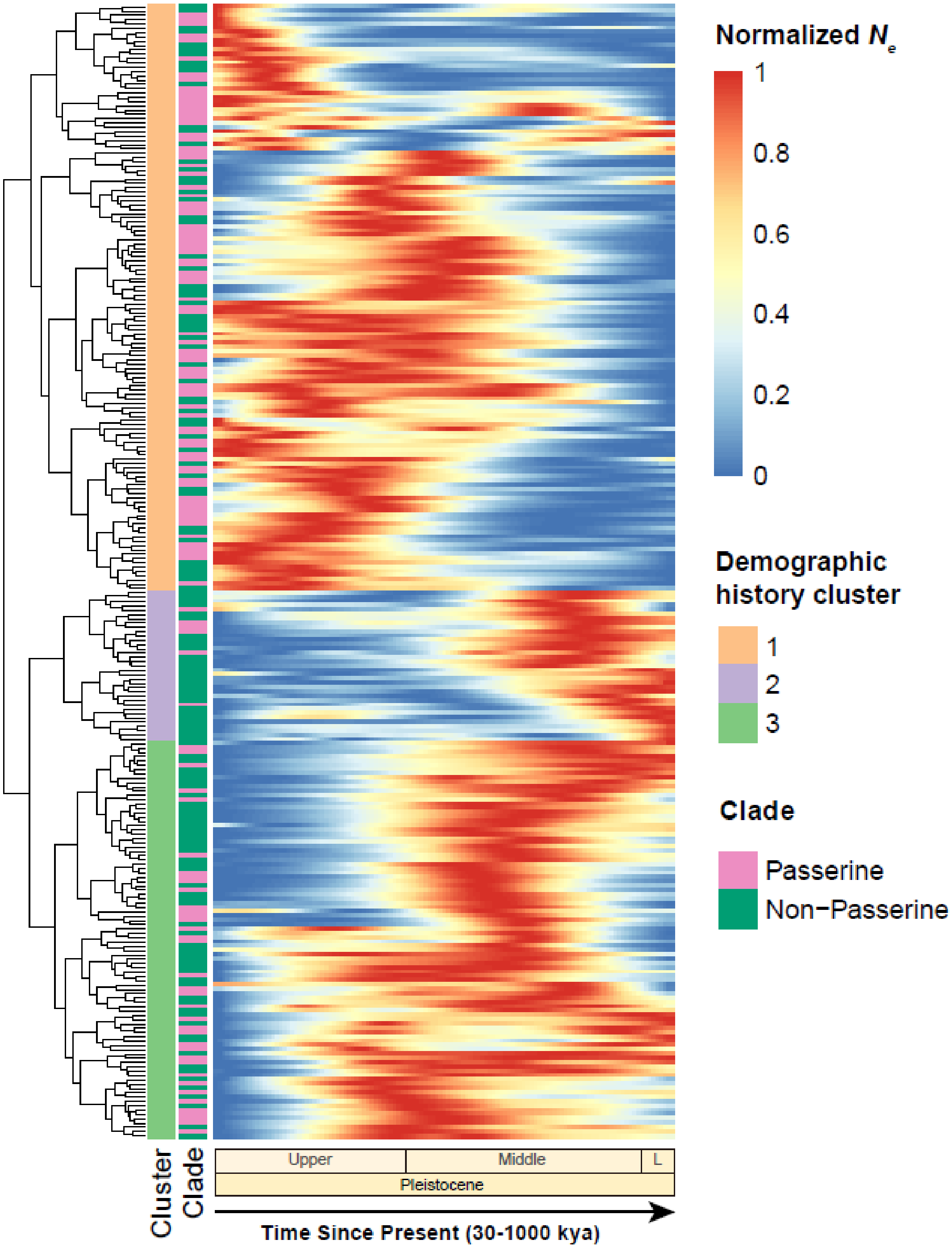

**Fig. S2.** Demographic histories of 263 avian species from 30,000 to 1 million years ago (all x-axes presented on the log10 scale), when splitting the clustering dendrogram into 3 groups based on the overall similarity of long-term $N_e$ patterns during the Upper/Middle/Lower(L) Pleistocene.

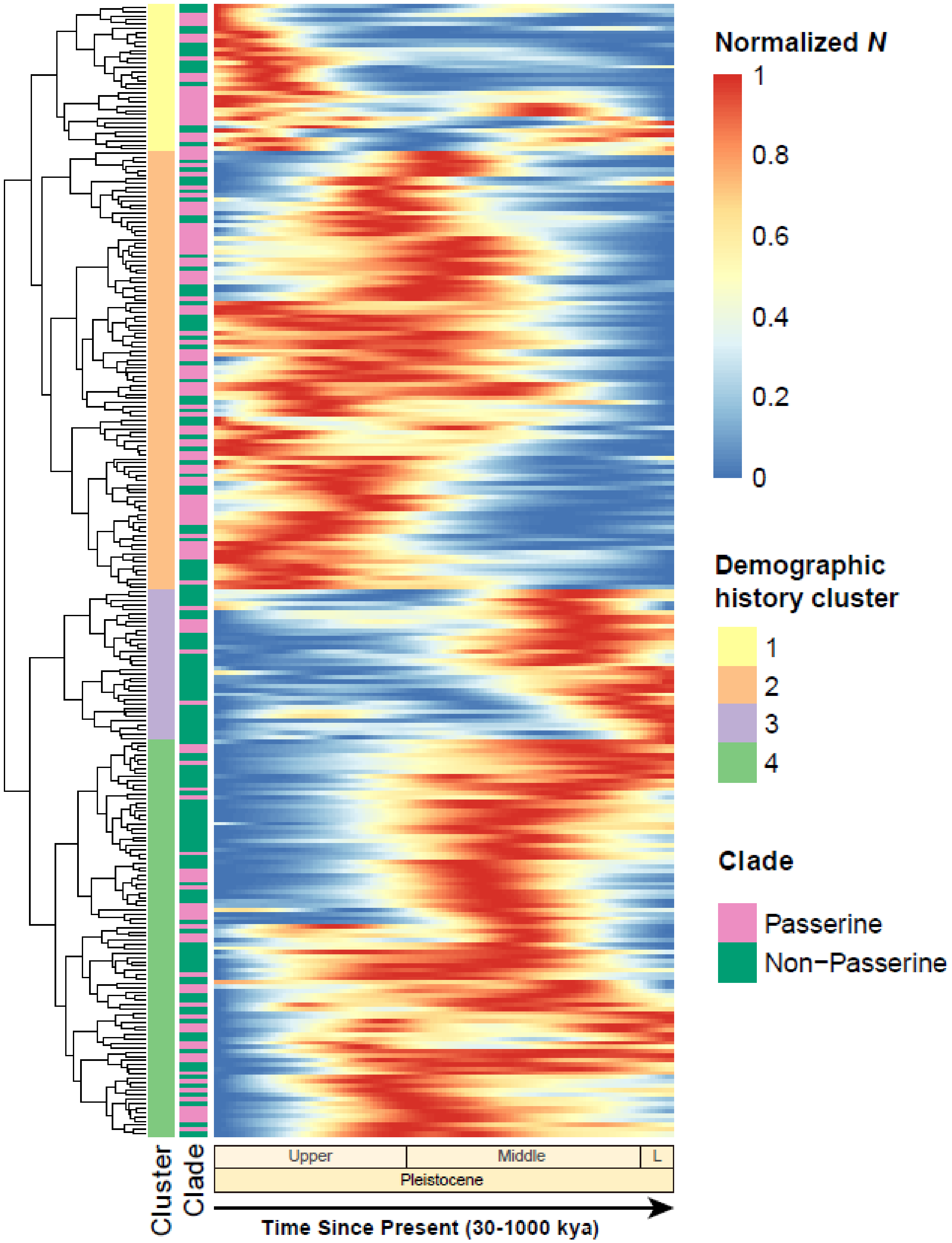

**Fig. S3**. Demographic histories of 263 avian species from 30,000 to 1 million years ago (all x-axes presented on the log10 scale), when splitting the clustering dendrogram into 4 groups based on the overall similarity of long-term $N_e$ patterns during the Upper/Middle/Lower(L) Pleistocene.

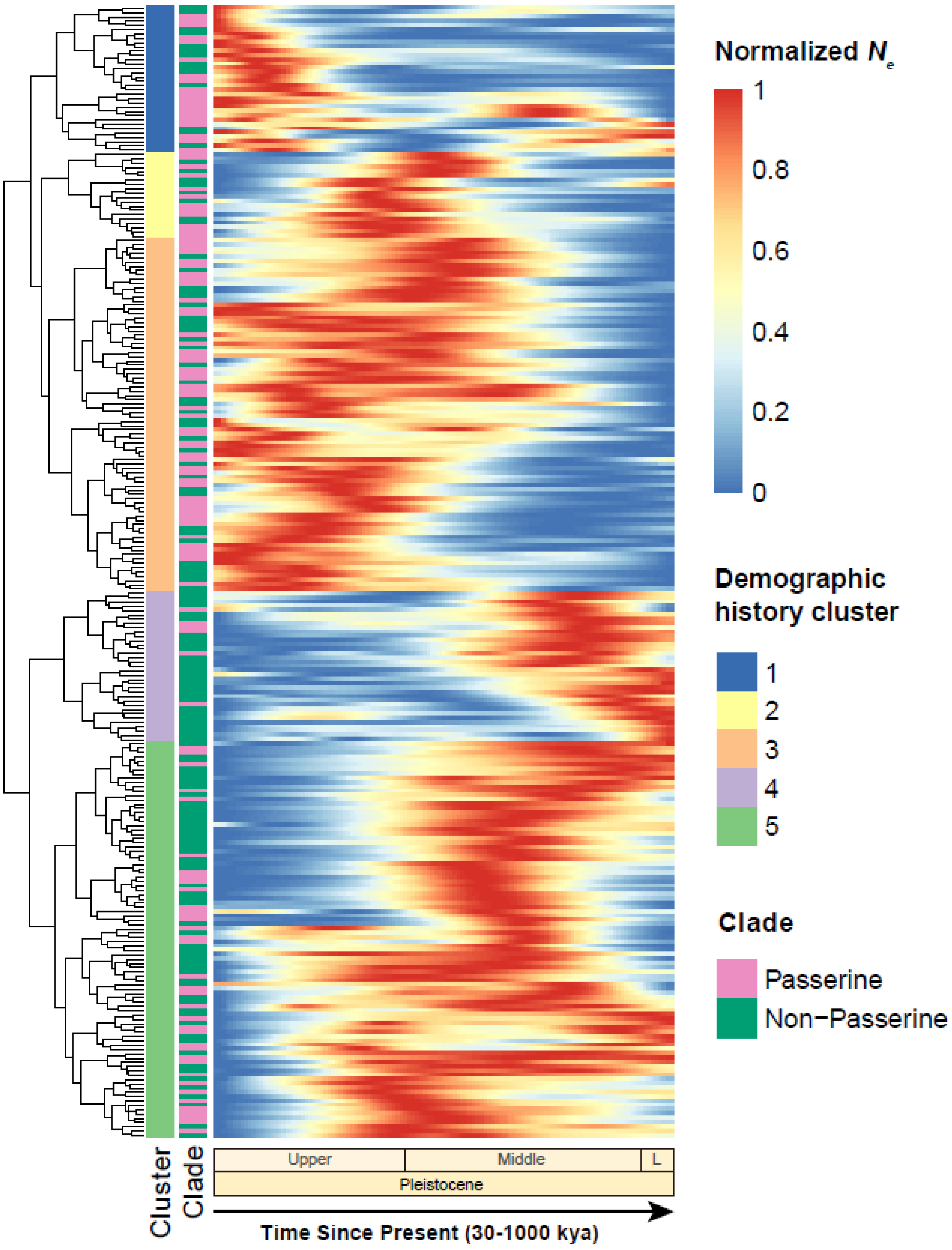

**Fig. S4**. Demographic histories of 263 avian species from 30,000 to 1 million years ago (all x-axes presented on the log10 scale), when splitting the clustering dendrogram into 5 groups based on the overall similarity of long-term $N_e$ patterns during the Upper/Middle/Lower(L) Pleistocene.

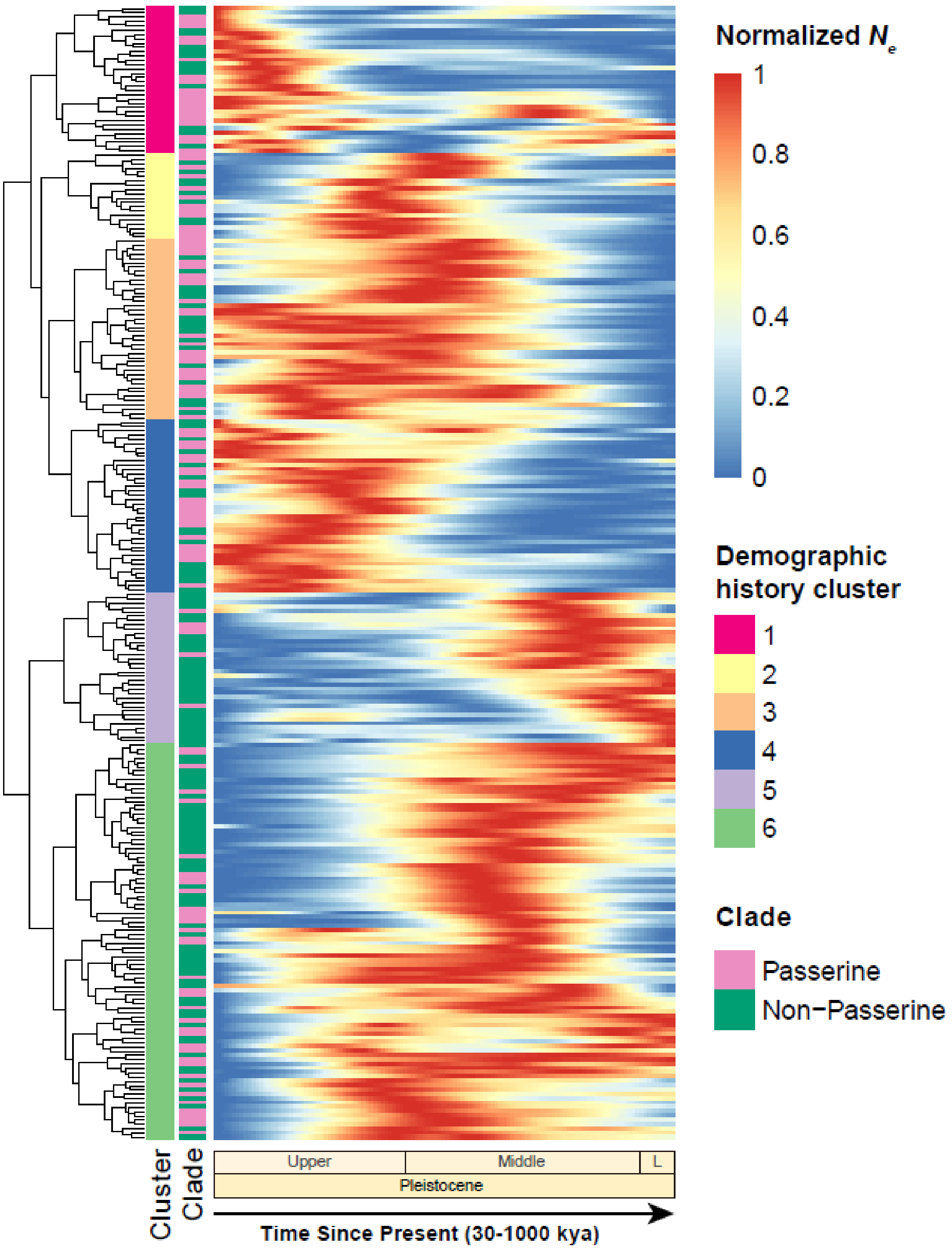

**Fig. S5**. Demographic histories of 263 avian species from 30,000 to 1 million years ago (all x-axes presented on the log10 scale), when splitting the clustering dendrogram into 6 groups based on the overall similarity of long-term $N_e$ patterns during the Upper/Middle/Lower(L) Pleistocene.

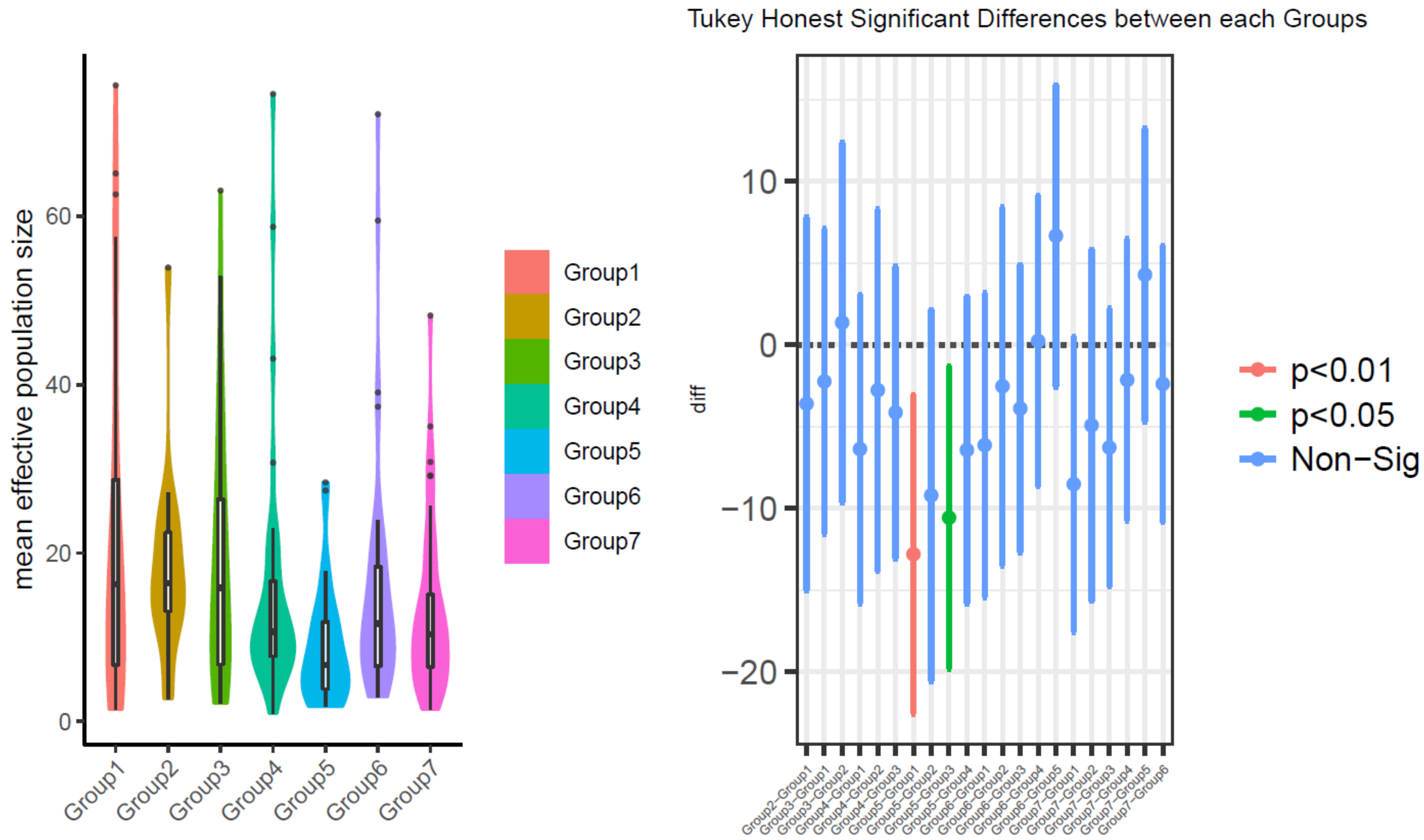

**Fig S6.** Analysis of variance (ANOVA) comparing intra-group and inter-group differences of mean $N_e$ values for the k=7 demographic history clusters (Groups 1–7).

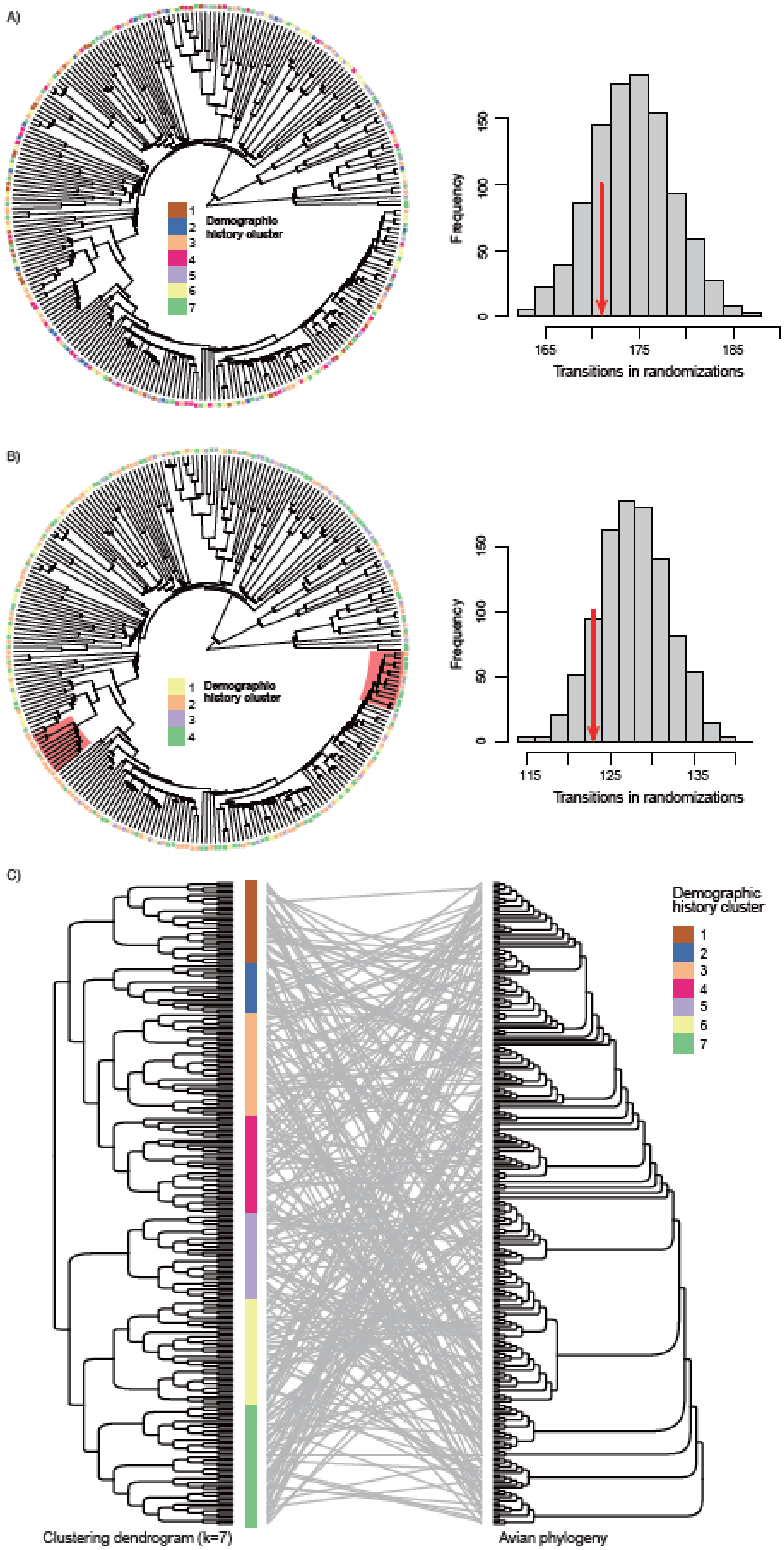
A)
Demographic history cluster
1
2
3
4
5
6
7
Frequency
0
50
100
150
165
175
185
Transitions in randomizations
B)
Demographic history cluster
1
2
3
4
Frequency
0
50
100
150
115
125
135
Transitions in randomizations
C)
Demographic history cluster
1
2
3
4
5
6
7
Clustering dendrogram (k=7)
Avian phylogeny

**Fig. S7**. Phylogenetic signals of the overall demographic clusters. A) The demographic cluster (k=7) mapped, as a categorical variable, within the phylogeny on the left. The null model of evolutionary transitions is shown in the right histogram with an arrow highlighting the number of observed transitions of clustering labels. B) The demographic cluster (k=4) mapped within the phylogeny on the left. Some phylogenetically related lineages showing the similar $N_e$ trajectories are labeled by the red background. The null model of evolutionary transitions is shown in the right histogram with an arrow highlighting the number of observed transitions of clustering labels. C) Comparison of the position of the species on the clustering dendrogram (k=7) and the most up-to date avian phylogeny developed using B10k resources (Stiller et al., in prep).

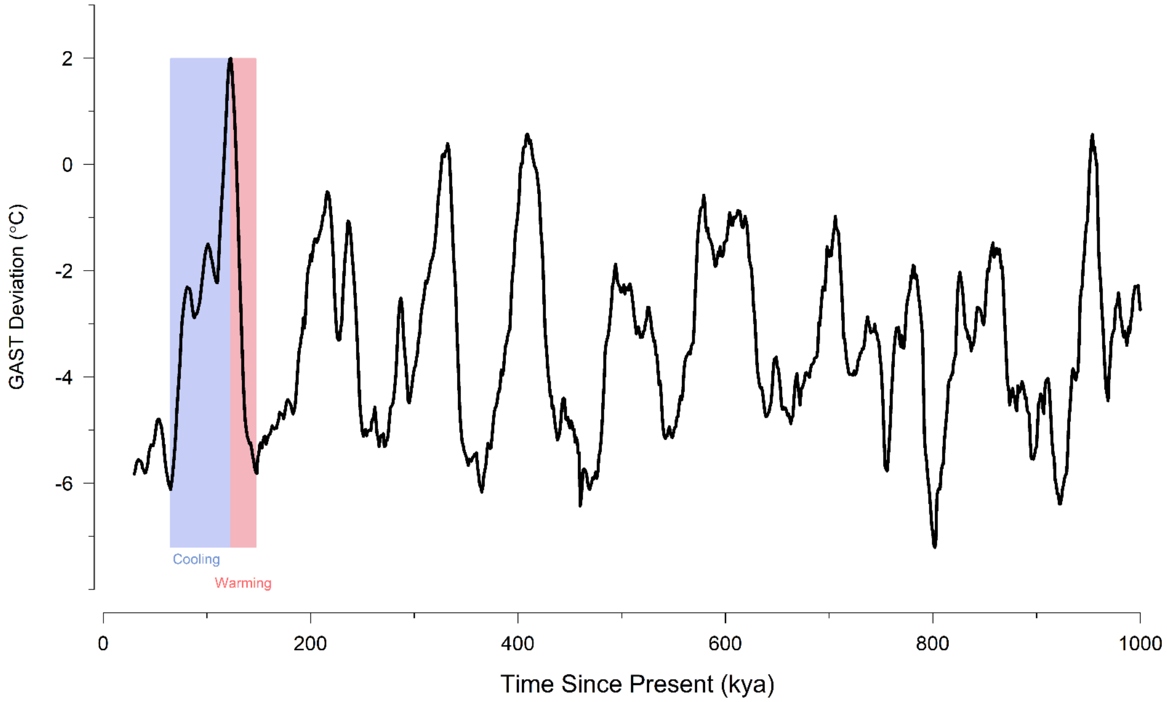

**Fig. S8.** Estimated deviation of global average surface temperature (GAST, °C) from present levels (black line, data from Snyder[51]). Blue and red bars depict focal periods of abruptly increasing (~147–123kya) and decreasing (~122–65kya) temperatures, used to calculate species-specific demographic responses during *Climate Warming* and *Climate Cooling*.

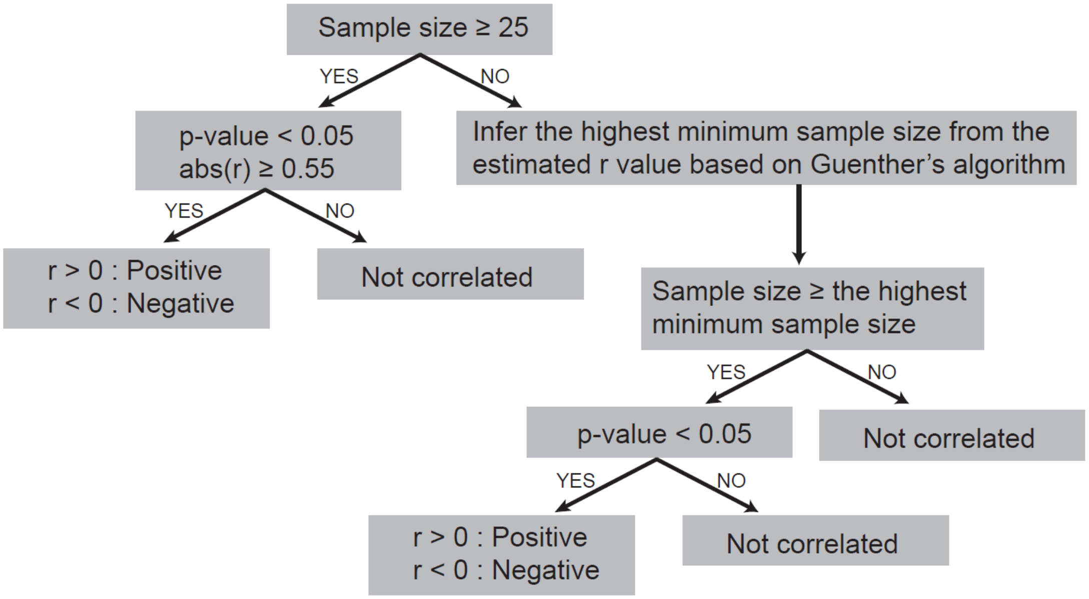

**Fig. S9.** Flow chart depicting the criteria used in quantifying demographic responses ("Increase", "Decrease", and "Unrelated") to changing climate. *r* represents Pearson's correlation coefficient.

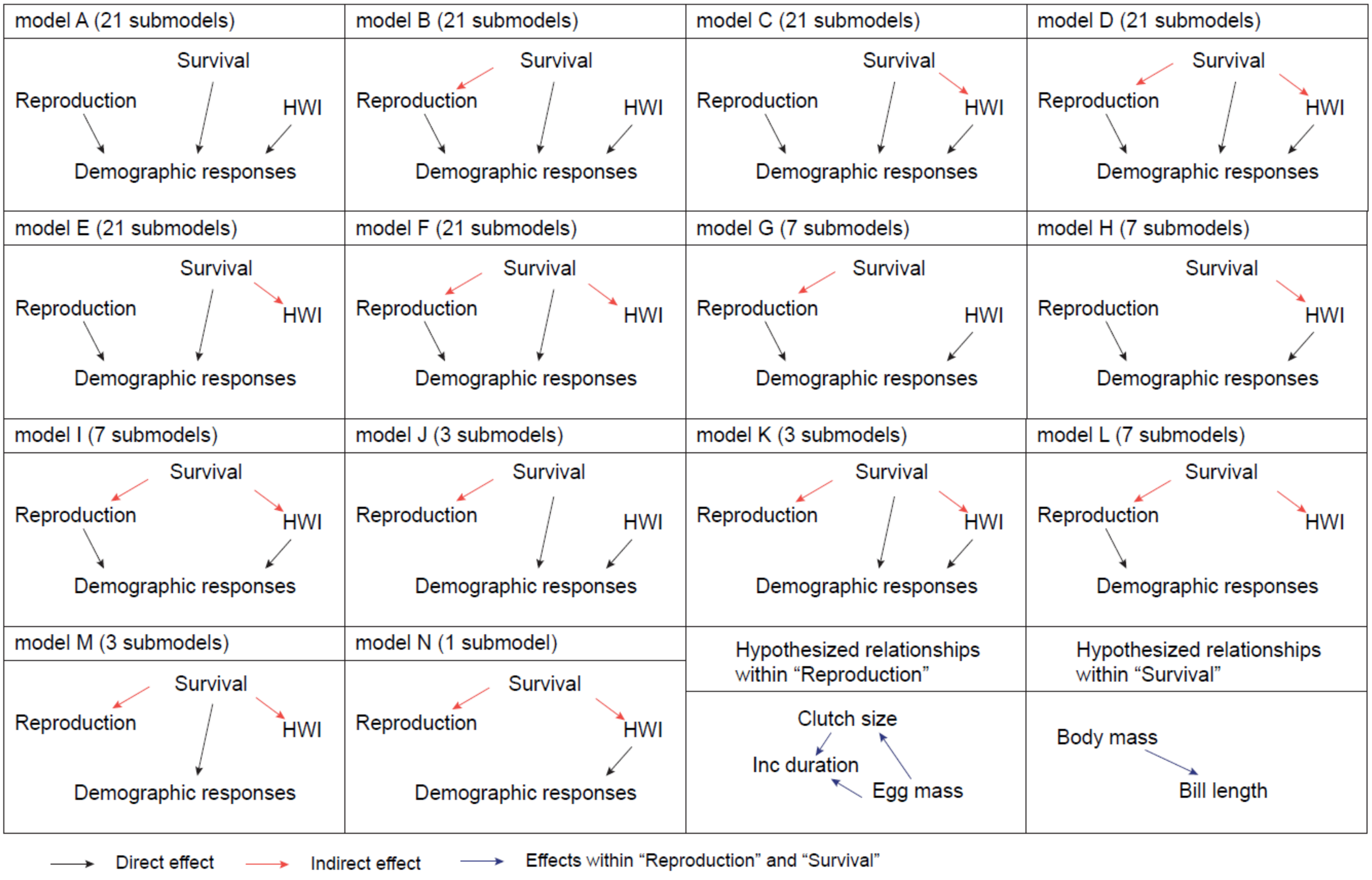

model A (21 submodels)
model B (21 submodels)
model C (21 submodels)
model D (21 submodels)
model E (21 submodels)
model F (21 submodels)
model G (7 submodels)
model H (7 submodels)
model I (7 submodels)
model J (3 submodels)
model K (3 submodels)
model L (7 submodels)
model M (3 submodels)
model N (1 submodel)
Survival
Reproduction
HWI
Demographic responses
Hypothesized relationships within “Reproduction”
Clutch size
Inc duration
Egg mass
Hypothesized relationships within “Survival”
Body mass
Bill length
Direct effect
Indirect effect
Effects within “Reproduction” and “Survival”

**Fig. S10.** 14 core models (164 models in total) shown as Directed Acyclic Graphs (DAGs) were tested in Phylogenetic Path Analysis. The “*Reproduction*” category contains three traits (egg mass, clutch size, and incubation duration) and the “*Survival*” category contains two traits (body mass and bill length), each of which have their hypothesized relationships labeled in blue. A direct effect (black) means a trait is directly causally linked to the demographic responses. An indirect effect (red) means a trait is a causal parent of other traits. Since the “*Reproduction*” and “*Survival*” categories contain multiple traits, the number of derived sub models is given in the heading next to the name of each core model.

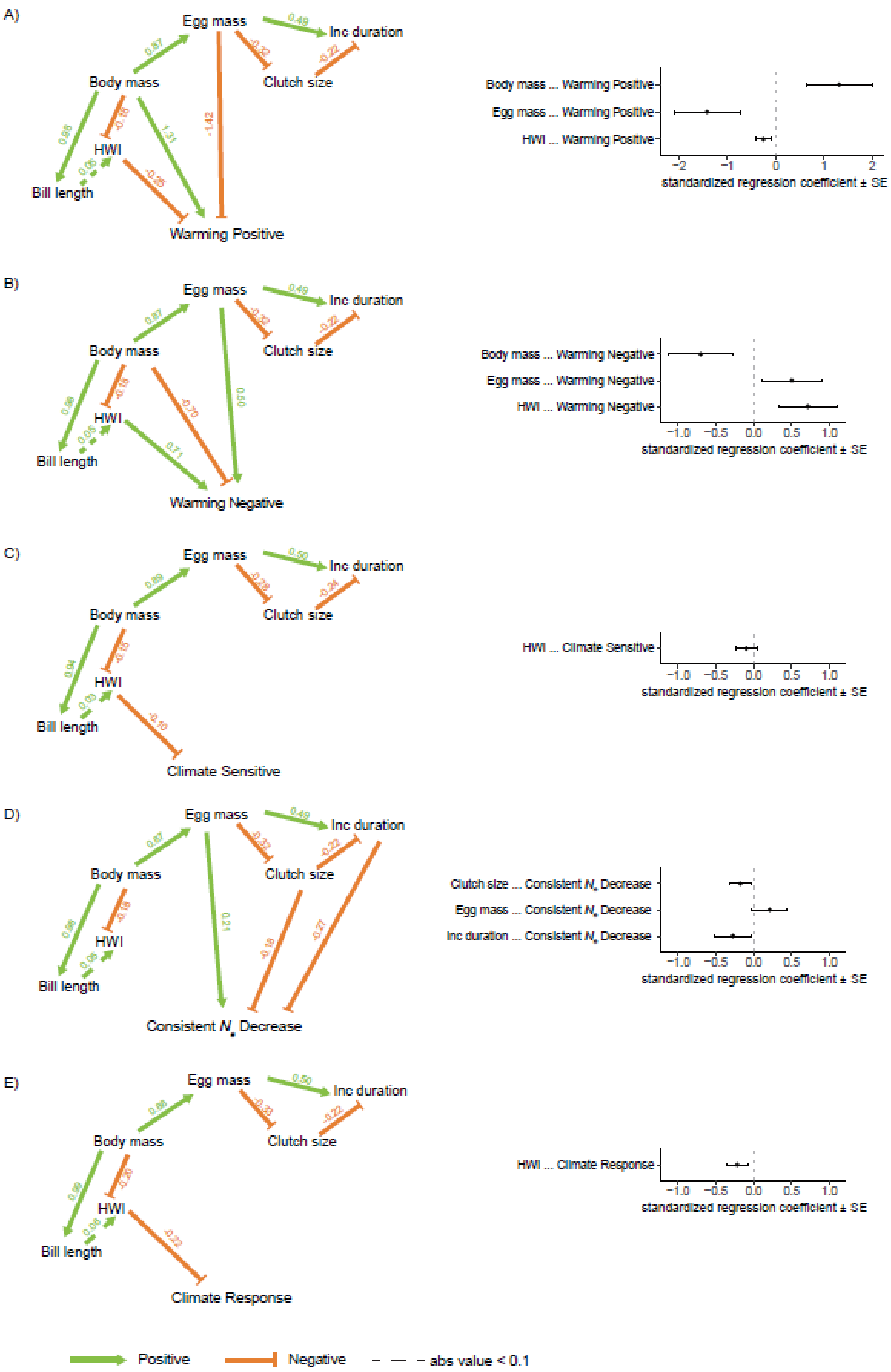

A)
Egg mass
Inc duration
Body mass
Clutch size
HWI
Bill length
Warming Positive
Body mass ... Warming Positive
Egg mass ... Warming Positive
HWI ... Warming Positive
standardized regression coefficient ± SE
B)
Egg mass
Inc duration
Body mass
Clutch size
HWI
Bill length
Warming Negative
Body mass ... Warming Negative
Egg mass ... Warming Negative
HWI ... Warming Negative
standardized regression coefficient ± SE
C)
Egg mass
Inc duration
Body mass
Clutch size
HWI
Bill length
Climate Sensitive
HWI ... Climate Sensitive
standardized regression coefficient ± SE
D)
Egg mass
Inc duration
Body mass
Clutch size
HWI
Bill length
Consistent $N_e$ Decrease
Clutch size ... Consistent $N_e$ Decrease
Egg mass ... Consistent $N_e$ Decrease
Inc duration ... Consistent $N_e$ Decrease
standardized regression coefficient ± SE
E)
Egg mass
Inc duration
Body mass
Clutch size
HWI
Bill length
Climate Response
HWI ... Climate Response
standardized regression coefficient ± SE
Positive
Negative
abs value < 0.1

**Fig. S11.** Directed acyclic graphs and corresponding standardized regression coefficients (± standard error) for the top-ranked models of five comparisons implemented via Phylogenetic Path Analysis without limiting the confidence level. A) “Warming Positive” responses versus all remaining species. B) “Warming Negative” versus all remaining species. C) species sensitive to *Climate Warming* or *Climate Cooling* versus species with consistent $N_e$ increases or decreases. D) species with consistent $N_e$ decreases under *Climate Warming* and *Climate Cooling* versus all remaining species. E) species with increasing $N_e$ tendency versus species with decreasing $N_e$ tendency under *Climate Warming* (see Supplementary Text S3). Positive paths are depicted with green arrows, while negative paths are given in orange and values above the line depict corresponding standardized regression coefficients. Dotted lines represent standardized regression coefficients less than 0.1. See table S6 for details of model outputs of Panels A-D and table S12 for details of model outputs of Panel E.

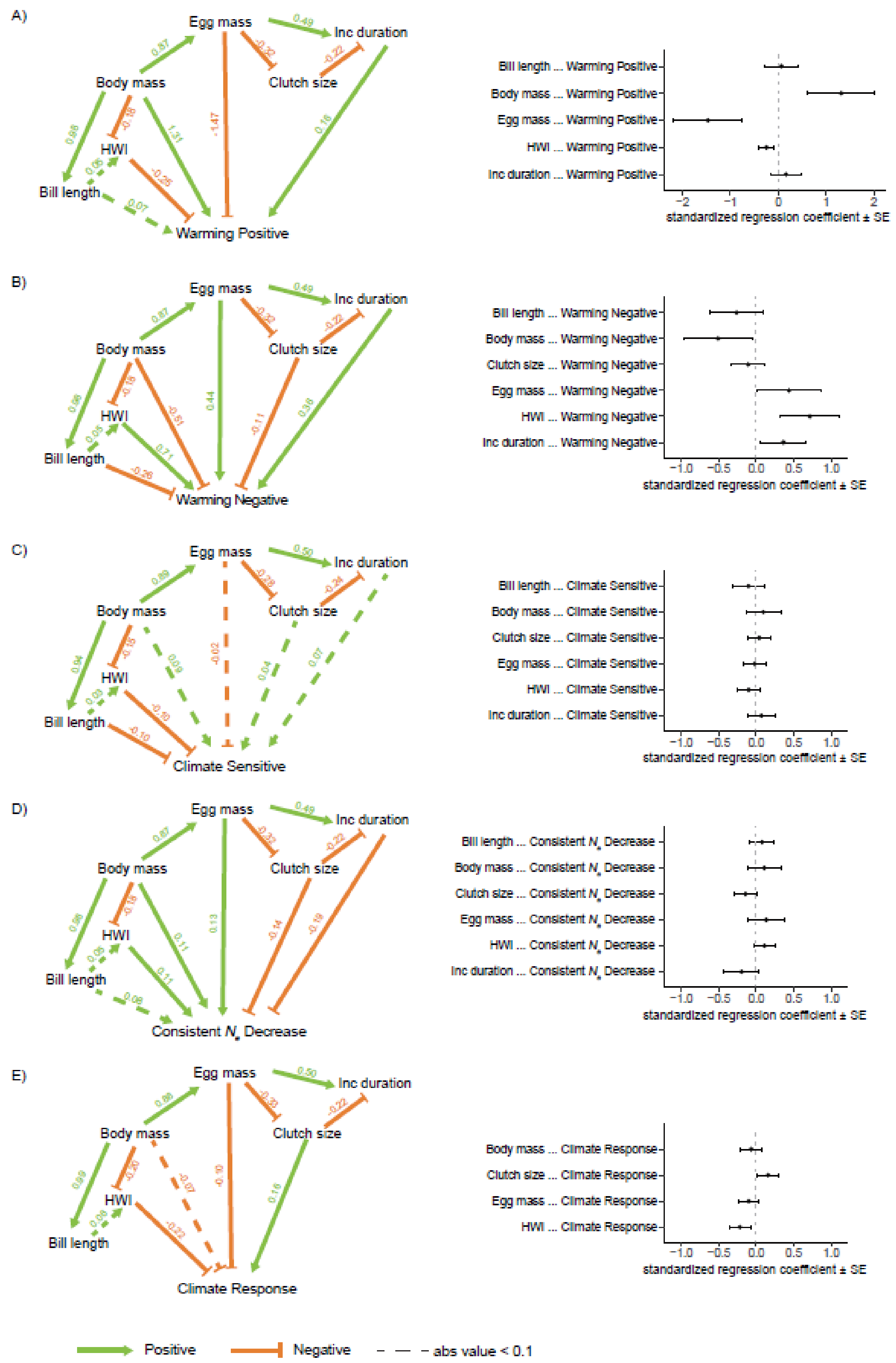

A)
Egg mass
Inc duration
Body mass
Clutch size
HWI
Bill length
Warming Positive
Bill length ... Warming Positive
Body mass ... Warming Positive
Egg mass ... Warming Positive
HWI ... Warming Positive
Inc duration ... Warming Positive
−2 −1 0 1 2
standardized regression coefficient ± SE
B)
Egg mass
Inc duration
Body mass
Clutch size
HWI
Bill length
Warming Negative
Bill length ... Warming Negative
Body mass ... Warming Negative
Clutch size ... Warming Negative
Egg mass ... Warming Negative
HWI ... Warming Negative
Inc duration ... Warming Negative
−1.0 −0.5 0.0 0.5 1.0
standardized regression coefficient ± SE
C)
Egg mass
Inc duration
Body mass
Clutch size
HWI
Bill length
Climate Sensitive
Bill length ... Climate Sensitive
Body mass ... Climate Sensitive
Clutch size ... Climate Sensitive
Egg mass ... Climate Sensitive
HWI ... Climate Sensitive
Inc duration ... Climate Sensitive
−1.0 −0.5 0.0 0.5 1.0
standardized regression coefficient ± SE
D)
Egg mass
Inc duration
Body mass
Clutch size
HWI
Bill length
Consistent $N_e$ Decrease
Bill length ... Consistent $N_e$ Decrease
Body mass ... Consistent $N_e$ Decrease
Clutch size ... Consistent $N_e$ Decrease
Egg mass ... Consistent $N_e$ Decrease
HWI ... Consistent $N_e$ Decrease
Inc duration ... Consistent $N_e$ Decrease
−1.0 −0.5 0.0 0.5 1.0
standardized regression coefficient ± SE
E)
Egg mass
Inc duration
Body mass
Clutch size
HWI
Bill length
Climate Response
Body mass ... Climate Response
Clutch size ... Climate Response
Egg mass ... Climate Response
HWI ... Climate Response
−1.0 −0.5 0.0 0.5 1.0
standardized regression coefficient ± SE
Positive
Negative
abs value < 0.1

**Fig. S12.** Directed acyclic graphs and corresponding standardized regression coefficients (± standard error) for the average best performing models (ΔCICc ≤2 from top-ranked model) of five comparisons implemented via Phylogenetic Path Analysis without limiting the confidence level. A) "Warming Positive" responses versus all remaining species. B) "Warming Negative" versus all remaining species. C) species sensitive to *Climate Warming* or *Climate Cooling* versus species with consistent $N_e$ increases or decreases. D) species with consistent $N_e$ decreases under *Climate Warming* and *Climate Cooling* versus all remaining species. E) species with increasing $N_e$ tendency versus species with decreasing $N_e$ tendency under *Climate Warming* (see Supplementary Text S3). Positive paths are depicted with green arrows, while negative paths are given in orange and values above the line depict corresponding standardized regression coefficients. Dotted lines represent standardized regression coefficients less than 0.1. See table S6 for details of model outputs of Panels A-D and table S12 for details of model outputs of Panel E.

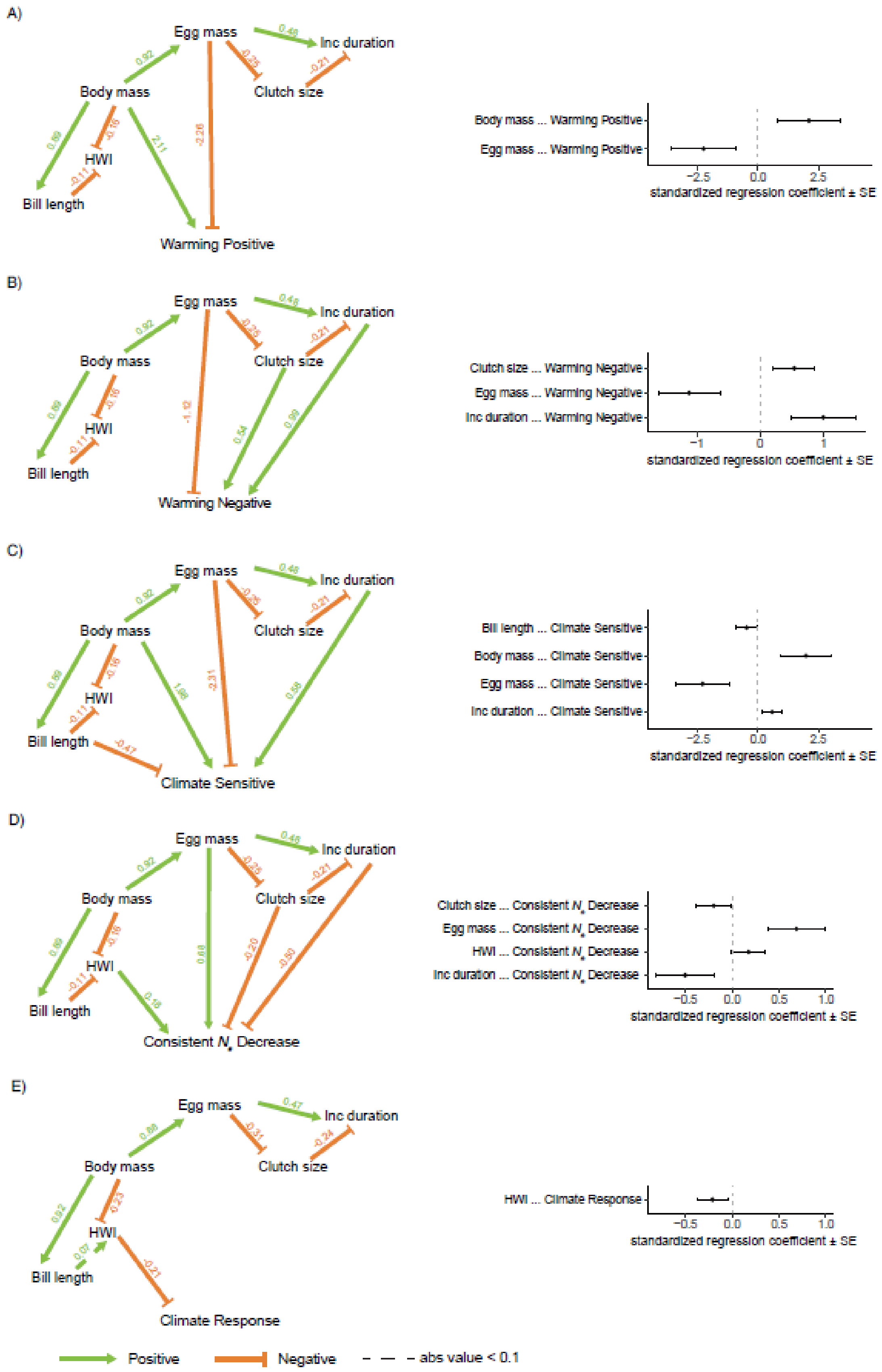
A)
Egg mass
Inc duration
Body mass
Clutch size
HWI
Bill length
Warming Positive
Body mass ... Warming Positive
Egg mass ... Warming Positive
standardized regression coefficient ± SE
B)
Warming Negative
Clutch size ... Warming Negative
Egg mass ... Warming Negative
Inc duration ... Warming Negative
C)
Climate Sensitive
Bill length ... Climate Sensitive
Body mass ... Climate Sensitive
Egg mass ... Climate Sensitive
Inc duration ... Climate Sensitive
D)
Consistent $N_e$ Decrease
Clutch size ... Consistent $N_e$ Decrease
Egg mass ... Consistent $N_e$ Decrease
HWI ... Consistent $N_e$ Decrease
Inc duration ... Consistent $N_e$ Decrease
E)
Climate Response
HWI ... Climate Response
Positive
Negative
abs value < 0.1

**Fig. S13.** Directed acyclic graphs and corresponding standardized regression coefficients (± standard error) for the top-ranked models of five comparisons implemented via Phylogenetic Path Analysis when setting the confidence level as 95%. A) “Warming Positive” responses versus all remaining species. B) “Warming Negative” versus all remaining species. C) species sensitive to *Climate Warming* or *Climate Cooling* versus species with consistent $N_e$ increases or decreases. D) species with consistent $N_e$ decreases under *Climate Warming* and *Climate Cooling* versus all remaining species. E) species with increasing $N_e$ tendency versus species with decreasing $N_e$ tendency under *Climate Warming* (see Supplementary Text S3). Positive paths are depicted with green arrows, while negative paths are given in orange and values above the line depict corresponding standardized regression coefficients. Dotted lines represent standardized regression coefficients less than 0.1. See table S7 for details of model outputs of Panels A-D and table S12 for details of model outputs of Panel E.

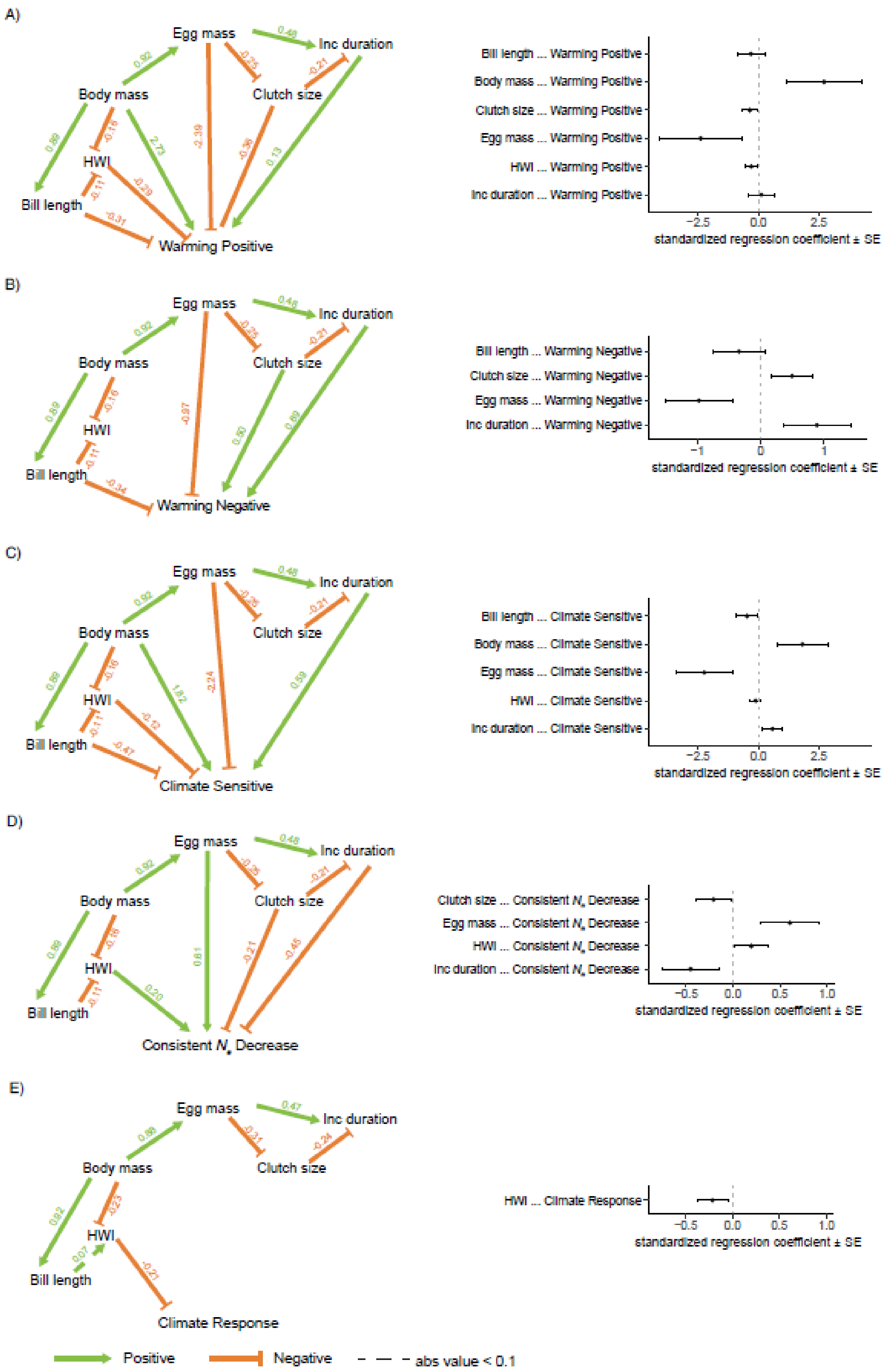
A)
Egg mass
Inc duration
Body mass
Clutch size
HWI
Bill length
Warming Positive
0.48
0.92
-0.35
-0.21
0.89
-0.16
2.73
-2.39
-0.36
0.13
-0.11
-0.29
-0.31
Bill length ... Warming Positive
Body mass ... Warming Positive
Clutch size ... Warming Positive
Egg mass ... Warming Positive
HWI ... Warming Positive
Inc duration ... Warming Positive
−2.5
0.0
2.5
standardized regression coefficient ± SE
B)
Egg mass
Inc duration
Body mass
Clutch size
HWI
Bill length
Warming Negative
0.48
0.92
-0.35
-0.21
0.89
-0.16
-0.97
0.50
0.89
-0.11
-0.34
Bill length ... Warming Negative
Clutch size ... Warming Negative
Egg mass ... Warming Negative
Inc duration ... Warming Negative
−1
0
1
standardized regression coefficient ± SE
C)
Egg mass
Inc duration
Body mass
Clutch size
HWI
Bill length
Climate Sensitive
0.48
0.92
-0.35
-0.21
0.89
-0.16
1.82
-2.24
0.59
-0.11
-0.12
-0.47
Bill length ... Climate Sensitive
Body mass ... Climate Sensitive
Egg mass ... Climate Sensitive
HWI ... Climate Sensitive
Inc duration ... Climate Sensitive
−2.5
0.0
2.5
standardized regression coefficient ± SE
D)
Egg mass
Inc duration
Body mass
Clutch size
HWI
Bill length
Consistent $N_e$ Decrease
0.48
0.92
-0.35
-0.21
0.89
-0.16
0.61
-0.21
-0.45
0.20
-0.11
Clutch size ... Consistent $N_e$ Decrease
Egg mass ... Consistent $N_e$ Decrease
HWI ... Consistent $N_e$ Decrease
Inc duration ... Consistent $N_e$ Decrease
−0.5
0.0
0.5
1.0
standardized regression coefficient ± SE
E)
Egg mass
Inc duration
Body mass
Clutch size
HWI
Bill length
Climate Response
0.47
0.88
-0.31
-0.24
0.82
-0.23
0.07
-0.21
HWI ... Climate Response
−0.5
0.0
0.5
1.0
standardized regression coefficient ± SE
Positive
Negative
abs value < 0.1

**Fig. S14.** Directed acyclic graphs and corresponding standardized regression coefficients (± standard error) for the average best performing models (ΔCICc ≤2 from top-ranked model) of five comparisons implemented via Phylogenetic Path Analysis when setting the confidence level as 95%. A) "Warming Positive" responses versus all remaining species. B) "Warming Negative" versus all remaining species. C) species sensitive to *Climate Warming* or *Climate Cooling* versus species with consistent $N_e$ increases or decreases. D) species with consistent $N_e$ decreases under *Climate Warming* and *Climate Cooling* versus all remaining species. E) species with increasing $N_e$ tendency versus species with decreasing $N_e$ tendency under *Climate Warming* (see Supplementary Text S3). Positive paths are depicted with green arrows, while negative paths are given in orange and values above the line depict corresponding standardized regression coefficients. Dotted lines represent standardized regression coefficients less than 0.1. See table S7 for details of model outputs of Panels A-D and table S12 for details of model outputs of Panel E.

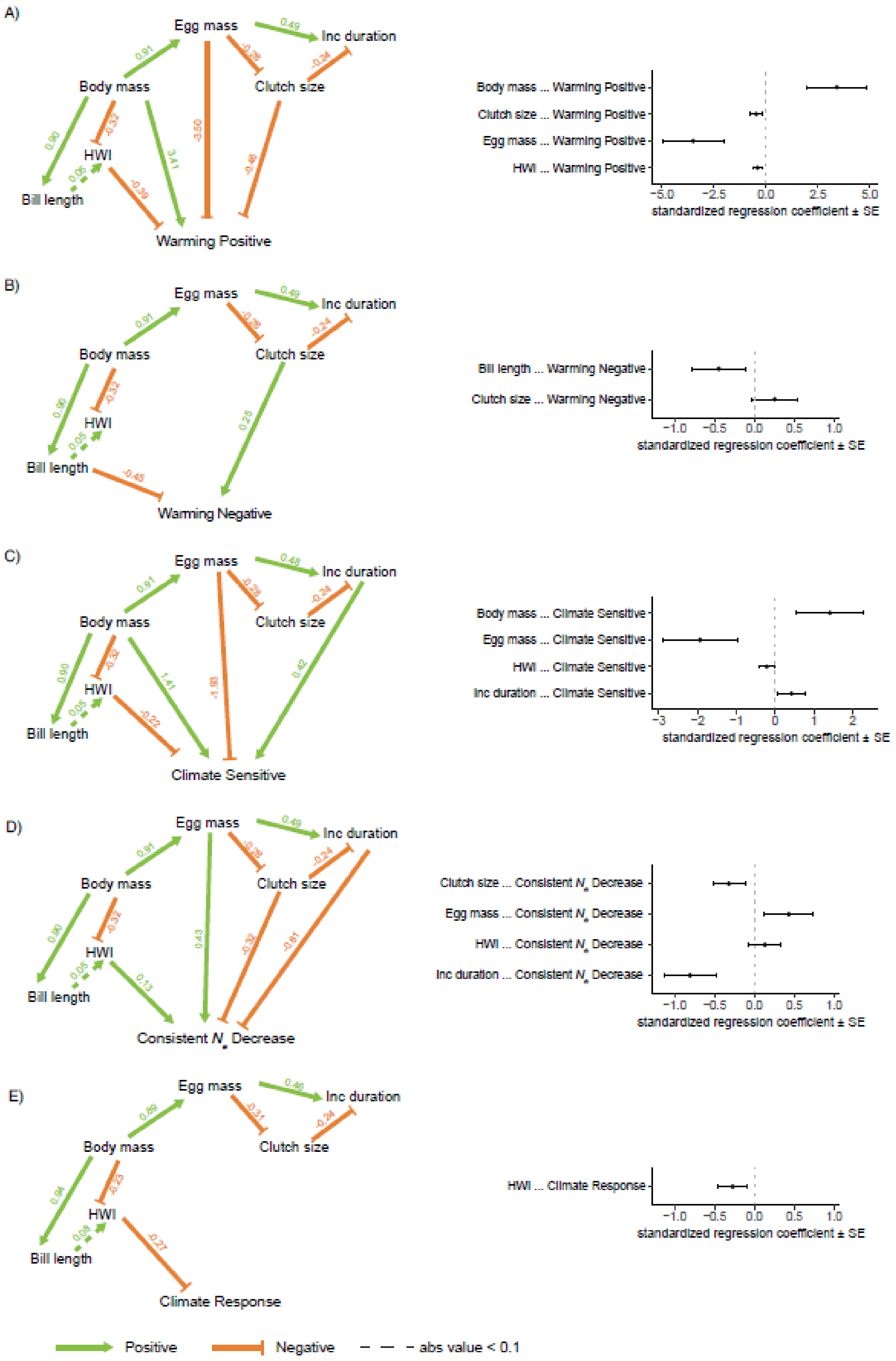
A)
Egg mass
Inc duration
Body mass
Clutch size
HWI
Bill length
Warming Positive
Body mass ... Warming Positive
Clutch size ... Warming Positive
Egg mass ... Warming Positive
HWI ... Warming Positive
−5.0
−2.5
0.0
2.5
5.0
standardized regression coefficient ± SE
B)
Egg mass
Inc duration
Body mass
Clutch size
HWI
Bill length
Warming Negative
Bill length ... Warming Negative
Clutch size ... Warming Negative
−1.0
−0.5
0.0
0.5
1.0
standardized regression coefficient ± SE
C)
Egg mass
Inc duration
Body mass
Clutch size
HWI
Bill length
Climate Sensitive
Body mass ... Climate Sensitive
Egg mass ... Climate Sensitive
HWI ... Climate Sensitive
Inc duration ... Climate Sensitive
−3
−2
−1
0
1
2
standardized regression coefficient ± SE
D)
Egg mass
Inc duration
Body mass
Clutch size
HWI
Bill length
Consistent $N_e$ Decrease
Clutch size ... Consistent $N_e$ Decrease
Egg mass ... Consistent $N_e$ Decrease
HWI ... Consistent $N_e$ Decrease
Inc duration ... Consistent $N_e$ Decrease
−1.0
−0.5
0.0
0.5
1.0
standardized regression coefficient ± SE
E)
Egg mass
Inc duration
Body mass
Clutch size
HWI
Bill length
Climate Response
HWI ... Climate Response
−1.0
−0.5
0.0
0.5
1.0
standardized regression coefficient ± SE
Positive
Negative
abs value < 0.1

**Fig. S15.** Directed acyclic graphs and corresponding standardized regression coefficients (± standard error) for the top-ranked models of five comparisons implemented via Phylogenetic Path Analysis when setting the confidence level as 90%. A) "Warming Positive" responses versus all remaining species. B) "Warming Negative" versus all remaining species. C) species sensitive to *Climate Warming* or *Climate Cooling* versus species with consistent $N_e$ increases or decreases. D) species with consistent $N_e$ decreases under *Climate Warming* and *Climate Cooling* versus all remaining species. E) species with increasing $N_e$ tendency versus species with decreasing $N_e$ tendency under *Climate Warming* (see Supplementary Text S3). Positive paths are depicted with green arrows, while negative paths are given in orange and values above the line depict corresponding standardized regression coefficients. Dotted lines represent standardized regression coefficients less than 0.1. See table S8 for details of model outputs of Panels A-D and table S12 for details of model outputs of Panel E.

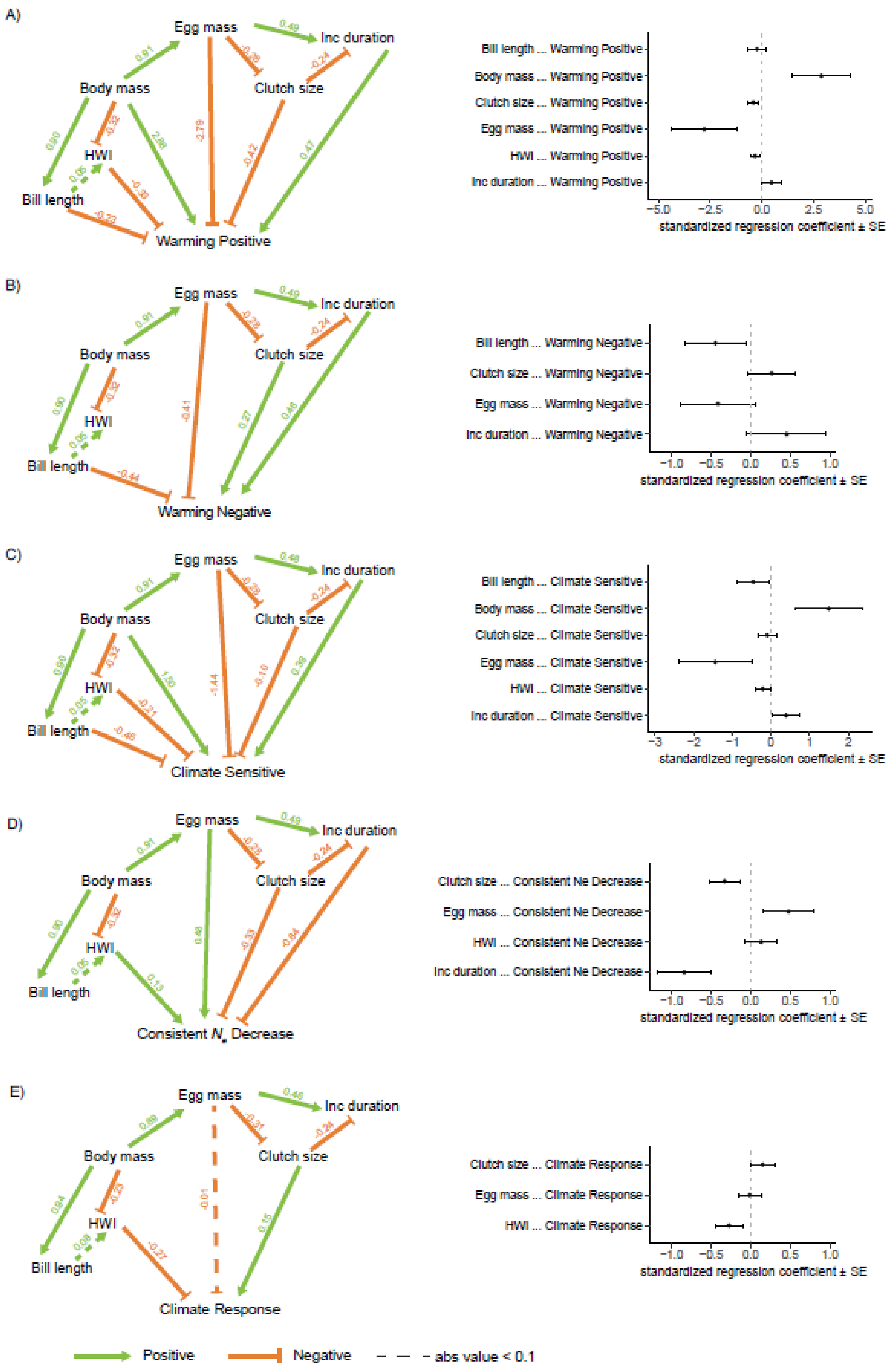

A)
Egg mass
Inc duration
Body mass
Clutch size
HWI
Bill length
Warming Positive
0.49
0.91
-0.28
-0.24
0.90
-0.32
2.86
-2.79
-0.42
0.47
0.05
-0.35
-0.23
Bill length ... Warming Positive
Body mass ... Warming Positive
Clutch size ... Warming Positive
Egg mass ... Warming Positive
HWI ... Warming Positive
Inc duration ... Warming Positive
−5.0
−2.5
0.0
2.5
5.0
standardized regression coefficient ± SE
B)
Egg mass
Inc duration
Body mass
Clutch size
HWI
Bill length
Warming Negative
0.49
0.91
-0.28
-0.24
0.90
-0.32
-0.41
0.27
0.46
0.05
-0.44
Bill length ... Warming Negative
Clutch size ... Warming Negative
Egg mass ... Warming Negative
Inc duration ... Warming Negative
−1.0
−0.5
0.0
0.5
1.0
standardized regression coefficient ± SE
C)
Egg mass
Inc duration
Body mass
Clutch size
HWI
Bill length
Climate Sensitive
0.48
0.91
-0.28
-0.24
0.90
-0.32
1.50
-1.44
-0.10
0.39
0.05
-0.21
-0.46
Bill length ... Climate Sensitive
Body mass ... Climate Sensitive
Clutch size ... Climate Sensitive
Egg mass ... Climate Sensitive
HWI ... Climate Sensitive
Inc duration ... Climate Sensitive
−3
−2
−1
0
1
2
standardized regression coefficient ± SE
D)
Egg mass
Inc duration
Body mass
Clutch size
HWI
Bill length
Consistent $N_e$ Decrease
0.49
0.91
-0.28
-0.24
0.90
-0.32
0.48
-0.33
-0.84
0.05
0.13
Clutch size ... Consistent Ne Decrease
Egg mass ... Consistent Ne Decrease
HWI ... Consistent Ne Decrease
Inc duration ... Consistent Ne Decrease
−1.0
−0.5
0.0
0.5
1.0
standardized regression coefficient ± SE
E)
Egg mass
Inc duration
Body mass
Clutch size
HWI
Bill length
Climate Response
0.46
0.89
-0.31
-0.24
0.84
-0.23
-0.01
0.15
0.08
-0.27
Clutch size ... Climate Response
Egg mass ... Climate Response
HWI ... Climate Response
−1.0
−0.5
0.0
0.5
1.0
standardized regression coefficient ± SE
Positive
Negative
abs value < 0.1

**Fig. S16.** Directed acyclic graphs and corresponding standardized regression coefficients (± standard error) for the average best performing models (ΔCICc ≤2 from top-ranked model) of five comparisons implemented via Phylogenetic Path Analysis when setting the confidence level as 90%. A) “Warming Positive” responses versus all remaining species. B) “Warming Negative” versus all remaining species. C) species sensitive to *Climate Warming* or *Climate Cooling* versus species with consistent $N_e$ increases or decreases. D) species with consistent $N_e$ decreases under *Climate Warming* and *Climate Cooling* versus all remaining species. E) species with increasing $N_e$ tendency versus species with decreasing $N_e$ tendency under *Climate Warming* (see Supplementary Text S3). Positive paths are depicted with green arrows, while negative paths are given in orange and values above the line depict corresponding standardized regression coefficients. Dotted lines represent standardized regression coefficients less than 0.1. See table S8 for details of model outputs of Panels A-D and table S12 for details of model outputs of Panel E.

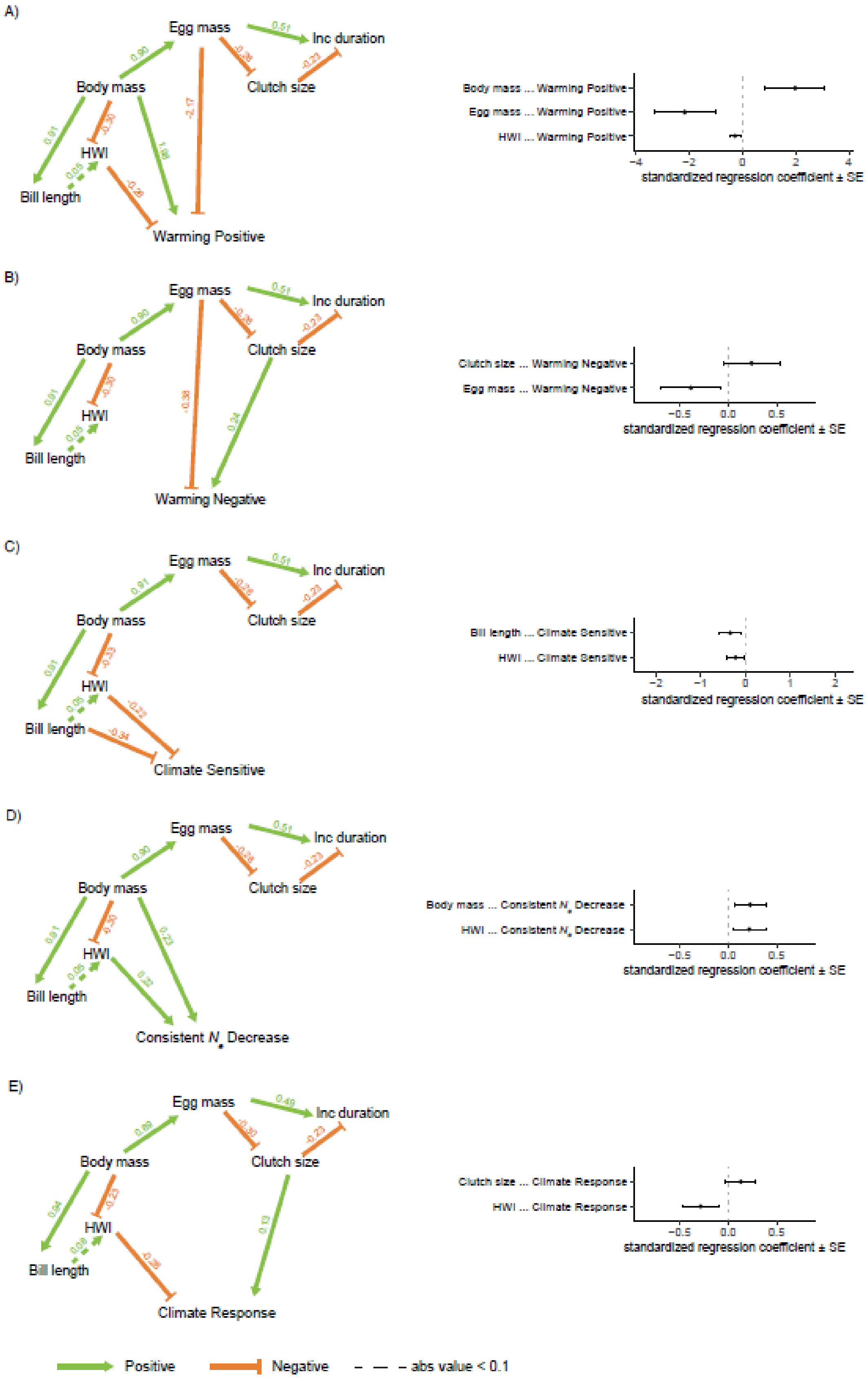

A)
Egg mass
Inc duration
Body mass
Clutch size
HWI
Bill length
Warming Positive
Body mass ... Warming Positive
Egg mass ... Warming Positive
HWI ... Warming Positive
standardized regression coefficient ± SE
B)
Egg mass
Inc duration
Body mass
Clutch size
HWI
Bill length
Warming Negative
Clutch size ... Warming Negative
Egg mass ... Warming Negative
standardized regression coefficient ± SE
C)
Egg mass
Inc duration
Body mass
Clutch size
HWI
Bill length
Climate Sensitive
Bill length ... Climate Sensitive
HWI ... Climate Sensitive
standardized regression coefficient ± SE
D)
Egg mass
Inc duration
Body mass
Clutch size
HWI
Bill length
Consistent $N_e$ Decrease
Body mass ... Consistent $N_e$ Decrease
HWI ... Consistent $N_e$ Decrease
standardized regression coefficient ± SE
E)
Egg mass
Inc duration
Body mass
Clutch size
HWI
Bill length
Climate Response
Clutch size ... Climate Response
HWI ... Climate Response
standardized regression coefficient ± SE
Positive
Negative
abs value < 0.1

**Fig. S17.** Directed acyclic graphs and corresponding standardized regression coefficients (± standard error) for the top-ranked models of five comparisons implemented via Phylogenetic Path Analysis when setting the confidence level as 85%. A) "Warming Positive" responses versus all remaining species. B) "Warming Negative" versus all remaining species. C) species sensitive to *Climate Warming* or *Climate Cooling* versus species with consistent $N_e$ increases or decreases. D) species with consistent $N_e$ decreases under *Climate Warming* and *Climate Cooling* versus all remaining species. E) species with increasing $N_e$ tendency versus species with decreasing $N_e$ tendency under *Climate Warming* (see Supplementary Text S3). Positive paths are depicted with green arrows, while negative paths are given in orange and values above the line depict corresponding standardized regression coefficients. Dotted lines represent standardized regression coefficients less than 0.1. See table S9 for details of model outputs of Panels A-D and table S12 for details of model outputs of Panel E.

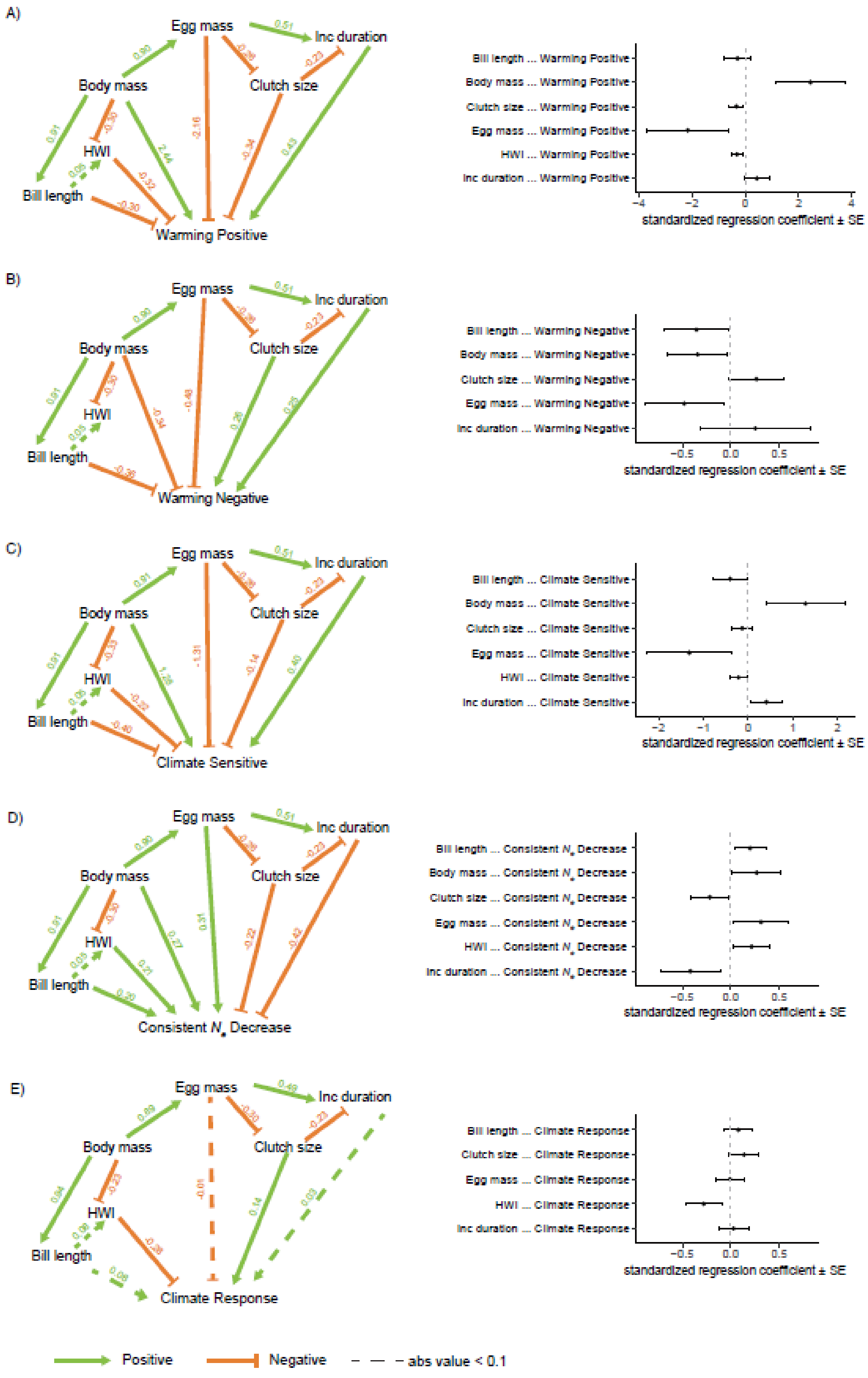
A)
Egg mass
Inc duration
Body mass
Clutch size
HWI
Bill length
Warming Positive
Bill length ... Warming Positive
Body mass ... Warming Positive
Clutch size ... Warming Positive
Egg mass ... Warming Positive
HWI ... Warming Positive
Inc duration ... Warming Positive
standardized regression coefficient ± SE
B)
Egg mass
Inc duration
Body mass
Clutch size
HWI
Bill length
Warming Negative
Bill length ... Warming Negative
Body mass ... Warming Negative
Clutch size ... Warming Negative
Egg mass ... Warming Negative
Inc duration ... Warming Negative
standardized regression coefficient ± SE
C)
Egg mass
Inc duration
Body mass
Clutch size
HWI
Bill length
Climate Sensitive
Bill length ... Climate Sensitive
Body mass ... Climate Sensitive
Clutch size ... Climate Sensitive
Egg mass ... Climate Sensitive
HWI ... Climate Sensitive
Inc duration ... Climate Sensitive
standardized regression coefficient ± SE
D)
Egg mass
Inc duration
Body mass
Clutch size
HWI
Bill length
Consistent $N_e$ Decrease
Bill length ... Consistent $N_e$ Decrease
Body mass ... Consistent $N_e$ Decrease
Clutch size ... Consistent $N_e$ Decrease
Egg mass ... Consistent $N_e$ Decrease
HWI ... Consistent $N_e$ Decrease
Inc duration ... Consistent $N_e$ Decrease
standardized regression coefficient ± SE
E)
Egg mass
Inc duration
Body mass
Clutch size
HWI
Bill length
Climate Response
Bill length ... Climate Response
Clutch size ... Climate Response
Egg mass ... Climate Response
HWI ... Climate Response
Inc duration ... Climate Response
standardized regression coefficient ± SE
Positive
Negative
abs value < 0.1

**Fig. S18.** Directed acyclic graphs and corresponding standardized regression coefficients (± standard error) for the average best performing models (ΔCICc ≤2 from top-ranked model) of five comparisons implemented via Phylogenetic Path Analysis when setting the confidence level as 85%. A) "Warming Positive" responses versus all remaining species. B) "Warming Negative" versus all remaining species. C) species sensitive to *Climate Warming* or *Climate Cooling* versus species with consistent $N_e$ increases or decreases. D) species with consistent $N_e$ decreases under *Climate Warming* and *Climate Cooling* versus all remaining species. E) species with increasing $N_e$ tendency versus species with decreasing $N_e$ tendency under *Climate Warming* (see Supplementary Text S3). Positive paths are depicted with green arrows, while negative paths are given in orange and values above the line depict corresponding standardized regression coefficients. Dotted lines standardized represent regression coefficients less than 0.1. See table S9 for details of model outputs of Panels A-D and table S12 for details of model outputs of Panel E.

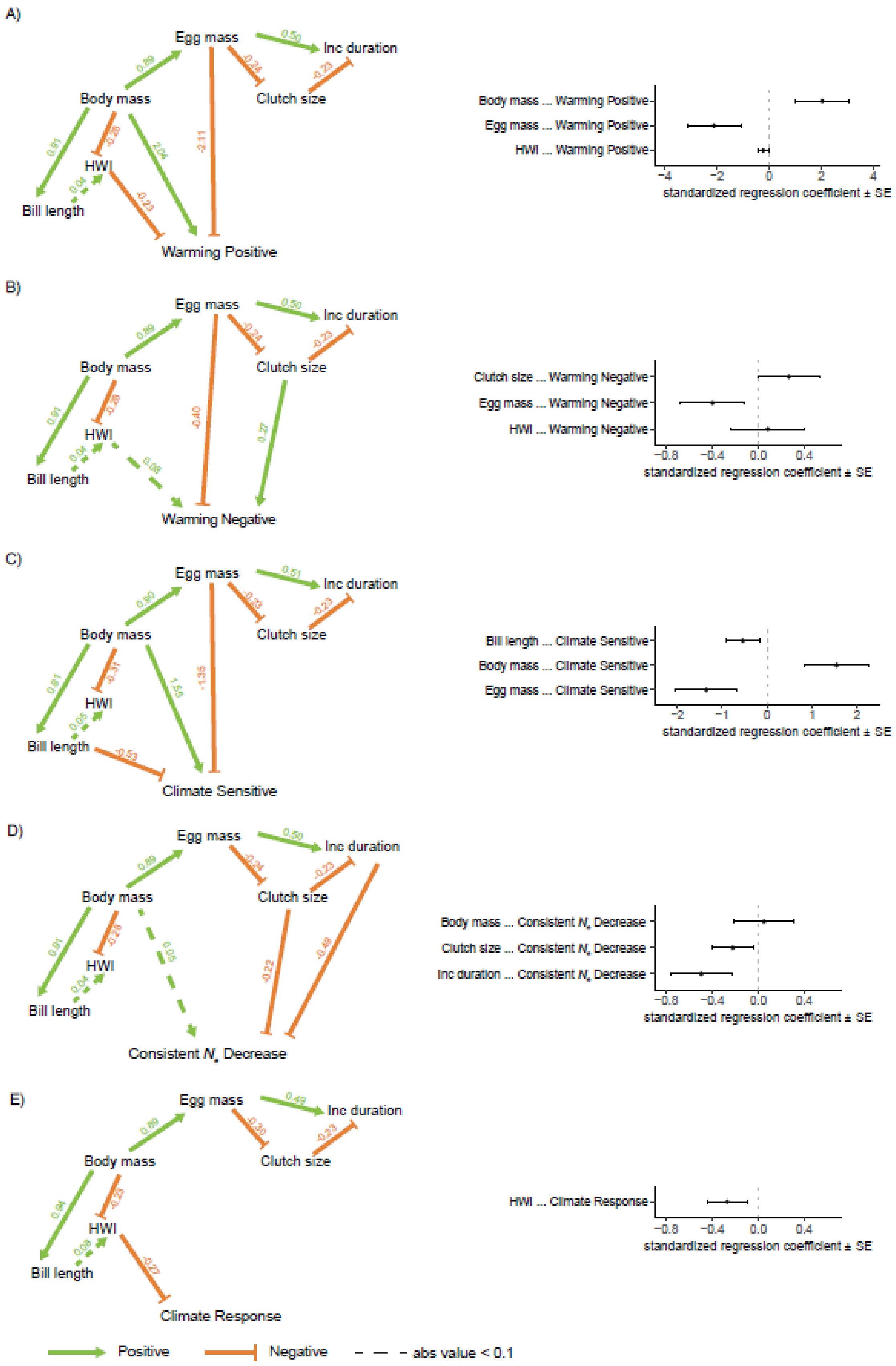

A)
Egg mass
Inc duration
Body mass
Clutch size
HWI
Bill length
Warming Positive
Body mass ... Warming Positive
Egg mass ... Warming Positive
HWI ... Warming Positive
standardized regression coefficient ± SE
B)
Warming Negative
Clutch size ... Warming Negative
Egg mass ... Warming Negative
HWI ... Warming Negative
C)
Climate Sensitive
Bill length ... Climate Sensitive
Body mass ... Climate Sensitive
Egg mass ... Climate Sensitive
D)
Consistent N, Decrease
Body mass ... Consistent N, Decrease
Clutch size ... Consistent N, Decrease
Inc duration ... Consistent N, Decrease
E)
Climate Response
HWI ... Climate Response
Positive
Negative
abs value < 0.1

730

**Fig. S19.** Directed acyclic graphs and corresponding standardized regression coefficients (± standard error) for the top-ranked models of five comparisons implemented via Phylogenetic Path Analysis when setting the confidence level as 80%. A) “Warming Positive” responses versus all remaining species. B) “Warming Negative” versus all remaining species. C) species sensitive to *Climate Warming* or *Climate Cooling* versus species with consistent $N_e$ increases or decreases. D) species with consistent $N_e$ decreases under *Climate Warming* and *Climate Cooling* versus all remaining species. E) species with increasing $N_e$ tendency versus species with decreasing $N_e$ tendency under *Climate Warming* (see Supplementary Text S3). Positive paths are depicted with green arrows, while negative paths are given in orange and values above the line depict corresponding standardized regression coefficients. Dotted lines represent standardized regression coefficients less than 0.1. See table S10 for details of model outputs of Panels A-D and table S12 for details of model outputs of Panel E.

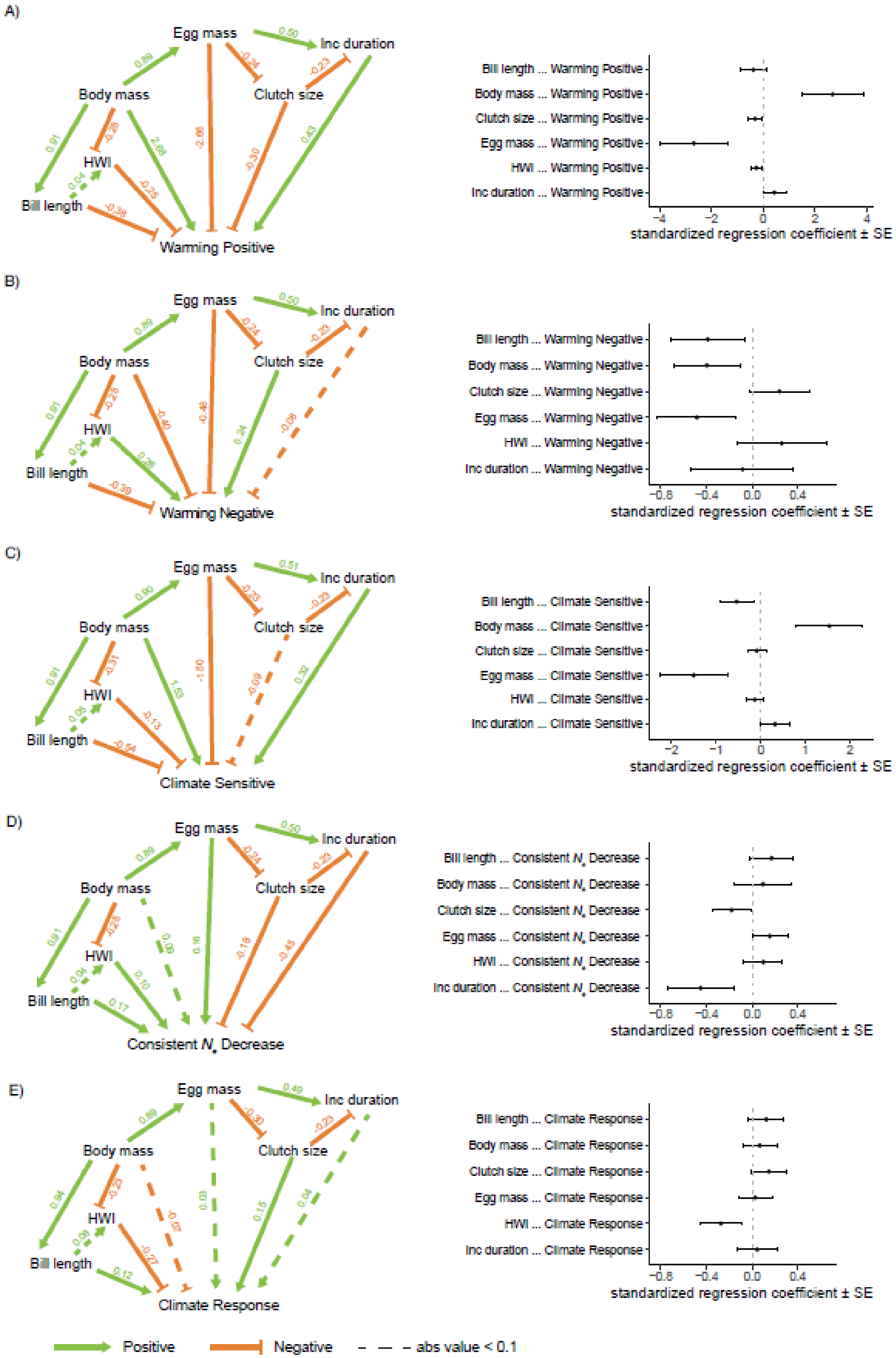
A)
Egg mass
Inc duration
Body mass
Clutch size
HWI
Bill length
Warming Positive
Bill length ... Warming Positive
Body mass ... Warming Positive
Clutch size ... Warming Positive
Egg mass ... Warming Positive
HWI ... Warming Positive
Inc duration ... Warming Positive
standardized regression coefficient ± SE
B)
Egg mass
Inc duration
Body mass
Clutch size
HWI
Bill length
Warming Negative
Bill length ... Warming Negative
Body mass ... Warming Negative
Clutch size ... Warming Negative
Egg mass ... Warming Negative
HWI ... Warming Negative
Inc duration ... Warming Negative
standardized regression coefficient ± SE
C)
Egg mass
Inc duration
Body mass
Clutch size
HWI
Bill length
Climate Sensitive
Bill length ... Climate Sensitive
Body mass ... Climate Sensitive
Clutch size ... Climate Sensitive
Egg mass ... Climate Sensitive
HWI ... Climate Sensitive
Inc duration ... Climate Sensitive
standardized regression coefficient ± SE
D)
Egg mass
Inc duration
Body mass
Clutch size
HWI
Bill length
Consistent $N_e$ Decrease
Bill length ... Consistent $N_e$ Decrease
Body mass ... Consistent $N_e$ Decrease
Clutch size ... Consistent $N_e$ Decrease
Egg mass ... Consistent $N_e$ Decrease
HWI ... Consistent $N_e$ Decrease
Inc duration ... Consistent $N_e$ Decrease
standardized regression coefficient ± SE
E)
Egg mass
Inc duration
Body mass
Clutch size
HWI
Bill length
Climate Response
Bill length ... Climate Response
Body mass ... Climate Response
Clutch size ... Climate Response
Egg mass ... Climate Response
HWI ... Climate Response
Inc duration ... Climate Response
standardized regression coefficient ± SE
Positive
Negative
abs value < 0.1

**Fig. S20.** Directed acyclic graphs and corresponding standardized regression coefficients (± standard error) for the average best performing models (ΔCICc ≤2 from top-ranked model) of five comparisons implemented via Phylogenetic Path Analysis when setting the confidence level as 80%. A) "Warming Positive" responses versus all remaining species. B) "Warming Negative" versus all remaining species. C) species sensitive to *Climate Warming* or *Climate Cooling* versus species with consistent $N_e$ increases or decreases. D) species with consistent $N_e$ decreases under *Climate Warming* and *Climate Cooling* versus all remaining species. E) species with increasing $N_e$ tendency versus species with decreasing $N_e$ tendency under *Climate Warming* (see Supplementary Text S3). Positive paths are depicted with green arrows, while negative paths are given in orange and values above the line depict corresponding standardized regression coefficients. Dotted lines represent standardized regression coefficients less than 0.1. See table S10 for details of model outputs of Panels A-D and table S12 for details of model outputs of Panel E.

757

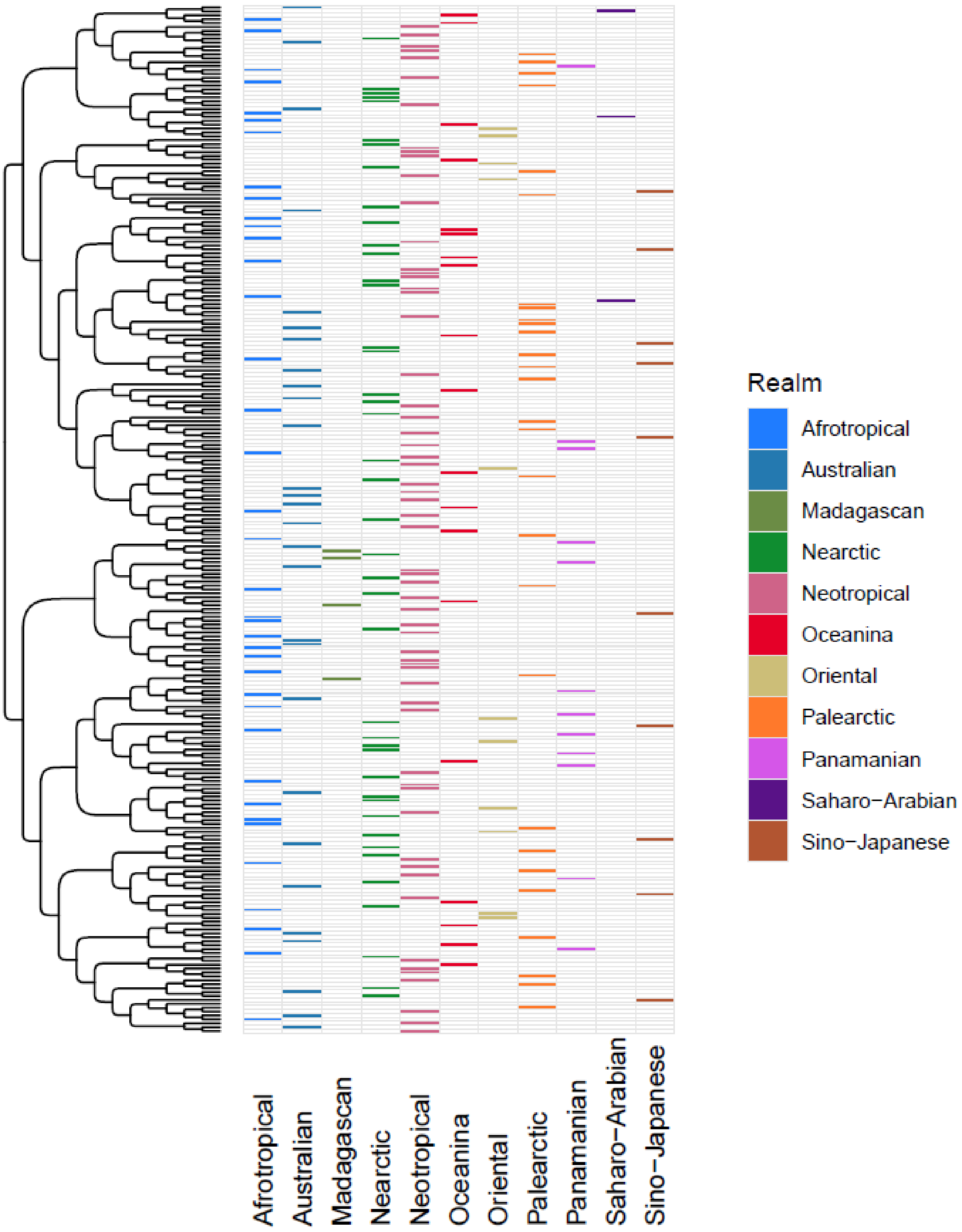
Realm
Afrotropical
Australian
Madagascan
Nearctic
Neotropical
Oceanina
Oriental
Palearctic
Panamanian
Saharo−Arabian
Sino−Japanese

**Fig. S21.** The clustering dendrogram presented in Fig 1A (at k = 7), with each species' zoogeographic realm identified. Each column along the bottom represents one of the 11 major zoogeographic realms[28], and species belonging to each realm are indicated in the corresponding color.

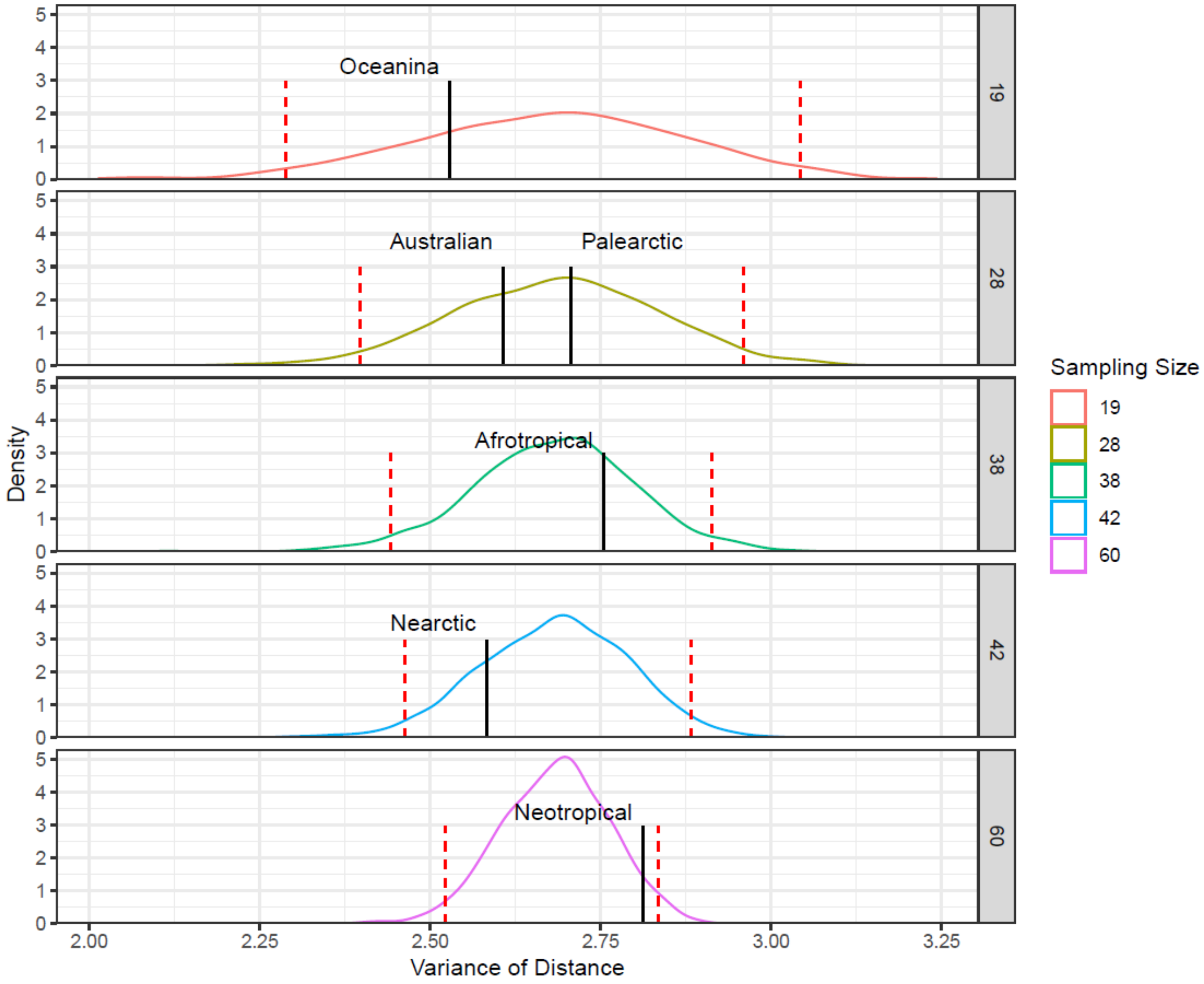

**Fig. S22**. The observed variance of distance values for six realms with n ≥ 19 species (black vertical lines) and background variances generated via randomly sampling the same number of species. For each panel, curves represent the distribution of variances generated from 1000 sampling permutations, and red dotted lines represent the 95$^{th}$ percentiles.

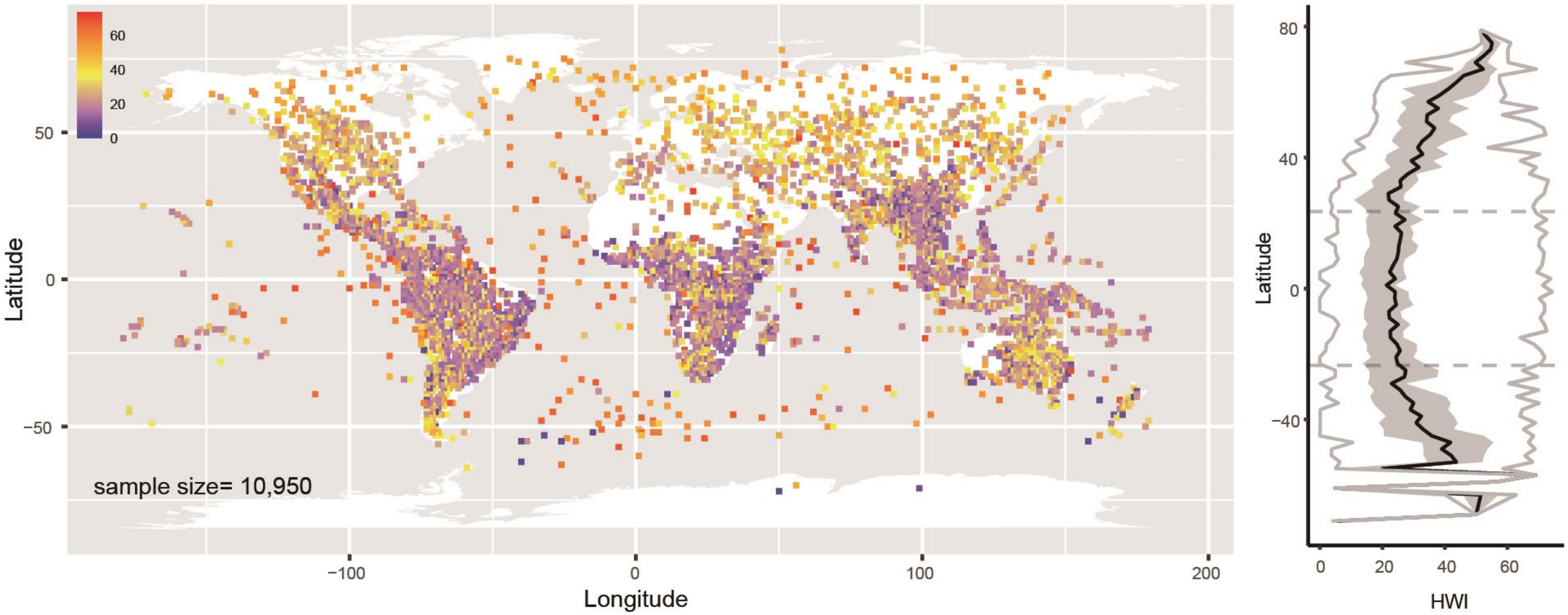

**Fig S23.** Contemporary global distribution of HWI for n = 10,950 avian species. Left: the median HWI of species within each grid cell (1° scale). Right: the HWI for each species within latitudinal bands (5° scale). The black line represents the median value, while the grey shading represents the interquartile range, and two grey lines represent the minimum and maximum of values. The dashed horizontal lines are S23.5° and N23.5°, representing the regional boundaries of the tropics.

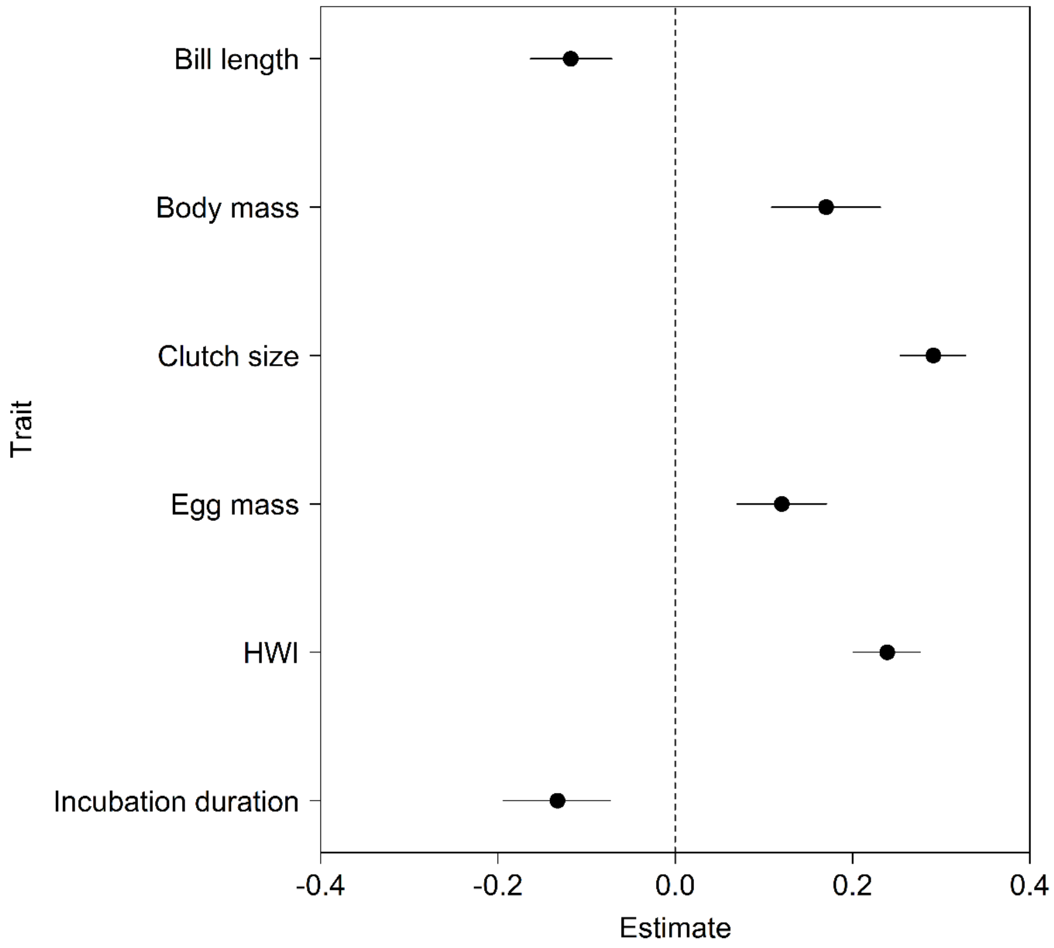

**Fig. S24**. Effects plot from linear model quantifying the relative influence of each key morphological/life-history trait on the latitude (absolute mean geographic centroid of breeding/resident range) of 2745 avian species. All possible combinations of the six predictor variables were tested, but results from the global model (*$R^2$* = 0.17) was unequivocally the most parsimonious (all other models ΔAIC > 16) and so are presented here. Points represent parameter estimates while whiskers depict 95% CIs. The dotted line represents a parameter estimate of zero, and effects are considered significant if CIs do not overlap zero.

### References

30. McKenna, A. *et al*. The Genome Analysis Toolkit: a MapReduce framework for analyzing next-generation DNA sequencing data. *Genome Res.* **20**, 1297–1303 (2010).

31. Nadachowska-Brzyska, K., Li, C., Smeds, L., Zhang, G. & Ellegren, H. Temporal dynamics of avian populations during Pleistocene revealed by whole-genome sequences. *Curr. Biol.* **25**, 1375–1380 (2015).

32. Jarvis, E. D. *et al.* Whole-genome analyses resolve early branches in the tree of life of modern birds. *Science* **346**, 1320–1331 (2014).

33. Rokas, A., Williams, B. L., King, N. & Carroll, S. B. Genome-scale approaches to resolving incongruence in molecular phylogenies. *Nature* **425**, 798–804 (2003).

34. Wolf, Y. I., Rogozin, I. B., Grishin, N. V. & Koonin, E. V. Genome trees and the tree of life. *Trends Genet.* **18**, 472–479 (2002).

35. Yang, Z. PAML: a program package for phylogenetic analysis by maximum likelihood. *Comput. Appl. Biosci.* **13**, 555–556 (1997).

36. Forest, F. Calibrating the Tree of Life: fossils, molecules and evolutionary timescales. *Ann. Bot.* **104**, 789–794 (2009).

37. Magallón, S. A. Dating Lineages: Molecular and paleontological approaches to the temporal framework of clades. *Int. J. Plant Sci.* **165**, S7–S21 (2004).

38. Nadachowska-Brzyska, K., Burri, R., Smeds, L. & Ellegren, H. PSMC analysis of effective population sizes in molecular ecology and its application to black-and-white Ficedula flycatchers. *Mol. Ecol.* **25**, 1058–1072 (2016).

39. Bird, J. P. *et al.* Generation lengths of the world's birds and their implications for extinction risk. *Cons. Biol.* **34**, 1252–1261 (2020).

40. Patton, A. H. *et al.* Contemporary demographic reconstruction methods are robust to genome assembly quality: A case study in Tasmanian devils. *Mol. Biol. Evol.* **36**, 2906–2921 (2019).

41. Liu, X. & Fu, Y.-X. Exploring population size changes using SNP frequency spectra. *Nat. Genet.* **47**, 555–559 (2015).

42. Terhorst, J. & Song, Y. S. Fundamental limits on the accuracy of demographic inference based on the sample frequency spectrum. *Proc. Natl. Acad. Sci. U.S.A.* **112**, 7677–7682 (2015).

43. Terhorst, J., Kamm, J. A. & Song, Y. S. Robust and scalable inference of population history from hundreds of unphased whole genomes. *Nat. Genet.* **49**, 303–309 (2017).

44. Lapierre, M., Lambert, A. & Achaz, G. Accuracy of demographic inferences from the site frequency spectrum: the case of the Yoruba population. *Genetics* **206**, 439–449 (2017).

45. Mazet, O., Rodríguez, W., Grusea, S., Boitard, S. & Chikhi, L. On the importance of being structured: instantaneous coalescence rates and human evolution—lessons for ancestral population size inference? *Heredity* **116**, 362–371 (2016).

46. Chikhi, L. *et al.* The IICR (inverse instantaneous coalescence rate) as a summary of genomic diversity: insights into demographic inference and model choice. *Heredity* **120**, 13–24 (2018).

47. Teixeira, H. *et al.* Impact of model assumptions on demographic inferences: the case study of two sympatric mouse lemurs in northwestern Madagascar. *BMC Ecol. Evol.* **21**, 197 (2021).

48. Galili, T. dendextend: an R package for visualizing, adjusting and comparing trees of hierarchical clustering. *Bioinformatics* **31**, 3718–3720 (2015).

49. Smith, M. R. TreeDist: Distances Between Phylogenetic Trees. R package version 2.5.0 (2020) doi:10.5281/ZENODO.3528124.

50. Paleo-López, R. *et al.* A phylogenetic analysis of macroevolutionary patterns in fermentative yeasts. *Ecol. Evol.* **6**, 3851–3861 (2016).

51. Snyder, C. W. Evolution of global temperature over the past two million years. *Nature* **538**, 226–228 (2016).

52. David, F. N. *Tables of the Ordinates and Probability Integral of the Distribution of the Correlation Coefficient in Small Samples*. (Cambridge University Press, 1938).

53. Guenther, W. C. Desk Calculation of probabilities for the distribution of the sample correlation coefficient. *Am. Stat.* **31**, 45–48 (1977).

54. Bates, D., Mächler, M., Bolker, B. & Walker, S. Fitting linear mixed-effects models using lme4. *J. Stat. Softw.* **67**, 1–48 (2015).

55. Barton, K. MuMIn: Multi-Model Inference. R package version 1.47.1 (2022).

56. White, G. & Burnham, K. Program MARK: survival estimation from populations of marked animals. *Bird Study* **46**, 120–139 (1999).

57. Nakagawa, S. & Schielzeth, H. A general and simple method for obtaining R2 from generalized linear mixed-effects models. *Methods Ecol. Evol.* **4**, 133–142 (2013).

58. van der Bijl, W. phylopath: Perform Phylogenetic Path Analysis. R package version 1.1.3 (2021).

59. Gonzalez-Voyer, A. & von Hardenberg, A. *An Introduction to Phylogenetic Path Analysis. in Modern Phylogenetic Comparative Methods and Their Application in Evolutionary Biology* (ed. Garamszegi, L.) (Springer Berlin Heidelberg, 2014).

60. Crisp, M. D. *et al.* Phylogenetic biome conservatism on a global scale. *Nature* **458**, 754–756 (2009).

61. Loarie, S. R. *et al.* The velocity of climate change. *Nature* **462**, 1052–1055 (2009).

62. Gilman, S. E., Wethey, D. S. & Helmuth, B. Variation in the sensitivity of organismal body temperature to climate change over local and geographic scales. *Proc. Natl. Acad. Sci. U.S.A.* **103**, 9560–9565 (2006).

63. Şekercioğlu, Ç. H., Primack, R. B. & Wormworth, J. The effects of climate change on tropical birds. *Biol. Cons.* **148**, 1–18 (2012).

64. Borchers, H. W. pracma: Practical Numerical Math Functions. R package version 2.4.2 (2022).

65. Pigot, A. L. *et al.* Macroevolutionary convergence connects morphological form to ecological function in birds. *Nat. Ecol. Evol.* **4**, 230–239 (2020).

66. Sol, D. *et al.* The worldwide impact of urbanisation on avian functional diversity. *Ecol. Lett.* **23**, 962–972 (2020).

67. Sheard, C. *et al.* Ecological drivers of global gradients in avian dispersal inferred from wing morphology. *Nat. Commun.* **11**, 2463 (2020).

68. Burnham, K. & Anderson, D. *Model Selection and Multi-Model Inference: A Practical Information-Theoretic Approach*. (Springer, 2002).

69. Stekhoven, D. J. & Bühlmann, P. MissForest—non-parametric missing value imputation for mixed-type data. *Bioinformatics* **28**, 112–118 (2012).

70. Penone, C. *et al.* Imputation of missing data in life-history trait datasets: which approach performs the best? *Methods Ecol. Evol.* **5**, 961–970 (2014).

71. Carmona, C. P. *et al.* Erosion of global functional diversity across the tree of life. *Sci. Adv.* **7**, eabf2675 (2021).

72. Jetz, W., Thomas, G. H., Joy, J. B., Hartmann, K. & Mooers, A. O. The global diversity of birds in space and time. *Nature* **491**, 444–448 (2012).

73. McGarigal, K., Cushman, S. A. & Stafford, S. *Multivariate Statistics for Wildlife and Ecology Research*. (Springer Science & Business Media, 2013).

74. Noon, B. R. The distribution of an avian guild along a temperate elevational gradient: the importance and expression of competition. *Ecol. Monogr.* **51**, 105–124 (1981).

75. Herring, G., Gawlik, D. E. & Beerens, J. M. Sex determination for the great egret and white ibis. *Waterbirds* **31**, 298–303 (2008).

76. Germain, R. R. *et al.* Changes in the functional diversity of modern bird species over the last million years. *Proc. Natl. Acad. Sci. U.S.A.* In Press, (2023).

77. Rotenberry, J. T. & Balasubramaniam, P. Estimating egg mass–body mass relationships in birds. *Auk* **137**, (2020).

78. Cooney, C. R. *et al.* Ecology and allometry predict the evolution of avian developmental durations. *Nat. Commun.* **11**, 2383 (2020).

79. Jetz, W., Sekercioglu, C. H. & Böhning-Gaese, K. The worldwide variation in avian clutch size across species and space. *PLOS Biol.* **6**, e303 (2008).

80. Werner, J. & Griebeler, E. M. Reproductive biology and its impact on body size: comparative analysis of mammalian, avian and dinosaurian reproduction. *PLOS ONE* **6**, e28442 (2011).

81. Bergmann, C. *Über die Verhältnisse der Wärmeökonomie der Thiere zu ihrer Größe*. (1848).

82. Olson, V. A. *et al.* Global biogeography and ecology of body size in birds. *Ecol. Lett*. **12**, 249–259 (2009).

83. Moreau, R. E. Clutch-size: a comparative study, with special reference to African birds. *Ibis* **86**, 286–347 (1944).

84. Lack, D. The significance of clutch-size. *Ibis* **89**, 302–352 (1947).

85. Cody, M. L. A general theory of clutch size. *Evolution* **20**, 174–184 (1966).

86. Dunn, P. O., Thusius, K. J., Kimber, K. & Winkler, D. W. Geographic and ecological variation in clutch size of tree swallows. *Auk* **117**, 215–221 (2000).

87. Martin, T. E., Auer, S. K., Bassar, R. D., Niklison, A. M. & Lloyd, P. Geographic variation in avian incubation periods and parental influences on embryonic temperature. *Evolution* **61**, 2558–2569 (2007).

88. Ruuskanen, S. *et al.* Geographical variation in egg mass and egg content in a passerine bird. *PLOS ONE* **6**, e25360 (2011).

89. Balasubramaniam, P. & Rotenberry, J. T. Elevation and latitude interact to drive life-history variation in precocial birds: a comparative analysis using galliformes. *J. Anim. Ecol.* **85**, 1528–1539 (2016).

90. Devictor, V. et al. Differences in the climatic debts of birds and butterflies at a continental scale. *Nat. Clim. Change* **2**, 121–124 (2012).

91. Mason, L. R. *et al.* Population responses of bird populations to climate change on two continents vary with species' ecological traits but not with direction of change in climate suitability. *Clim. Change* **157**, 337–354 (2019).

92. Sheridan, J. A. & Bickford, D. Shrinking body size as an ecological response to climate change. *Nat. Clim. Change* **1**, 401–406 (2011).

93. Shultz, S., B. Bradbury, R., L. Evans, K., D. Gregory, R. & M. Blackburn, T. Brain size and resource specialization predict long-term population trends in British birds. *Proc. R. Soc. B* **272**, 2305–2311 (2005).

94. Sayol, F. *et al.* Environmental variation and the evolution of large brains in birds. *Nat. Commun.* **7**, 13971 (2016).

95. Sæther, B.-E. *et al.* Generation time and temporal scaling of bird population dynamics. *Nature* **436**, 99–102 (2005).

96. Sæther, B.-E. *et al.* How life history influences population dynamics in fluctuating environments. *Am. Nat.* **182**, 743–759 (2013).

97. Rosenheim, J. A. & Tabashnik, B. E. Influence of Generation Time on the rate of response to selection. *Am. Nat.* **137**, 527–541 (1991).

98. Owens, I. P. F. & Bennett, P. M. Ecological basis of extinction risk in birds: Habitat loss versus human persecution and introduced predators. *Proc. Natl. Acad. Sci. U.S.A.* **97**, 12144–12148 (2000).

99. Sol, D. *et al.* Unraveling the life history of successful invaders. *Science* **337**, 580–583 (2012).

100. Stevenson, I. R. & Bryant, D. M. Climate change and constraints on breeding. *Nature* **406**, 366–367 (2000).

101. Jàrvinen, A. Global warming and egg size of birds. *Ecography* **17**, 108–110 (1994).

102. Lundblad, C. G. & Conway, C. J. Intraspecific variation in incubation behaviours along a latitudinal gradient is driven by nest microclimate and selection on neonate quality. *Funct. Ecol.* **35**, 1028–1040 (2021).

103. Both, C. *et al.* Avian population consequences of climate change are most severe for long-distance migrants in seasonal habitats. *Proc. R. Soc. B* **277**, 1259–1266 (2010).

104. Hosner, P. A., Tobias, J. A., Braun, E. L. & Kimball, R. T. How do seemingly non-vagile clades accomplish trans-marine dispersal? Trait and dispersal evolution in the landfowl (Aves: Galliformes). *Proc. R. Soc. B* **284**, 20170210 (2017).
